# Comparative Secretome Analysis and Enzyme Cocktail Optimization of Six Fungal Species Under Solid-State and Submerged Fermentation for Lignocellulosic Saccharification of Flax Shives

**DOI:** 10.64898/2026.06.03.729743

**Authors:** Nastassia Kaugarenia, Barbara Deracinois, Quentin Haguet, Svetlana Heyte, Renato Froidevaux, Vincent Phalip, Egon Heuson

## Abstract

Lignocellulosic biomass represents a promising renewable feedstock for sustainable biorefinery applications, yet efficient enzymatic saccharification remains challenging due to the recalcitrant structure of plant cell walls. This study presents a comprehensive comparative analysis of enzymatic activities, saccharification performance, and secretome composition of six fungal species cultivated under solid-state fermentation (SSF) and submerged fermentation (SmF) conditions using untreated flax shives as substrate. While SmF yielded approximately 4-fold higher total protein concentrations (0.38 ± 0.13 g.L^-1^ *vs.* 0.08 ± 0.02 g.L^-1^), SSF-derived enzymes demonstrated superior specific enzymatic activities, particularly for endo-xylanase and endo-cellulase, resulting in more efficient biomass saccharification. Proteomics analysis revealed distinct secretome profiles between fermentation modes, with SSF showing higher proportions of polysaccharide metabolism proteins (71.0%) compared to SmF (49.3%), while SmF exhibited greater enzyme diversity including more lytic polysaccharide monooxygenases (LPMOs) and auxiliary activity enzymes. *Trichoderma* species consistently demonstrated the highest saccharification efficiency, with glucose yields reaching 2.37 mM under SSF conditions. A Scheffé simplex-lattice mixture design comprising 65 enzyme cocktail combinations revealed significant synergistic interactions between several cocktails, with the binary mixture of *Trichoderma* 2SA21 and *P. chrysogenum* achieving 54% synergy - in terms of higher sugar release above expectations - and the highest total monosaccharide release (1.80 mM). These findings provide practical guidance for developing cost-effective enzyme cocktails for lignocellulosic biorefinery applications, emphasizing the importance of fermentation mode selection and strategic strain combination over enzyme supplementation complexity. The methodology established here, combining systematic screening, comparative proteomics, and statistical mixture design, offers a robust framework for optimizing fungal enzyme systems across diverse biomass substrates.

**BULLET POINTS:** Superior enzymatic activity (xylanase, cellulase) and saccharification in solid-state fermentation Superior total protein content and diversity in submerged fermentation Specific enzyme cocktails combination can lead to synergistic effects, justifying a combinatorial approach

**GRAPHICAL ABSTRACT:** 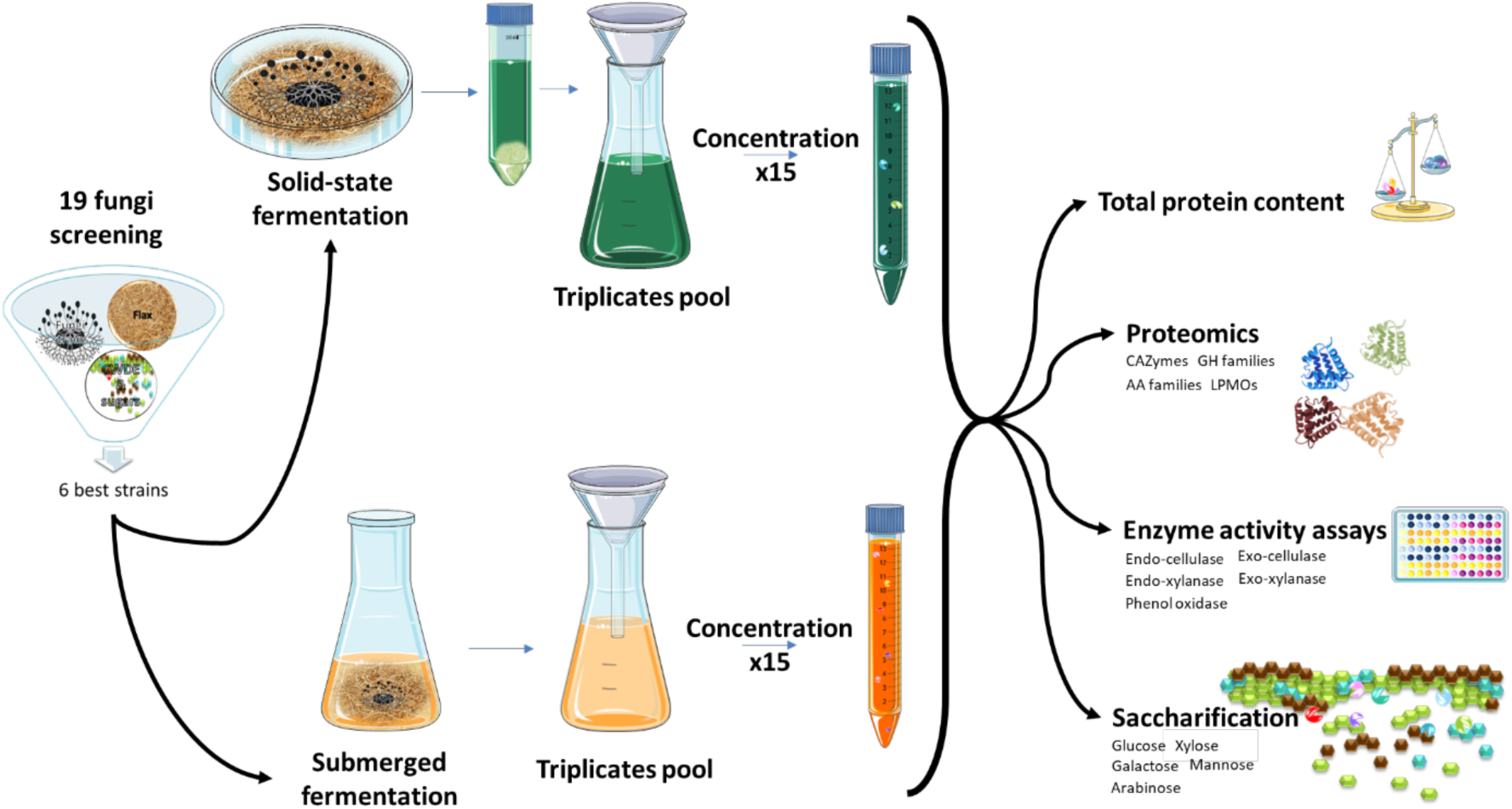

## 1. Introduction

Saccharification, defined as the hydrolysis of carbohydrate-containing polymeric materials into monomeric carbohydrates or units with low degrees of polymerization (Moron *et al*., 2025), represents a critical step in lignocellulosic biomass valorization. This process enables the conversion of biomass into biofuels, saccharides, and other oxygen-containing chemicals, making it an essential component of the sustainable circular economy (Ashokkumar *et al*., 2022; Deng *et al*., 2023; Mujtaba *et al*., 2023; Srivastava *et al*., 2018). The compact structure of lignocellulose, characterized by complex bonding among cellulose, hemicellulose, and lignin, presents a significant challenge for biomass degradation. However, plant cell wall-degrading microorganisms have evolved specialized machinery tailored for this task, with fungi being the most efficient organisms in this biomass recycling ecosystem (Janusz *et al*., 2017; Srivastava *et al*., 2018). Fungi possess a unique extracellular enzyme system comprising two main categories: hydrolytic enzymes that degrade various polysaccharides, and oxidative enzymes that depolymerize lignin and open rings from organic compounds (Kumar and Chandra, 2020). Based on their degradation patterns and substrate preferences, these fungi are classified into three main categories: soft-rot fungi, white-rot fungi and brown-rot fungi. Soft-rot fungi are predominantly ascomycetes that degrade polysaccharides in the surface layers of plants, particularly in wet environments (Goodell and Jellison, 2008). Representative species include *Aspergillus niger*, *Trichoderma reesei*, and *Penicillium chrysogenum*. White-rot fungi, such as *Trametes versicolor*, are basidiomycetes capable of degrading all three major lignocellulose components: cellulose, hemicellulose, and lignin (Andlar *et al*., 2018). Brown-rot fungi are also basidiomycetes but differ in their degradation profile, efficiently metabolizing cellulose and hemicellulose while only slightly modifying lignin (Arantes and Goodell, 2014; Sánchez, 2009).

Over the past decades, plant cell wall-degrading filamentous fungi have been extensively studied for producing enzyme cocktails with various catalytic activities applicable to lignocellulosic feedstock hydrolysis. Numerous commercial enzyme cocktails, like Cellic® CTec2, Celluclast® 1.5L Viscozyme L® (Berlin *et al*., 2007; Dąbkowska *et al*., 2017; Gabiatti Junior *et al*., 2020) are produced from fungi (*Trichoderma reesei*, *Aspergillus* sp.) due to their natural predisposition to degrade recalcitrant plant cell walls.

The fermentation process critically influences microbial growth, enzyme production, and overall productivity. Two types of fermentation are widely employed for lignocellulolytic enzyme production: solid-state fermentation (SSF) and submerged fermentation (SmF) (Ahmed and Bibi, 2018). Industrial enzyme production strategies predominantly utilize submerged fermentation, where fungi are fully suspended in high volumes of medium. This well-established approach is favored for industrial-scale enzyme production due to the availability of established methods for monitoring and controlling critical parameters including culture homogeneity, pH, temperature, oxygenation, tank scale-up, and downstream process compatibility (Reihani and Khosravi-Darani, 2018; Singhania *et al*., 2010).

However, SSF has gained increasing attention for its advantages when cultivating filamentous fungi. In this implementation, fungi grow on and within solid humidified particles of plant substrate and secrete enzymes to invade and digest them, closely mimicking their natural habitat. This alignment between growth conditions and natural environment, without free water, usually leads to superior microbe-substrate interactions, usually resulting in higher enzyme titers (Kalim and Ali, 2016). Indeed, cellulolytic filamentous fungi penetrate cellulosic substrates through hyphal extensions, often presenting their cellulase systems in confined cavities within cellulosic particles. Importantly, aerial hyphae - which occur exclusively in SSF - are primarily responsible for oxygen uptake in fungal cultures (Lynd *et al*., 2002; te Biesebeke *et al*., 2002). Enzymes produced via SSF exhibit remarkable stability toward temperature and pH variations (Dutta *et al*., 2008). Furthermore, enzymes produced using the same biomass substrate intended for bioconversion demonstrate higher efficiency compared to those produced on alternative cellulosic substrates (Siqueira *et al*., 2020). Beyond biological advantages, SSF offers significant economic benefits including simpler bioreactor technology (stacked trays), reduced downstream processing requirements, lower energy consumption, and cost-effective media (Tišma *et al*., 2021; Webb and Manan, 2017). These advantages have driven increased focus on SSF development and efforts to overcome its challenges, particularly parameter control during scale-up processes (Rodriguez-Couto and Sanromán, 2006).

For instance, Coradi *et al*., (2013) demonstrated superior lipase activity from *T. harzianum* in SSF mode. Conversely, Teixeira da Silva *et al*., (2016) reported that SmF with the thermophilic fungus *Myceliophthora heterothallica* grown on wheat bran and sugarcane bagasse produced higher endoglucanase and β-glucosidase activities, though pH and temperature stability of the enzymes showed minimal differences between modes. However, for the white-rot fungus *Pleurotus ostreatus* grown on cottonseed hull, corncob, or poplar wood, An e*t al.,* (2016) observed three- to four-fold higher laccase, xylanase, and endoglucanase activities in SmF compared to SSF. These contrasting results highlight the complex interaction between fungal species, substrate type and fermentation mode.

Cunha *et al.,* (2012) introduced sequential solid-state fermentation (SF), supposedly combining the advantages of both approaches. This strategy employs SSF for enzyme induction during the initial days of incubation (enabling physical contact between fungus and substrate) followed by SmF for the remainder of incubation. Using *Aspergillus niger* on sugarcane bagasse in both 500 mL Erlenmeyer flasks and 5 L bubble column bioreactors, they achieved notable results: in Erlenmeyer flasks, SF yielded only 20% higher endoglucanase activity compared to SSF alone. However, in bubble column bioreactors, SF demonstrated three-fold greater endoglucanase activity. Despite the shortened incubation time of 30 hours (compared to the typical several days), SF induced earlier endoglucanase and xylanase activity, though activity levels decreased by half over time. In contrast, SmF showed increasing enzyme activities over time, correlating with carbon source depletion. Interestingly, reducing sugar release peaked within the first 6 hours in both conditions, decreasing to near zero after 12 hours, indicating insufficient enzyme production for efficient saccharification (Cunha *et al*., 2012).

More recently, Arif *et al.,* (2024) compared enzymatic activities across three fermentation modes (SSF, SmF and SF) using four fungal strains - *Penicillium crustosum*, *Fusarium nygamai*, *Trichoderma capillare*, and *Aspergillus calidoustus* - on olive stone grounds, a notably recalcitrant substrate. *Penicillium crustosum* was unique in producing exo-glucanase activity regardless of fermentation mode, achieving optimal cellulase activity (both endo- and exo-) in SF mode. SSF and SmF modes were approximately half as efficient with similar profiles, except for laccase activity, which was present only in SSF and SF modes. *Fusarium nygamai* exhibited an unusual profile, with increasing total protein content in both SSF and SF modes (up to 600 μg.mL^-1^ by day 9), yet nearly no cellulase activity and only remarkable laccase activity. This pattern suggested a lignin degradation orientation. Interestingly, *Trichoderma capillare* displayed a strikingly similar profile, which is equally atypical for this species, otherwise well known for its high cellulase production (Florencio *et al*., 2015). Here, no cellulase activity was detected; only laccase activity was observed in SSF and SF modes. While laccase activities were comparable (SF showing ∼25% higher activity), SF protein content was four-fold higher than SSF. *Aspergillus calidoustus* achieved the highest endoglucanase activity in SF mode, followed by SSF and then SmF. Protein content followed the same general trend, with SF showing 4-5 times more protein on days 2-3 than SSF and SmF. Subsequently, SF protein content decreased while SSF content increased, eventually reaching equivalent levels. Laccase activity was highest in SSF mode and absent in SmF. Overall, this study demonstrated SF proved to be the most productive fermentation mode for cellulase production in *A. calidoustus* and *P. crustosum*, with high protein content, followed by SSF. SmF performed poorest for enzyme activities and protein content across all fungi, with almost no laccase activity. This pattern was attributed to mycelial pellet formation in SmF: fungi form compact mycelial pelleted in liquid culture, whereas in SSF and SF (during induction), they developed dispersed filamentous mycelia more favorable for enzyme production.

Sankar *et al*., (2023) extended the comparison beyond enzyme activities to include saccharification performance and proteomics analysis. Working with *Penicillium janthinellum* grown on wheat bran supplemented with cellulose, they evaluated enzyme performance on pretreated rice straw. SSF-derived enzymes outperformed their SmF counterparts, releasing higher glucose and xylose levels with lower cellobiose accumulation. Proteomics analysis revealed no difference in protein types between SSF and SmF (221 total proteins identified), but substantial differences in expression levels. At day 5, SSF showed 9- to 390-fold higher β-glucosidase expression and 14- to 63-fold higher monooxygenase expression compared to SmF. These differences diminished when fermentation extended to 9 days.

Somadder *et al*., (2025) compared *Aspergillus awamori* and *A. oryzae* (tested separately) grown on sorghum milled grain or bran. SSF for both strains produced up to two-fold higher total reducing sugars than SmF. However, for *A. awamori*, SmF performance matched SSF when grown on sorghum bran, highlighting the influence of carbon source-fermentation mode interactions previously observed by An *et al*., (2016). Glucoamylase, α-amylase, and protease activities were also higher in SSF mode. While direct total protein quantification was not performed, precluding determination of whether higher activities resulted from increased enzyme production or enhanced specific activity. Particle size analysis revealed that the 0.6-1.18 mm range was optimal for sugar production and enzyme activities (maximizing surface area for enzyme access), with particles >1.18 mm ranking second. Also, during SSF, pH and moisture content remained stable (pH decrease of 1.26 units from initial, 3.41% water loss).

Leadbeater *et al.,* (2025) conducted an extensive study with *Parascedosporium putredinis*, a soft-rot ascomycete, on wheat straw. They reported six-fold higher soluble protein production in SSF by day 14, though dry mass differed substantially (7.5 g for SmF *vs.* 175 g for SSF). Unlike in Sankar *et al.,* (2023), SSF exhibited lower CAZyme diversity but higher abundance of specific families (GH1, GH3, GH6, GH7). GH7 (cellobiohydrolase) became the dominant family, reaching 54% of total CAZymes in SSF by day 21. Whether these diversity changes reflect fungus-specific characteristics or methodological influences remains unclear, though previous work has demonstrated that this fungus alter secretome expression depending on substrate composition (Scott *et al*., 2024). SmF displayed volatile protein expression patterns characterized by rapid initial enzyme secretion followed by decline and a higher proportion of Auxiliary Activity (AA) enzymes. Despite this, SSF achieved 221-fold greater substrate conversion efficiency at day 14. Significant differences emerged in carbohydrate-binding domain (CBM) profiles between soluble and insoluble secretomes. Notably, SmF showed abundant CBM20, CBM18, and CBM87 (associated with starch, chitin, and galactosaminogalactan binding, respectively), which were nearly absent in SSF. They conclude that this pattern suggested higher starvation stress in SmF, evidenced by autolysis and self-degradation of fungal cell walls to support nutrient recycling and mobilization of internal starch reserves. Conversely, both fermentation modes exhibited high abundances of CBM1, CBM35, and CBM91, which target cellulose, xylans, and mannans, respectively. This importance was also highlighted by (Liu *et al*., 2020) where higher proportion of CBM-containing enzymes was induced in SmF (CMB1, CMB20), as CMB enhance substrate binding at the beginning of hydrolysis by increasing enzyme-substrate collision probability. When normalized for total protein content, saccharification performance revealed distinct temporal patterns: SmF peaked earlier (day 7) with rapid enzyme secretion, while SSF peaked later (day 14) but maintained high activity through day 21. Despite these different enzyme profiles and temporal dynamics, both modes achieved comparable peak saccharification activities.

While these studies provide valuable insights into SSF versus SmF performance, significant limitations constrain our understanding of optimal fermentation strategies. Previous research has typically examined only a few species at a time, with studies either monitoring enzymatic activity without assessing saccharification efficiency, or conducting comprehensive proteomic analyses without measuring enzymatic activity. These constraints limit our understanding of inter-species variation in fermentation mode response, prevent direct comparative assessment of fungal strain efficiency across identical conditions, and restrict identification of optimal fungus-fermentation combinations. Furthermore, if we focus on the substrate specificity aspect now, a given fungus often exhibit different enzyme secretion depending on the lignocellulosic carbon source leading to variable product efficiency as we previously demonstrated (Raulo *et al*., 2021; Alananbeh *et al*., 2024; Irfan *et al*., 2014), creating a practical challenge for industrial applications. Biorefinery operators have to work with locally available agricultural residues or industrial byproducts, making it essential to identify which fungal strains perform best on specific substrates (Brown *et al*., 2024). Consequently, selecting the optimal fungus-substrate pairing remains largely empirical (Arif *et al*., 2024; Coradi *et al*., 2013; Debeire *et al*., 2014; Sohail *et al*., 2016; Teixeira da Silva *et al*., 2016).

The present study addresses these knowledge gaps through a systematic three-phase investigation designed to comprehensively evaluate fungal performance across fermentation modes and identify optimal enzyme production strategies. Firstly, 19 fungal strains (9 different genera) were compared on a selected lignocellulosic carbon source (flax shives) under SSF conditions. Performances were evaluated based on three complementary metrics: total protein content, enzymatic activity profiles (cellulases, hemicellulases, and auxiliary enzymes), and saccharification efficiency. This comprehensive screening enables identification of the highest-performing strains and reveal patterns linking fungal taxonomy, enzyme production, and substrate conversion efficiency. Six of them, representing diverse performance profiles, were selected for further comparative analysis under both SSF and SmF conditions based on comprehensive proteomics analysis to characterize secretome composition. In the final phase, it was evaluated whether combining enzymes from different fungal sources can enhance saccharification beyond what individual strains achieve. These enzyme cocktails were formulated by mixing secretomes from the six strains in various ratios using a dedicated design of experiment to cover the complete combination space.

## 2. Materiel and Methods

### 2.1. Biomass

Dry flax shives (untreated) were provided by a local farm (Hauts-de-France, France). The flax was further ground with a batch grinder (BLET Measurement Group, Saint-Germain-en-Laye, France), then meshed with shifter Type 3D (Bioblock Scietific Retsch GmbH, Ile-de-France, France) and the fraction between 200 µm and 1 cm was kept. Mild sterilization was performed at 115°C for 15 minutes.

### 2.2. Fungi

*Fusarium* sp. 2DA63, *Penicillium* sp. 2DA2, *Trichoderma* sp. 2DA61, *Trichoderma* sp. 2DA62, *Trichoderma* sp. 2DA67, *Trichoderma* sp. 2SA21, *Paecilomyces* sp. 11SA2, *Paecilomyces* sp. 11SE, *Talaromyces* sp. 47, were isolated from used coffee grounds provided by Gecco (Avelin, France) (Beaudor, 2023). *Trichoderma harzianum* MUCL 29707, *Trametes versicolor* MUCL 1011*, Fusarium graminearum* MUCL 11946*, Mucor circinelloides* MUCL 15438, were purchased at Belgian Co-ordinated Collection of Microorganisms (BCCM). *Aspergillus terreus* 826 was from DSMZ collection (Leibnitz Institute, Germany). *Aspergillus oryzae* UMIP 1042.42 was from Institut Pasteur collection (France)*. Penicillium chrysogenum, Coriolopsis polyzona, Mucor* sp. and *Mucor fragilis,* were isolated in the laboratory from a decaying tree trunk in Lille (France).

Fresh conidia collection was obtained by cultivation of provided strains on potato dextrose agar (PDA) medium (39 g.L^-1^, 20 mL into 90 mm Petri dishes) for 7 days at 22 ± 1 °C. Conidia were harvested by rubbing the mycelium within physiological buffer (0.85% NaCl, 0.01% Tween 80, sterilized) then the suspension was filtered through sterilized Miracloth filter (Millipore, Merck Group, Darmstadt, Germany). Stock aliquots were made with 0.5 mL of filtrated conidia and 0.5 mL of glycerol 60% and kept at -80 °C until use. The conidia count was realized with optical microscope and Thoma counting chamber (Paul Marienfeld GmbH, Lauda-Königshofen, Deutchland).

### 2.3. Commercial enzymes cocktails

Commercial enzymes were purchased from Sigma-Aldrich (St. Louis, MO, USA): Xylanase (Pentopan Mono BG®) 2500 U/g (X2753), Hemicellulase from *Aspergillus niger* 0.3-3.0 U/mg (H2125), Cellulase from *Trichoderma reseei* (Celluclast®) 1.5L 770-910 EGU/mL (C2730), Cellulase Enzyme Blend (Cellic® CTec2) 1000 – 1300 U/mL (SAE0020), Viscozyme® L 110-130 FBGU/mL (V2010) and Pectinase from *Aspergillus niger* (Pectinex^®^ Ultra SPL) 3800 U/mL (P2611).

### 2.4. Chemicals

All chemicals were purchased at Sigma-Aldrich. Methanol was LC/MS grade. Chloroform was analytical grade 99%.

### 2.5. Enzyme’s production

#### 2.5.1. Solid State Fermentation (SSF) of 19 fungi

SSF was performed with 2.00 ± 0.05 g of previously described flax in 90 mm Petri dishes moisturized with 16 mL of sterilized M3 medium (Mitchell *et al*., 1997) (pH 6.9). All substrates were inoculated with fungi at 2.5×10^5^ conidia/g_flax_. A negative control consisted of the substrate moisturized with M3 medium without fungus. Fungi were incubated at 22 ± 1 °C for 7 days. All cultures were conducted in biological triplicates.

#### 2.5.2. Submerged Fermentation (SmF) of the six selected fungi

SmF was performed in 250 mL flasks with 2.00 ± 0.05 g of previously described flax and 80 mL of M3 medium. Flasks were sealed with sterile AeraSeal^TM^ films (Dutscher SAS, Bernolsheim, France). All substrates were inoculated with fungi at 2.5×10^5^ conidia/g_flax_. A negative control consisted of the substrate and 80 mL of M3 medium without fungus. Fungi were incubated at 22 ± 1 °C and 60 rpm for 7 days. All cultures were conducted in biological triplicates.

### 2.6. Soluble proteins extraction

Soluble proteins from SmF were recovered by filtration with Miracloth filter. Proteins from SSF were extracted with 80 mL sodium acetate buffer (50 mM, pH 5.0) at 30 °C, 200 rpm during 1 h, then the supernatant was recovered from substrate and mycelium by filtration with Miracloth. All biological triplicates were pooled before concentration step with Vivaspin® 20 PES 10 kDa ultrafiltration membrane (Sartorius, Göttingen, Germany). The concentration was performed until 14 mL were reached and this enzyme extract was filtrated (0.22 µm PVDF sterile) then kept at -80 °C until use.

### 2.7. Total protein quantification

Total protein content was quantified using the Bradford assay against a standard Bovin Serum Albumin (BSA) curve, with culture media blank (SmF) or extraction buffer blank (SSF): to 5 µL of supernatants were added to 195 µL of Bradford Reagent (Sigma). Absorbances were read at 595 nm after 10 min of reaction. Commercial enzyme cocktails in liquid form were measured for total protein content after at least a 10-fold dilution in acetate buffer.

### 2.8. Enzymatic assay analysis

Xylanase and cellulase activities were measured according to (Raulo *et al*., 2021): AZO-xylan (Birchwood) and AZO-CM-cellulose (Megazyme, Ireland) were used to assay endo-xylanases (endo-1,4-β-Xylanase) and endo-cellulases (endo-1,4-β-Glucanase) activities while two pNP-linked substrates 4-Nitrophenyl-β-D-xylopyranoside and 4-Nitrophenyl-β-D-glucopyranoside (Sigma Aldrich, UK) were used to measure exo-xylanases (or β-xylosidase, exo-1,4-β-Xylosidase) and exo-cellulases (or β-glucosidase, exo-1,4-β-Glucosidase) activities. In a 96 U-well plate, 30 µL of the concentrated enzyme extract (*cf.* 2.6) were assayed with 150 µL of substrate 0.5% (w/v), 75 µL of 100 mM sodium acetate pH 5.0 and 45 µL H_2_O in a total volume of 300 µL. The reaction mixtures were incubated at 25 °C for 2 h with 5 min of initial shaking at 600 rpm. The reactions were then stopped by adding 50 µL of the reaction mixture to 200 µL of ethanol (95% v/v) (AZO-dyed substrates) or trisodium phosphate pH 12 (2% w/v) (pNP substrates). The remaining substrates were precipitated by centrifugation at 2,120 *g* for 10 min. 100 µL of the supernatants were transferred to a flat-bottom 96-well reading plate and the optical densities were measured at 595 nm (ε_595_ = 5,724 M^-1^.cm^-1^) and 405 nm (ε_405_ = 16,789 M^-1^.cm^-1^), AZO-substrates and pNP-substrates, respectively, with a plate reader (FilterMax F5, Molecular Devices, USA). 1 U is defined as 1 µmol of product (soluble AZO dye or pNP) released per minute. The extinction coefficients were calculated by preparing Remazol Brilliant Blue R and p-Nitrophenol in the same conditions as the sample.

PPOs activities were determined according to the method described by Winder & Harris (1991) with modifications. Briefly, 50 μL of enzymatic concentrated extract were mixed with 50 μL of 350 mM sodium phosphate monobasic buffer (pH 7.2) containing DMF 20% (v/v) and 50 μL of 24 mM aqueous solution of 3-Methyl-2-benzothiazolinone hydrazone hydrochloride monohydrate (MBTH). Reaction was initiated by addition of 50 μL of substrate (20 mM dopamine aqueous solution). The plate was stirred 3 s before measurement. Absorbance variations were recorded at 485 nm over 30 min at 30 °C. One unit of PPOs activity was defined as the formation of 1 μmol.min^-1^ (U) of pink pigment obtained from Michael addition between MBTH and the quinone produced by phenolic substrate oxidation (ɛ_505_ nm = 29,000 M^−1^.cm^−1^, pH 6.9) (Winder and Harris, 1991).

### 2.9. Fungi saccharification

2.9.1. *Standard procedure*

The saccharification assay was performed in 2 ml Eppendorf tube volume with 1 mL of enzymatic concentrated extract with 15.4 ± 0.3 mg of flax and an addition of 25 µL sodium acetate buffer 2.05 M pH 5.0. Reactions were incubated for 48 h at 30 °C with 200 rpm shaking. Enzymes were inactivated by heating at 100 °C for 5 min followed by centrifugation for 10 min at 11,000 *g* to pellet the solids. The supernatant was filtrated (0.22 µm PTFE) then the monosaccharides were quantified with HPLC-RID.

#### 2.9.2. DOE mixture procedure

After normalization of enzyme extracts to a total protein content of 0.04 g.L^-1^, they were combined in a total volume of 0.7 mL according to a Scheffé simplex-lattice mixture design, in order to evaluate synergistic interactions among six fungal strains (Cornell, 2002). The experimental design comprised 65 combinations distributed as follows:

- 6 pure strains (Mix 1-6)
- 15 binary mixtures at 50% proportions (Mix 7-21)
- 20 ternary mixtures at 33% proportions (Mix 22-41)
- 15 quaternary mixtures at 25% proportions (Mix 42-56)
- 6 quinary mixtures at 20% proportions (Mix 57-62)
- 3 replicates of the complete hexanary mixture at 16.67% proportions (Mix 63-65)

Saccharification protocol was applied with following modification due to concentrated enzyme extract availability: the saccharification assay was performed in 2 ml Eppendorf tube volume with 0.7 mL enzymatic mix extract with 10.5 ± 0.3 mg of flax and an addition of 17.5 µL sodium acetate buffer 2.05 M pH 5.0. Before saccharification, mixed extracts were filtered through a 96-well filter plate MultiScreen HTS-HV, 0.45 µm Durapore® (Merck Millipore, Cork, Ireland), previously sterilized under UV light. Flax extract without fungus was used as negative control. The mixing procedure, filtration and saccharification were performed using the Biomek i7 pipetting platform (Beckman Coulter, Brea, USA). Detailed steps of the program are described in Supplementary Data 3.

#### 2.9.3. Saccharification with commercial enzyme cocktails

The same standard procedure was used except that commercial enzyme cocktails were prepared to a final volume of 1 mL in 50 mM acetate buffer (pH 5.0) at the following activity levels based on the supplier’s theoretical activity: Xylanase 10 U.mL^-1^; Hemicellulase 10 U.mL^-1^; CTR 7 EGU.mL^-1^; CEB 10 U.mL^-1^; Viscozyme 10 FBGU.mL^-1^; Pectinase 76 U.mL^-1^.

### 2.10. Sugars analysis and quantification

The saccharification samples qualitative and quantitative analysis were performed on an HPLC (Shimadzu, Japan) equipped with pump LC-30AD SP, Autosampler SIL-20AC HT, column oven CTO-20A coupled with Refractive Index Detector RID-20A, using a Phenomonex Rezex^TM^ RPM-Monosaccharrides Pb^2+^ (300 x 7.8 mm) column with a Phenomonex Rezex^TM^ RPM-Monosaccharrides Pb^2+^ (50 x 7.8 mm) guard column. Injection volumes of 30 µL were used for all the samples. The mobile phase consisted of MilliQ water. Elution was carried out at 0.8 mL.min^-1^, with a 75 °C oven temperature for the column and 50 °C for the detection cell. Cellobiose, glucose, xylose, galactose, arabinose and mannose were used as standards and were analyzed in the same conditions as samples.

### 2.11. Secretome analysis

#### 2.11.1. Protein precipitation

One mL of previously obtained enzyme extract (*cf.* 2.6) was desalted with Vivaspin® 20 three times with 20 mL of MilliQ water. The last concentration set the sample at 400 µL, then the sample was split into 2 x 200 µL. All duplicates (200 µL) were extracted with methanol/chloroform/water (800/200/600 µL), followed by centrifugation (10,000 *g*, 2 min) and removal of the upper phase. The lower phase together with the interphase protein pellet was washed with 800 µL methanol, centrifuged, and dried at room temperature after removal of the upper phase (Wessel and Flügge, 1984). Bovine serum albumin (BSA, 1 mg.mL^-1^) was used as an experimental control. One replicate was used for SDS-PAGE (sodium dodecyl sulfate-polyacrylamide gel electrophoresis) analysis (*cf.* 2.11.2), and the other for proteomics analysis (*cf.* 2.11.3).

#### 2.11.2. SDS–PAGE analysis

Protein content was estimated by SDS-PAGE using the Laemmli method (Laemmli, 1970). Briefly, precipitated samples were resuspended in 25 µL (or 200 µL for BSA) of Laemmli buffer (final composition: 1× buffer supplemented with 5% β-mercaptoethanol), heated for 10 min at 95 °C, and centrifuged (10,000 *g*, 5 min). A volume of 20 µL, as well as 5 µL of a solution of molecular weight markers (Bio-Rad), was loaded onto Any kD™ Mini-PROTEAN® TGX Stain-Free™ gels (Bio-Rad, Marnes-la-Coquette, France). The migration was performed for one hour in a base buffer containing Tris (30.3 g.L^-1^), glycine (144 g.L^-1^), and SDS (10 g.L^-1^) at pH 8.3. The electrophoretic migration was performed under 120 V during 40-50 min. Following electrophoresis, proteins were visualized using Stain-Free imaging technology.

#### 2.11.3. Proteomics analysis

Precipitated samples were resuspended in 21 µL of water and 30 µL of 50 mM ammonium bicarbonate. Reduction was performed by adding 3 µL of 100 mM dithiothreitol (DTT) and incubating for 10 min at 80 °C, followed by alkylation with 6 µL of 100 mM iodoacetamide (IAA) for 20 min at room temperature in the dark. Trypsin-based hydrolysis was carried out after an addition of a trypsin/Lys-C (Promega, Madison, WI, USA) mixture at 3 ng.µL^-1^ followed by an incubation for 3 h at 37 °C and a second addition of the same enzyme quantity was performed followed by an incubation for 16 h at 37 °C and finally centrifuged for 10 min at 8,000 *g*. Pellets were discarded and the supernatants were directly submitted to peptidomics analysis.

The peptides generated by trypsinolysis were centrifuged for 5 min at 10,000 *g* before analysis by reverse-phase high-performance liquid chromatography coupled to tandem mass spectrometry (RP-HPLC-MS/MS). Separation was carried out using a C18 column (Uptisphere CS Evolution, 250 × 3.0 mm, 2.6 µm, Interchim, France) on a Dionex Ultimate™ 3000 UPLC system (Thermo Fisher Scientific) operated by Chromeleon Xpress software. The system was coupled to a maXis Impact II ESI-QTof mass spectrometer (Bruker Daltonics GmbH, Champs-sur-Marne, France) controlled by Compass OtofControl software (v. 4.1.3.4, Bruker). The coupling of HPLC and MS/MS was managed using Compass HyStar software (v. 4.1.21.2, Bruker). Ten microliters of each protein hydrolysate were injected, and peptide elution was carried out at 30 °C and a flow rate of 0.5 mL.min⁻¹ using the following linear gradient of solvent A (H₂O with 0.1% (v/v) FA) and solvent B (acetonitrile (ACN) with 0.1% (v/v) FA): 5% solvent B to 30% solvent B for 40 min, then 30% to 100% solvent B for 10 min, 100% solvent B for 5 min, and back to 0% solvent B over 5 min. MS and MS/MS analyses were performed in positive ion mode using data-dependent acquisition (DDA) over a mass-to-charge (m/z) range of 50–1,800. Capillary voltage was set at 3.5 kV, nebulizer gas pressure at 2.8 bar, desolvation temperature at 180 °C, and drying gas flow rate at 10 L.min⁻¹. MS/MS spectra were acquired for the top 10 most intense precursor ions with a minimum intensity threshold of 2,000 counts. Fragmentation was performed by collision-induced dissociation (CID), with a collision energy ramp from 23 to 65 eV depending on m/z. Prior to analysis, the system was calibrated using a sodium formate solution prepared as follows: 0.5 volume of 1 M NaOH, 0.025 volume of FA, and 50 volumes of 50/50:isopropanol/H₂O. All mass spectra were acquired and analyzed using Compass DataAnalysis software (v. 4.4, Bruker). The RP-HPLC-MS/MS analysis was carried out in triplicate.

Mass spectrometry data processing and the protein database search were performed via Peaks Studio X+ software (Bioinformatics Solutions, Waterloo, ON, Canada) with different databases downloaded from UniProtKB : Swiss-Prot (571,587 entries on May 29, 2024), *Linium usitatissimum* proteome (943 entries on June 27, 2025), *Basidiomycete* proteome (162,908 entries on June 27, 2025), *Aspergillus terreus* proteome (26.686 entries on June 27, 2025), *Trichoderma* proteome (248,718 entries on June 27, 2025), *Trichoderma harzianum* proteome (37.883 entries on June 27, 2025), *Fusarium* proteome (1,547,694 entries on June 27, 2025) and *Penicillium chrysogenum* proteome (24,758 entries on June 27, 2025). Database searches were performed specifying trypsin as the protease, and three missed cleavage sites allowed. Precursor and fragment ion mass tolerances were set to 10 ppm and 0.1 Da, respectively. Variable methionine oxidations and cysteine carbamidomethylation were specified and post-translational modifications (PTMs) were searched against more than 650 modifications available in the PEAKS® database, with a maximum of 3 variable modifications per peptide. Peptide sequences identified by the Peaks Studio X+ were filtered with a false discovery rate (FDR) strictly lower than 0.1% and a minimum of two unique peptides per protein was required for high-confidence protein identification.

## 3. Results and Discussion

### 3.1. Screening of 19 fungi with SSF: Enzymatic specific activity and saccharides release

To start our study, we selected19 different fungal strains, which were then evaluated for their ability to produce saccharification enzymes and release sugar from our selected biomass. All strains were assessed under solid-state fermentation (SSF) conditions to identify those producing the most efficient enzymatic systems for free sugar generation, as this mode is described to be the most proficient for all soft-rot and white-rot fungi (Andlar *et al*., 2018). Resulting enzyme activities (including endo-xylanase, endo-cellulase, exo-xylosidase, exo-glucosidase and phenol oxidase activity) are reported in **Fig. 1** and the corresponding saccharide profiles, cellobiose and glucose from cellulose, xylose and galactose from hemicellulose, are quantified in **Fig. 2**. Overall, these results demonstrated a high diversity of both enzymatic and saccharification profiles and the identified profiles are detailed in the next sub-chapters.

**Fig. 1:**
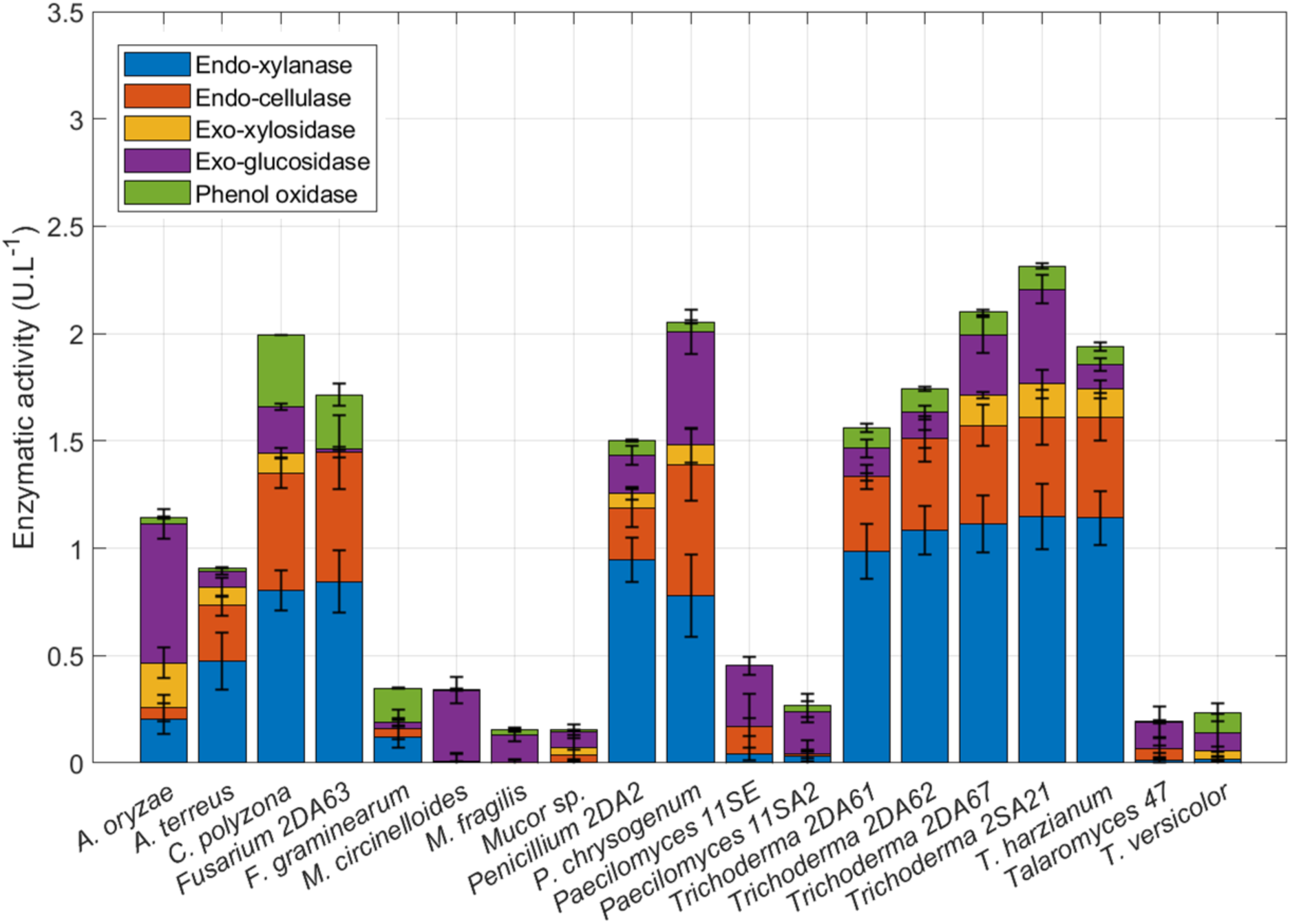
Comparative analysis of lignocellulolytic enzyme activities across fungal species in SSF with 2 g of flax shives (16 mL M3 medium, 22 °C for 7 days, pooled biological triplicates). Stacked bar chart showing the enzymatic activity (U.L^-1^) of five key lignocellulose-degrading enzymes: endo-xylanase (blue), endo-cellulase (orange), β-xylosidase (yellow), β-glucosidase (purple), and phenol oxidase (green) produced by 19 different fungal strains. Endo activities were measured with AZO substrates. Exo activities were measured with pNP substrates. Phenol oxidase activity was measured with MBTH-dopamine assay. Error bars indicate standard deviation (n = 3 replicates).

**Fig. 2:**
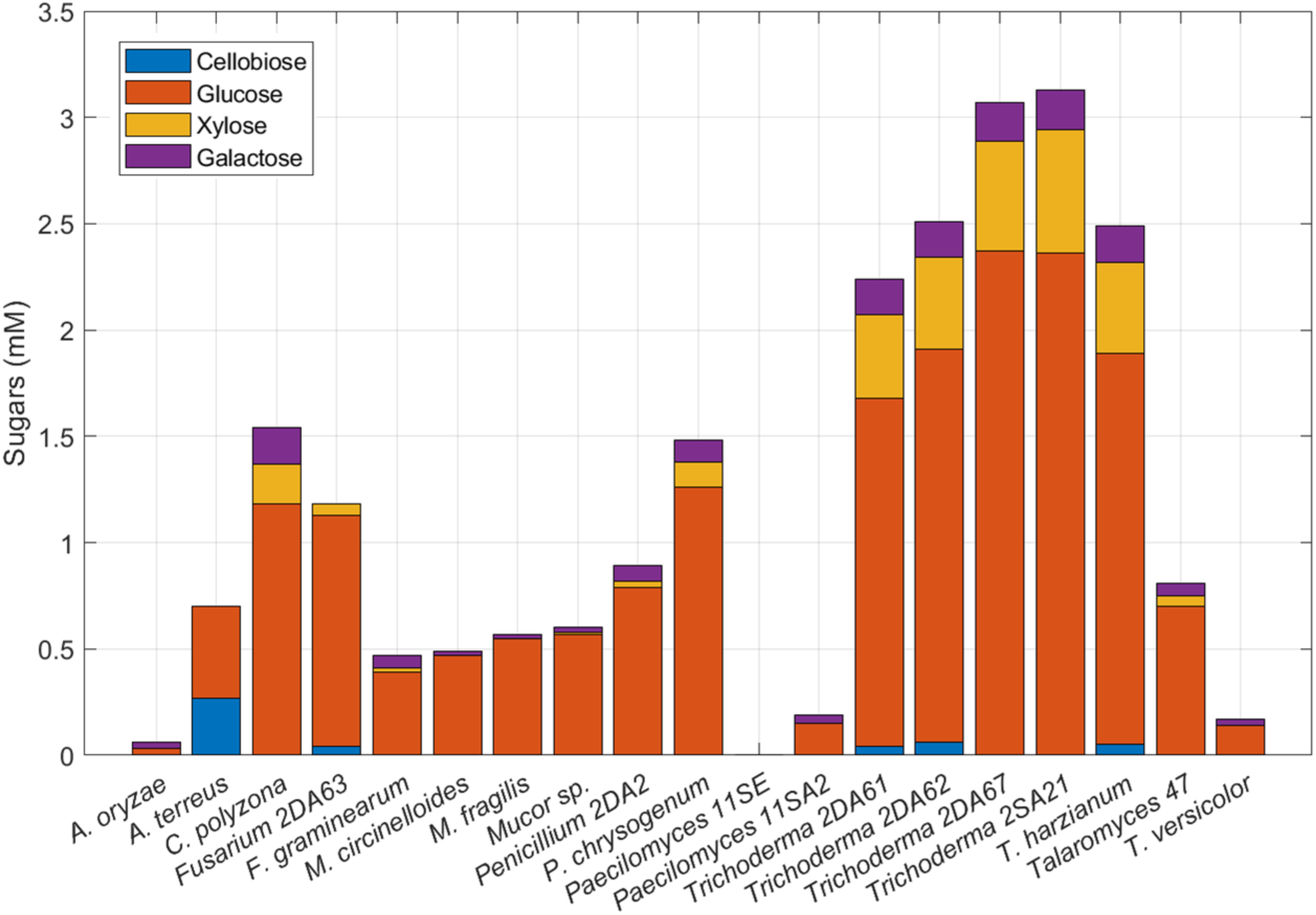
Comparative analysis of saccharification across fungal species. Stacked bar chart showing the concentration of sugars released: cellobiose (blue), glucose (orange), xylose (yellow), galactose (purple) produced by 19 different fungal strains. The sugars (in negligible quantities) present before saccharification were subtracted from the post-saccharification quantification.

#### 3.1.1. Other type of enzymatic activity

Going into more detail, two fungi demonstrated unexpected behavior, exhibiting enzymatic activity without corresponding sugar release. *A. oryzae* showed moderate multi-enzyme activities (endo-xylanase: 0.20 U.L^-1^, β-glucosidase: 0.65 U.L^-1^) and protein secretion (0.09 g.L^-1^), yet released minimal sugars (0.06 mM total). This aligns with Coutinho *et al*., (2009), who demonstrated that *Aspergillus* spp. grow poorly on cellulose as a sole carbon source despite harboring significant putative cellulase genes. The observation becomes particularly relevant given flax’s composition: 85.3% cellulose, 8.3% hemicellulose, and 3.5% lignin (Abutaleb *et al*., 2020) - substantially higher cellulose content than common lignocellulosic biomasses like wheat straw, bagasse, or rice straw (25-55% cellulose, 10-40% hemicellulose, 5-35% lignin) (Sharma *et al*., 2020; Wang *et al*., 2024). As a saprophytic but non-phytopathogenic fungus (Zakaria, 2024), *A. oryzae*’s secretome appeared limited in degrading native and untreated lignocellulosic substrate. Similarly, *Paecilomyces* 11SE displayed measurable activities (endo-cellulase: 0.13 U.L^-1^, β-glucosidase: 0.29 U.L^-1^) with the lowest protein content (0.04 g.L^-1^) but produced no detectable sugars, suggesting enzyme-substrate accessibility issues.

#### 3.1.2. Efficient saccharification with limited enzymatic arsenals

In contrast, several fungi achieved substantial sugar production despite narrow enzymatic profiles. *Mucor* spp. exhibited β-glucosidase-dominated activity (0.13-0.18 U.L^-1^ for exo-enzymes) with minimal endo-activities and low protein secretion (0.07-0.08 g.L^-1^), yet efficiently released multiple sugars (glucose: 0.55-0.57 mM, xylose: 0.01 mM, galactose: 0.02 mM), indicating highly efficient exo-enzyme action on accessible substrate regions. *Talaromyces* 47 showed almost exclusively exo- and endo-cellulases activity (0.13 U.L^-1^ and 0.05 U.L^-1^ respectively), leading to relatively high glucose content (glucose: 0.7 mM, xylose: 0.05 mM). This was as much as *Penicillium* 2DA2, which had all enzymatic activity assayed at high level. This discrepancy was also reported by Fujii e*t al.,* (2009) for *Talaromyces cellulolyticus*, where its cellulase and xylanase activities on standard substrates were comparable to or lower than *Trichoderma reesei* and commercial enzymes, however, saccharification assays on various lignocellulosic biomasses at the same protein loading showed equal or superior glucose and xylose release. Although they detected some β-xylosidase activity with pNP-xylopyranoside and endo-xylanase activity with birchwood xylan substrate. Subsequent studies revealed that *T. cellulolyticus* produces a GH30-7 endo-xylanase with specificity for decorated glucuronoxylan, consequently, less activity was detected by conventional xylan assays (Nakamichi *et al*., 2019). Additionally, this strain’s GH10 xylanase showed enhanced activity on insoluble, branched xylan substrates via a carbohydrate-binding module (Inoue *et al*., 2015). Therefore, here, low/not detected enzymatic activity likely reflected the limitations/non-optimization of enzymatic assays rather than the actual saccharification capacity on flax lignocellulose. First, HPLC and enzymatic assays differ in their analytical sensitivities. Second, β-xylosidases and β-glucosidases could be assayed within a pH range of 4.0-6.0 and at temperatures between 40 and 60 °C, which correspond to their reported optimal conditions (Kirikyali *et al*., 2014; Asha *et al*., 2016; Olajuyigbe *et al*., 2016; Rizzatti *et al*., 2001; Bai *et al*., 2013). In the present study, however, the enzymatic assays were conducted at a lower temperature (25 °C) to preserve the stability of other enzymes present in the extract, which may be less thermostable. The storage stability of the enzyme extract is discussed at the beginning of Section 3.2.2.

Interestingly, *A. terreus*, with more endophytic behavior than *A. oryzea* (S. El-hawary *et al*., 2020), demonstrated an endo-dominant profile (endo-xylanase: 0.67 U.L^-1^, endo-cellulase: 0.26 U.L^-1^) with limited exo-activities and moderate protein secretion (0.05 g.L^-1^), producing exclusively glucose-related sugars (cellobiose: 0.27 mM, glucose: 0.43 mM). The cellobiose accumulation indicates incomplete degradation, probably caused by possible β-glucosidase inhibition with released glucose (Workman and Day, 1982).

#### 3.1.3. White-rot fungi: lignin modification and polysaccharide degradation

The white-rot fungi exhibited specialized profiles reflecting their ecological niches. *Trametes versicolor* displayed a distinctive enzymatic distribution where phenol oxidase represented 39% of total activity (0.01 U.L^-1^), consistent with its lignin degradation specialization (Andlar *et al*., 2018). However, minimal exo-activities and the lowest protein secretion (0.02 g.L^-1^) resulted in only 0.17 mM total sugars. *Coriolopsis polyzona* (*Trametes polyzona,* (Justo *et al*., 2017)) ranked second in phenol oxidase activity (0.33 U.L^-1^) while maintaining among higher endo-activities (endo-xylanase: 0.81 U.L^-1^, endo-cellulase: 0.54 U.L^-1^). Combined with high protein secretion (0.11 g.L^-1^), this dual capacity enabled third efficient sugar release (glucose: 1.18 mM, xylose: 0.19 mM, galactose: 0.17 mM) after *Trichoderma* spp. and *Penicillium chrysogenum*.

#### 3.1.4. Fusarium species: phytopathogenic adaptation

*Fusarium* 2DA63 presented a remarkable profile: exclusively endo-enzyme activities (endo-xylanase: 0.84 U.L^-1^, endo-cellulase: 0.60 U.L^-1^) with zero β-xylosidase and negligible β-glucosidase (0.02 U.L^-1^), yet substantial glucose (1.09 mM) and xylose (0.05 mM) production. While distinct enzymatic assay substrate affinity among co-expressed endoxylanases from single fungal species have been documented – for instance, within the GH12 family of *Fusarium graminearum* (Habrylo *et al*., 2012) – the present data could suggest the presence of processive endoglucanases. Those enzymes remain substrate-attached, repeatedly cleaving bonds and releasing oligo-glucans, cellobiose or monomers of glucose. While predominantly bacterial (Chen *et al*., 2025), fungal processive endoglucanases with preference toward long polymers in enzymatic assays have been identified in *Aspergillus ochraceus* and *Myceliophthora thermophila* (Asha *et al*., 2016; Karnaouri *et al*., 2017). This aspect is discussed in Section 3.2.2. Briefly, enzymes found in *Fusarium* secretome belong to families with reported processive action. Additionally, its highest phenol oxidase activity (0.25 U.L^-1^) and moderate protein secretion (0.08 g.L^-1^) reflect the genus’s phytopathogenic nature, where colonization through xylem vessels requires both polysaccharide and lignin modification capabilities (Roncero *et al*., 2003). This laccase activity aligns with findings for *F. nygamai* (Arif *et al*., 2024). In comparison, *F. graminearum* produced half the sugars (glucose: 0.39 mM, xylose: 0.02 mM) with lower endo-xylanase activity (0.12 U.L^-1^) but maintained high phenol oxidase (0.16 U.L^-1^), showing an intermediate profile between saccharification and lignin modification.

#### 3.1.5. High performers: P. chrysogenum and Trichoderma species

*Penicillium* spp. is an ubiquitous mold, easily dispersed through air and thriving in almost every environment: soil, feed, decaying organic matter, indoor environments, and food. Consequently, it is widely used in food fermentation and biotechnological applications for enzyme and secondary metabolite production (Akaniro *et al*., 2023). Both *Penicillium* 2DA2 and *P. chrysogenum* demonstrated among the highest total enzymatic activities, but the latter produced 47% more total sugars, probably due to superior endo-cellulase (0.34 U.L^-1^) and β-glucosidase (0.33 U.L^-1^) activities. Despite high protein secretion (0.11 g.L^-1^ - second highest overall), *P. chrysogenum* released approximately 1 mM fewer sugars (glucose: 1.26 mM; xylose: 0.12 mM) than *Trichoderma* species, suggesting suboptimal enzyme synergy, adaptation to substrate or substrate penetration.

The five *Trichoderma* strains consistently excelled in sugar production (total sugars: 2.03-2.89 mM), correlating with the highest protein secretion (0.10-0.11 g.L^-1^), consistent endo-enzyme activities across strains (endo-xylanase: 0.98-1.15 U.L^-1^, endo-cellulase: 0.35-0.46 U.L^-1^), and substantial exo-enzyme activities, particularly β-glucosidase (0.28-0.44 U.L^-1^). Notably, *Trichoderma* 2DA62 maintained comparable xylose levels (0.39 mM) and total sugar level, despite having the lowest β-xylosidase (0.14 U.L^-1^) among *Trichoderma* spp., while *Trichoderma* 2SA21 with the highest β-glucosidase (0.44 U.L^-1^) produced the most glucose (2.37 mM). *T. harzianum*, despite the lowest β-glucosidase among *Trichoderma* spp. (0.28 U.L^-1^), matched *Trichoderma* 2DA62’s output (glucose: 1.84 mM), suggesting compensatory mechanisms or differing enzyme-substrate interactions not fully captured by measured activities. These results confirm *Trichoderma* spp. as the principal genus used for cellulase and xylanase production (Dhaver *et al*., 2022; Fang *et al*., 2019), reflecting its saprophytic lifestyle that evolved to release abundant polysaccharide hydrolases, even though its primary lifestyle is mycoparasitism with abundant chitinase release (Do Vale *et al*., 2012; Zhang *et al*., 2014). Their medium phenol oxidase activities (0.11 U.L^-1^), similar to several other fungi, had minimal impact on their superior saccharification performance.

#### 3.1.6. Correlation analysis: enzyme synergy over individual activity

To assess relationships between released sugars and enzymatic activities, Pearson correlation analysis was performed (Supplementary Data 1, **Fig. S1**). Given flax’s predominantly cellulosic composition, glucose was expectedly the major monosaccharide released, produced by most strains even without complete enzymatic arsenals (*Mucor* spp., *F. graminearum*, *Talaromyces* 47). Cellobiose showed negative or near-zero correlations with all measurements except endo-activities, indicating that accumulating fungi possess insufficient β-glucosidase activity, experience cellobiose inhibition (Nong *et al*., 2024), or glucose-mediated feedback inhibition (Bohlin *et al*., 2010). Indeed, β-glucosidases exhibit differential glucose tolerance across four classes: Class I (strongly inhibited by low glucose concentrations), Class II (glucose-tolerant), Class III (stimulated by low and inhibited by high glucose concentrations), and Class IV (not inhibited at high glucose concentrations) (Salgado *et al*., 2018). This hypothesis however remains highly speculative in the absence of inhibitory experiments on the enzymes studied.

Similarly, β-xylosidase showed weak correlation with xylose release due to possible monosaccharide inhibition (Rohman *et al*., 2019) and hemicellulose complexity. Flax’s low hemicellulose proportion combined with xylan’s heteropolysaccharide structure requires concerted action of various xylan-degrading enzymes: α-l-arabinofuranosidase (EC 3.2.1.55), α-d-glucuronidase (EC 3.2.1.139), acetylxylan esterase (EC 3.1.1.72), and p-coumaric/ferulic acid esterases (EC 3.1.1.73) to release side chain substituents. Endo-β-1,4-xylanase then hydrolyses internal β-(1,4) linkages to produce xylooligosaccharides, while β-xylosidase removes xylose from non-reducing termini (Polizeli *et al*., 2005). Consequently, endo-activities showed strong correlations with monosaccharide release (endo-xylanase/xylose r = 0.818, endo-cellulase/glucose r = 0.88), explaining why *Fusarium* 2DA63 (endo-dominant, minimal exo-activity) achieved substantial sugar production while *P. chrysogenum* (highest exo-enzymatic activity) produced less than *Trichoderma* species. A subtle relation must be present between released monosaccharides and enzyme characteristics, as (Rodrigues *et al*., 2015) and (Berlin *et al*., 2007) observed a significative hydrolysis increase on pretreated wheat straw and corn stover with β-glucosidase (Novozym 188) supplementation to Celluclast 1.5L and Cellic® CTec2, and at the same time the glucose and gluconic acid (oxidized glucose by oxidative enzymes like GH 61 family) increasing concentration inhibit β-glucosidase activity (Cannella *et al*., 2012). Liang *et al*., (2025) observed similar patterns among seventeen fungal strains (predominantly *Trichoderma* spp., *Aspergillus* spp., and *Penicillium* spp.): despite 20-60% cellulose conversion rates, correlation with β-glucosidase activity was negligible (r = 0.041), while endoglucanase and cellobiohydrolase, showed relatively high correlations with glucose (r = 0.5 and 0.71, respectively). This reflects cellulose’s unbranched β-1,4-linked D-glucose backbone requiring more coordinated action of endoglucanases (endo-1-4-β-glucanase; EC 3.2.1.4) for random internal cleavage and exoglucanases (EC 3.2.1.91) for cellobiose release from chain ends or oligosaccharides, than β-glucosidases activity (EC 3.2.1.21) for cellobiose-to-glucose conversion (Andlar *et al*., 2018). For instance, (Gao *et al*., 2010) determined than the optimal blend giving higher glucose and xylose yield had to have 3.5 to 8-fold more cellobiohydrolase, endo-glucacase and endo-xyalanase activities than exo-glucosidase and exo-xylosidases activities. This endo-exo synergism proves critical, as endoglucanase increases cellobiohydrolase-catalyzed cellulose hydrolysis rate constants (Fang *et al*., 2019; Jalak *et al*., 2012). Phenol oxidase activity showed moderate correlation with monosaccharide release (r = 0.29 - 0.48), suggesting that while lignin modification may facilitate saccharification, its impact diminishes with low substrate lignin content (Müller *et al*., 2015). Liang *et al.,* (2025) reported similar weak correlation between Lytic Polysaccharide Monooxygenase (LPMO) activity and glucose release from pretreated poplar wood (cellulose ± 52%, hemicellulose ± 31%, lignin ± 19% (Krutul *et al*., 2019)), despite LPMO’s established role in oxidizing glycosidic bonds in crystalline polysaccharides (Rani Singhania *et al*., 2021).

Overall, the lack of strict correlation between specific enzymatic activities and sugar release indicates that enzyme synergy could matter more than individual enzyme levels, product inhibition limits some high-activity producers, and substrate accessibility varies with fungal penetration strategies. It could also be due to phenomena not yet elucidated that cannot be observed through our rather simple enzymes activity assay and sugar quantification.

### 3.2. SSF vs SmF comparison

The six best candidate strains (*A. terreus*, *C. polyzona*, *Fusarium* 2DA63, *P. chrysogenum*, *Trichoderma* 2SA21, and *T. harzianum*) were grown under submerged fermentation to produce lignocellulosic enzymes to compare their performance with already obtained SSF results and determine the most suitable process for subsequent mixing steps.

#### 3.2.1. Total protein quantification

Submerged fermentation yielded substantially higher protein concentrations compared to SSF across all fungal species examined. The average protein concentration in SmF (0.38 ± 0.13 g.L^-1^) was approximately 4-fold higher than in SSF (0.08 ± 0.02 g.L^-1^) (**Fig. 3**). *Trichoderma* 2SA21 exhibited the highest protein production in SmF (0.62 ± 0.06 g.L^-1^), representing a 5.9-fold increase compared to its SSF counterpart (0.11 ± 0.01 g.L^-1^). Similarly, *Coriolopsis polyzona* demonstrated a 4.5-fold increase (0.42 ± 0.03 g.L^-1^ in SmF vs 0.09 ± 0.01 g.L^-1^ in SSF). This is not in accordance with (Arif *et al*., 2024) where the total protein content were equivalent in SF and SSF modes and 3-4 times higher than SmF from days 5-9, nor (Leadbeater *et al*., 2025) with two fold more proteins in SSF, but there was 23 times more substrate initially in SSF. Although, it was in accordance with (Liu *et al*., 2020), with twice more proteins in SmF than in SFF was found, where the fermentation of white-rot fungus *Phanerochaete chrysosporium* step was close to those implemented here (3 g of lignocellulosic substrate for SSF and 1 g for SmF).

**Fig. 3:**
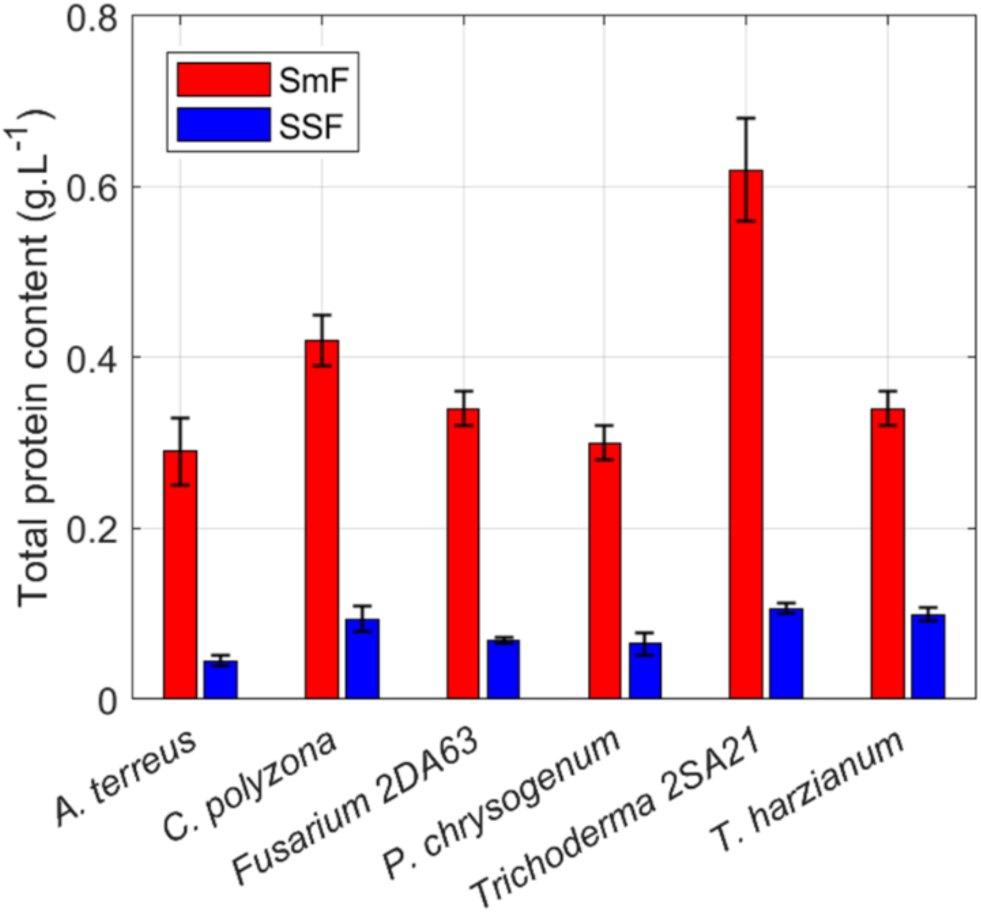
Total protein content measured (n = 3) on concentrated enzyme extracts (pooled biological triplicate) from six fungi grown with submerged fermentation (SmF) and solid-state fermentation (SSF). Extraction after 7 days of culture on 2 g flax.

Several hypothesis could be discussed upon those discrepancies: a review of the literature on xylanase and cellulase production from various fungi and lignocellulosic substrates reveals that “standard” extraction protocols involve shaking at 100-200 rpm, temperatures ranging from 4 to 37 °C, and durations of 10 min to 2 h. Various extraction buffers have been employed, including 50 mM acetate or citrate buffer (pH 4.8-5.0), with or without surfactants, though water alone has proven sufficient for maximum recovery in several cases, and often no protease inhibitors were used (Irfan *et al*., 2014; Pal and Khanum, 2010; Alananbeh *et al*., 2024; Azzouz *et al*., 2020; Sankar *et al*., 2023; Leadbeater *et al*., 2025; Pirota *et al*., 2013; Kalsoom *et al*., 2019; Teixeira da Silva *et al*., 2016; Liu *et al*., 2020). Additionally, no sequential extraction was reported, as (Pirota *et al*., 2013) demonstrated that most endoglucanase activity was recovered during the first extraction step. Therefore, our extraction protocol was designed as a consensus of the conditions most commonly reported in the literature, consisting of 50 mM sodium acetate buffer (pH 5.0), with stirring at 200 rpm for 1 h at 30 °C.

Another point is the initial substrate quantity which significantly influences enzyme production in SSF. Studies using 5-30 g of substrate in 250 mL Erlenmeyer flasks (bottom diameter approximately 140 mm) have shown that 6-10 g typically yields maximum enzyme activity (Kavya and Padmavathi, 2009; Irfan *et al*., 2014; Lee *et al*., 2018). At higher loadings, enzyme production decreases, likely due to limitations in oxygen transfer, mineral availability, reduced mycelial penetration into the compacted substrate bed, or carbon source inhibition. In the present study, 2 g of biomass in 90 mm Petri dishes was used, which should not impose growth limitations, though it may not represent maximum production capacity. This lower substrate quantity could partly explain the lower absolute protein yields compared to studies employing 5- 10 g of biomass. Although, a similar trend was observed by (Abdeljalil *et al*., 2014) with *Stachybotrys microspora* cultivated on 20 g wheat bran, where SmF yielded approximately 10-fold more total protein than SSF. Moreover, their xylanase and cellulase activities were comparable or even twice as high in SSF, resulting in greater specific enzymatic activity (*cf.* 3.2.2). Unfortunately, their extraction procedure involved grinding the mycelium after fermentation, which may have introduced contamination from intracellular proteins.

Then, standard aqueous extraction methods may underestimate total enzyme production, as a significant fraction of proteins can remain tightly bound to the lignocellulosic substrate. To date, only Leadbeater *et al*., (2025) have investigated substrate-bound enzymes using a biotin-tagging approach capable of recovering proteins attached to the biomass. Interestingly, their study revealed no significant differences in insoluble protein profiles between SSF and SmF conditions, suggesting that similar quantities and types of proteins remain (label-free MS quantification) associated with the substrate regardless of fermentation mode. This finding implies that protein losses due to substrate binding may be comparable in both systems.

Also, the relationship between total protein content and enzyme activity is not strictly linear during fungal growth. Irfan *et al*., (2014) monitored daily protein content and xylanase activity throughout the fermentation process and observed that while protein content increased linearly from day 1 to day 7 before declining at a similar rate, xylanase activity increased sharply at day 2 and remained relatively constant until day 10, with a minor peak at day 7. This temporal disconnect suggests that the nature of secreted proteins shifts during fermentation – from substrate-degrading enzymes (xylanases, cellulases) in early phases to proteins supporting fungal maintenance and development in later stages. Such shifts may be more pronounced in SmF, where nutrient dynamics differ substantially from SSF (Salgado-Bautista *et al*., 2020; Roy *et al*., 2013).

The next difference concerns the biomass moisture. Moisture content is a critical parameter for SSF performance. Several studies have reported optimal initial moisture levels of 70 - 75%, with enzymatic activity decreasing - sometimes by more than half - at higher moisture contents (80 - 130%) (Kavya and Padmavathi, 2009; Kalsoom *et al*., 2019; Pal and Khanum, 2010). The consensus is that the substrate must be sufficiently hydrated to support fungal growth under conditions mimicking their natural environment, however, optimal moisture content is substrate-dependent, as different biomasses and particle sizes exhibit distinct water absorption capacities (Lin *et al*., 2016). No fermentation studies with flax shives were identified in the literature. Preliminary experiments in this study indicated that 70% moisture was insufficient to completely wet the flax shive substrate. Complete hydration required 85.7% moisture content. An additional 4 mL of water was added to compensate for evaporation over the 7-day incubation period.

Stirring is another parameter that may explain discrepancies between studies. While SSF benefits from static conditions that allow gradual fungal colonization, most SmF studies employ relatively high agitation rates of 100–200 rpm (Abdeljalil *et al*., 2014; Liu *et al*., 2020; Leadbeater *et al*., 2025; Somadder *et al*., 2025; Teixeira da Silva *et al*., 2016), which may adversely affect fungal growth and morphology. In the present study, a lower agitation rate of 60 rpm was used to ensure adequate medium mixing while minimizing mechanical stress, potentially favoring fungal development and protein secretion compared to higher agitation rates. Although, the effect of agitation on fungal enzyme production remains context-dependent. (Sankar *et al*., 2023) did not report stirring conditions, while Arif *et al*., (2024) maintained SmF under static conditions, yet SSF generally yielded higher protein content across four different strains. An *et al*., (2016) compared stirred (150 rpm) and static SmF for *Pleurotus ostreatus* and found that stirring increased cellulase, xylanase, and laccase activities by 2- to 3-fold, though protein content was not measured. Ibrahim *et al*., (2015) investigated the impact of agitation on *Aspergillus niger* growth in SmF and determined that mycelium growth and pectinase activity increased with stirring speed up to 150 rpm; beyond this threshold, hyphae existed as free mycelium unable to maintain viability, leading to decreased enzyme activity. Cui *et al*., (1997) described that these observations reflect antagonistic effects: at low agitation, compact fungus pellets form with limited free mycelium, whereas increasing shear stress mechanically fragments pellets, with each fragment potentially initiating new pellet formation. On the other hand, improved agitation enhances oxygen transfer, promoting pellet growth. This could explain increased protein production under moderate agitation, as each newly formed pellet may independently produce its enzymatic arsenal.

Finally, these could be a distinct metabolic regulation mechanism. In SSF, enzymes remain localized near the mycelium and substrate, facilitating efficient nutrient provision without triggering additional enzyme secretion. Glucose, a catabolite repressor, can inhibit further enzyme production when present at high local concentrations (Suto and Tomita, 2001; Viniegra-González *et al*., 2003). Conversely, in SmF, hydrolysis products rapidly diffuse away from the site of production, potentially reducing local glucose concentrations below repressive thresholds and allowing continued enzyme secretion. This dilution effect may paradoxically stimulate enzyme production in SmF as the fungus responds to perceived nutrient scarcity (Leadbeater *et al*., 2025).

At the end, due to the considerable variability in experimental procedures across studies - including differences in substrate type and quantity, moisture content, agitation conditions, and extraction methods - it remains unclear why the majority of studies report higher enzyme production in SSF while some, including the present work with multiple fungal strains on a single substrate, observe the opposite trend. This should ideally be studied at a larger scale, varying both condition and fungi/substrate couples, with possible pre-treatment of the latter to limit enzyme adsorption typically by making it more hydrophobic for example.

#### 3.2.1. Analysis of the protein composition of secretomes

Following the quantitative analysis of proteins secreted (and recovered) after each mode of fermentation (SmF *vs.* SSF), a characterization of the protein families content was performed using electrophoresis (**Fig. 4**) and peptidomics analysis (**Table 1**). In this objective, after ultrafiltration, protein extracts from SmF and SSF were desalted and further concentrated using precipitation. As seen with the total protein content (**Fig. 3**), SmF conditions leading to more loaded wells with bright bands (**Fig. 4**), and more bands in general, consistent with more proteins identified with mass spectrometry (**Table 1**). A full list of identified proteins is available in Supplementary Data 2. These proteomic results will be discussed in general terms in this section, and the detailed results, including the different families and enzymes identified, will be supplemented in sections 3.2.2 and 3.2.3 below in light of the enzymatic activities measured.

**Fig. 4:**
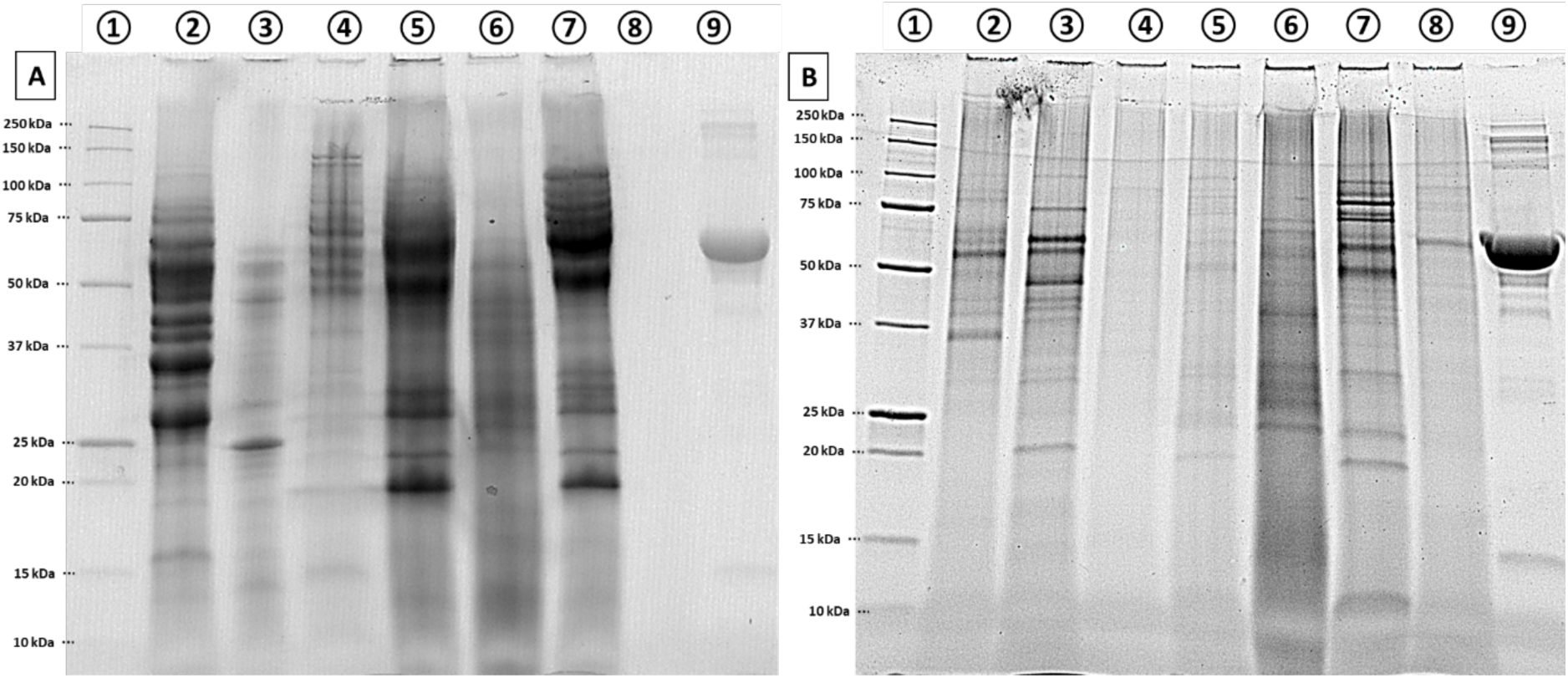
SDS-PAGE of concentrated proteins extracts, stain-free imaging. 1: Mark Ladder. 2: *P. chrysogenum.* 3: *Fusarium* 2DA63. 4: *A. terreus.* 5: *Trichoderma* 2SA21. 6: *C. polyzona.* 7: *Trichoderma harzianum.* 8: Negative control. 9: Bovine Serum Albumin (BSA). (A) SmF (B) SSF. Standards migration fronts were fit to the molecular weight through polynomial curve of degree four (R² = 0.99). To allow for a clear visualization of protein bands, contrast and sharpness were independently adjusted for the SSF gel. BSA was loaded at equal concentrations (20 μg) on both gels to serve as an internal loading and intensity reference

**Fig. 5:**
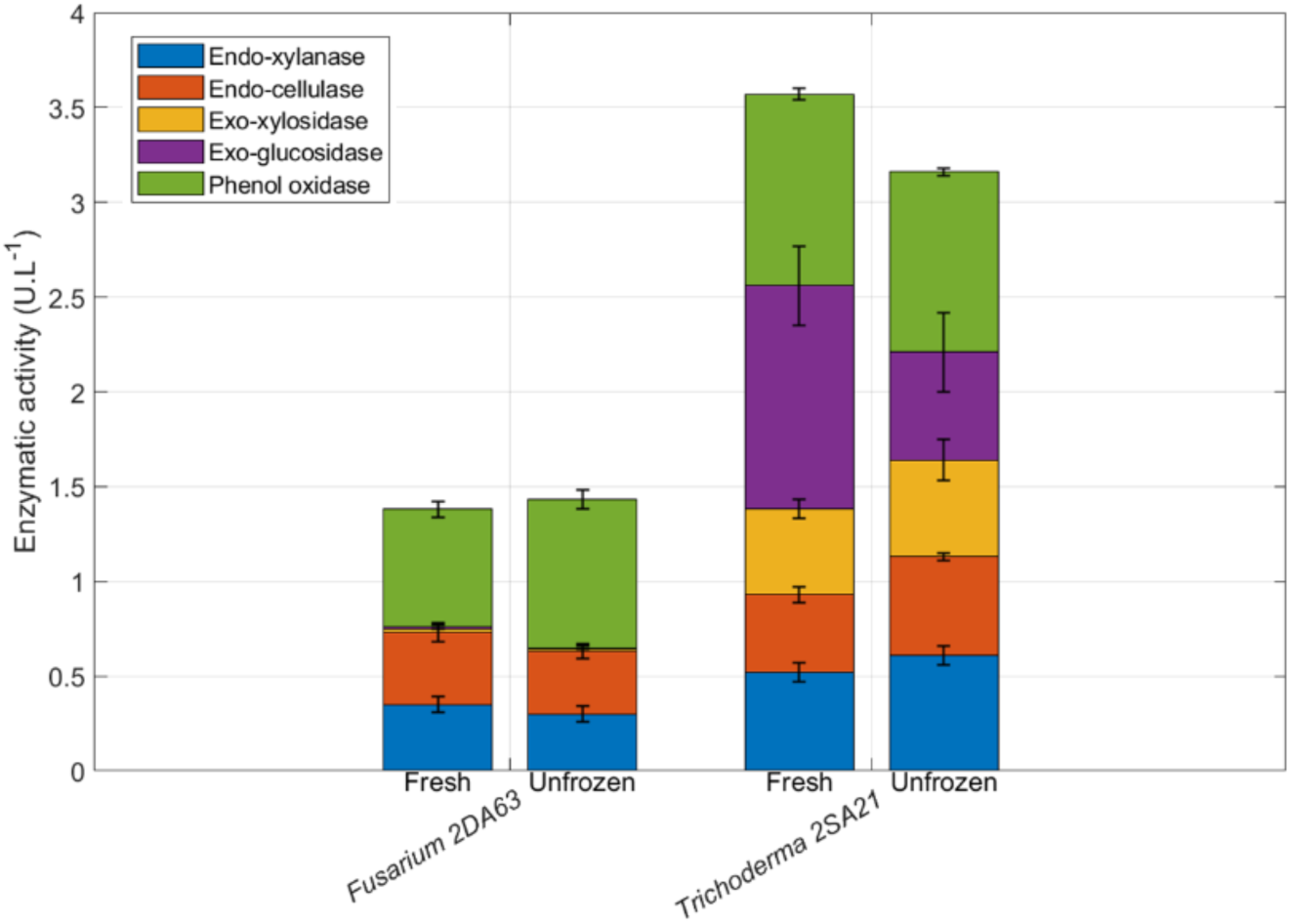
Enzymatic activity profiles of *Trichoderma* 2SA21 and *Fusarium* 2DA63 under submerged fermentation using fresh (right after ultrafiltration step) and -80°C stored concentrated enzymes extracts. Enzyme activities measured include: endo-xylanase (blue), endo-cellulase (orange), exo-xylosidase (yellow), exo-glucosidase (purple), and phenol oxidase (green). Error bars represent standard deviation (n = 3).

**Table 1:** Summary of identified proteins by peptidomics per fungus and per fermentation condition (SmF and SSF) classified into four main identified metabolisms, presented in percentages. The protein biological functions classification is detailed in Supplementary Data, **Table S7**.

| Condition | Fungus | Biological metabolism | Pathogenesis | Polysaccharides metabolism | Stress | Total proteins* | Average of proteins** | Protein*** |
| --- | --- | --- | --- | --- | --- | --- | --- | --- |
| SmF | *Aspergillus terreus* | 23.9% | 14.9% | 40.3% | 20.9% | 67 | 17 +/- 6 | 0.29 +/- 0.04 |
|  | *Corioliopsis polyzona* | 17.5% | 35.0% | 47.5% | 0.0% | 40 | 13 +/- 5 | 0.3 +/- 0.03 |
|  | *Fusarium 2DA63* | 12.5% | 20.8% | 58.3% | 8.3% | 24 | 6 +/- 5 | 0.62 +/- 0.06 |
|  | *Penicillium chrysogenum* | 13.3% | 26.7% | 53.3% | 6.7% | 45 | 11 +/- 8 | 0.34 +/- 0.02 |
|  | *Trichoderma 2SA21* | 13.5% | 30.8% | 46.2% | 9.6% | 52 | 13 +/- 8 | 0.42 +/- 0.03 |
|  | *Trichoderma harzianum* | 11.9% | 31.0% | 50.0% | 7.1% | 42 | 11 +/- 7 | 0.34 +/- 0.02 |
| SSF | *Aspergillus terreus* | 25.0% | 0.0% | 75.0% | 3.2% | 4 | 2 +/- 1 | 0.04 +/- 0.01 |
|  | *Corioliopsis polyzona* | 0.0% | 22.2% | 77.8% | 4.7% | 9 | 5 +/- 3 | 0.06 +/- 0.02 |
|  | *Fusarium 2DA63* | 0.0% | 0.0% | 100.0% | 6.3% | 4 | 4 +/- 0 | 0.11 +/- 0.004 |
|  | *Penicillium chrysogenum* | 0.0% | 29.4% | 64.7% | 6.1% | 17 | 6 +/- 4 | 0.1 +/- 0.01 |
|  | *Trichoderma 2SA21* | 6.3% | 40.6% | 53.1% | 7.7% | 32 | 11 +/- 6 | 0.09 +/- 0.01 |
|  | *Trichoderma harzianum* | 2.8% | 36.1% | 55.6% | 6.9% | 36 | 9 +/- 8 | 0.07 +/- 0.001 |

\* Total unique identified proteins per fungus
\*\* Average of identified proteins for the four selected metabolisms +/- standard error
\*\*\* Total protein content +/- standard error ( $\text{g.L}^{-1}$ ) determined by Bradford assay

Therefore, on a general aspect, a total of 391 proteins were identified across all fungal species in SmF, compared to 131 proteins in SSF (2.9-fold difference). Among this, in SmF 5 proteins were “Domain of Unknown Function” and 5 uncharacterized proteins. For SSF, 3 proteins were “Domain of Unknown Function”. Those proteins were excluded from repartition analysis, leading to a total of 371 proteins with annotation in SmF and 128 proteins with annotations with SSF. Among them, especially for SmF, several homologue enzymes were expressed. Counting only unique proteins found, the number fallen to 270 total unique proteins in SmF and 102 proteins in SSF.

From SDS-PAGE and proteomics summary a clear expression difference was seen. Unfortunately, the high loading BSA contaminated the negative control well in SSF experience (**Fig. 4B**). Another SDS-PAGE was made to confirm no protein were present in negative control, available in Supplementary Data 1, **Fig. S2**.

The number of distinct secreted proteins was markedly higher in SmF conditions. At the individual species level, *Trichoderma harzianum* showed the least dramatic shift, secreting 36 proteins under SSF versus 42 proteins under SmF. Notably, *Aspergillus terreus* exhibited the largest fold-change in proteomic detection, with only 4 distinct proteins in SSF increasing to 67 proteins in SmF.

Polysaccharide metabolism proteins constituted the dominant category in both conditions, but with different proportions. In SSF, 71.0% of identified proteins were involved in polysaccharide metabolism, compared to 49.3% in SmF. Pathogenesis-related proteins showed a contrasting pattern, with relative higher abundance in SmF (26.5%) compared to SSF (21.4%). *Trichoderma* species exhibited the highest pathogenesis-related protein expression in both conditions, with *T. harzianum* reaching 36.1% and *Trichoderma* 2SA21 achieving 40.6% in SSF. Fungal biological metabolism proteins were more abundant in SmF (15.4%) than in SSF (5.7%). Stress-response proteins remained relatively low in both conditions with slight increase in SmF (8.8%) compared to SSF (5.8%). *A. terreus* exhibited the highest stress response in SmF (20.9%).

Still from a macro perspective, when comparing SDS-PAGE (**Fig. 4**) and MS analysis (Supplementary Data 2), only *A. terreus* (lane 4) presented a striking contrast between fermentation conditions. In SmF, multiple bands spanning a wide molecular weight range were detected, while in SSF, only one band was slightly visible at 90 kDa with no reliable MS identification, likely due to insufficient protein content, stringent filtration parameters (minimum of 2 unique peptides, false discovery rate < 0.1%), or effects of denaturing conditions on protein integrity and migration patterns. Otherwise, the gel bands correlated well with proteomics data in both fermentation conditions, with proteins showing high spectral counts (> 100 spectra in SmF; > 50 spectra in SSF) consistently appearing as more intense bands.

Looking now, still on a macro scale, to the enzyme activities detected, a comprehensive enzymatic toolkits for plant cell wall degradation was effectively observed across all fungi tested, including enzymes targeting cellulose (cellulases, endoglucanases, β-glucosidases, lytic polysaccharide/cellulose monooxygenases), hemicellulose (xylanases, β-xylosidases, glucanases, mannanases, arabinofuranosidases), and auxiliary activities that deramify complex polysaccharides (acetylxylan esterases, feruloyl esterases, carboxylic ester hydrolases, various carbohydrate esterases) or degrade the pectin matrix (polygalacturonases, pectate lyases, rhamnogalacturonases) (Andlar *et al*., 2018; Lopes *et al*., 2018; Malgas *et al*., 2019; Østby *et al*., 2020; Østby and Várnai, 2023).

The presence of both endo-acting enzymes (which create internal breaks in long polysaccharide chains, generating new chain ends and helping to depolymerize and deramify complex glucans) and exo-acting enzymes (which remove terminal sugar units from non-reducing ends, completing the degradation process) demonstrates the synergistic nature of these enzyme systems across both SmF and SSF conditions (Jalak *et al*., 2012). Additionally, the oxidative enzymes (lytic polysaccharide monooxygenases from AA9 family) provide an alternative, oxidative mechanism to disrupt crystalline cellulose and lignin, complementing the hydrolytic activities and creating entry points for other enzymes in both fermentation systems (Müller *et al*., 2015; Rani Singhania *et al*., 2021; Janusz *et al*., 2017; Kumar and Chandra, 2020).

Finaly, while direct lignin-degrading enzymes were not extensively identified in these proteomic analyses (one tyrosinase copper-binding domain-containing protein found in *A. terreus* SmF with 23 spectra, one peroxidase and one catalase-peroxidase found in *T. harzianum* in SSF and SmF with 4 and 36 spectra respectively, and one catalase-peroxidase found in *Trichoderma* 2SA21 in SmF with 8 spectra) for either fermentation condition, the phenol oxidase activity measured in the enzymatic activity assays (*cf.* 3.2.2) (particularly high in SmF conditions) could be due highly active but lower expressed enzymes, or LPMO interfering, as monophenols were found to be preferred electron donors for LPMO activity (Frommhagen *et al*., 2016; Liao *et al*., 2025)

#### 3.2.2. Enzymatic activity measured in secretomes

In parallel of the proteomic study, five total enzymatic activities were measured in the secretomes produced by the six fungi selected (*A. terreus*, *C. polyzona*, *Fusarium* 2DA63, *P. chrysogenum*, *Trichoderma* 2SA21, and *T. harzianum*) and two fermentation modes (SSF *vs.* SmF), to follow their main biomass degradation capability: endo-xylanase and endo-cellulase for large polysaccharides cleavage (xyloglucans, xylans, cellulose); exo-xylosidase and exo-glucosidase to release xylose and glucose monomers; phenol oxidase or laccase activity which indicate lignin degrading enzymes (Kumar and Chandra, 2020).

Before performing the measurement, the stability of concentrated enzyme extracts during storage was evaluated. Due to the duration of enzymatic assays (2 h per sample), same-day analysis following the concentration step was not feasible. Consequently, all enzyme extracts were immediately stored at - 80 °C after ultrafiltration and analyzed the following day or within the same week after slow thawing on ice. To validate this storage protocol, enzyme stability was assessed using SSF-derived extracts by comparing all five enzymatic activities after one and two freeze-thaw cycles, with a two-week interval between measurements. No significant loss of activity was detected, confirming that storage at – 80 °C with minimal freeze-thaw cycles preserves enzymatic integrity. Lyophilization was also evaluated as an alternative preservation method and resulted in substantial activity losses (approximately 50%) for endo-cellulase, endo-xylanase, and polyphenol oxidase activities (data not shown).

Direct comparison between freshly concentrated and frozen extracts was performed for two fungal strains. *Fusarium* 2DA63 exhibited no detectable activity loss after freezing, whereas *Trichoderma* 2SA21 showed a 50% reduction in exo-glucosidase activity. Despite this variability, the distribution of enzymatic activities and the qualitative enzyme profiles remained equivalent, without total loss of activity. These observations suggest that while absolute activity values may be subject to some loss during freezing, this effect was likely consistent across all strains processed under identical conditions, thereby preserving the validity of comparative analyses. Moreover, enzymes from SmF were reported to be more instable than those produces from SSF (Kar *et al*., 2013; Leadbeater *et al*., 2025; Roy *et al*., 2013), this could support that if freezing had little impact on SmF enzyme, there could be no impact on SSF enzymes from the freezing.

Now, looking at the detailed comparison of the enzymatic activities measured with each fermentation mode/strain, first of all, endo-xylanase activity - targeting the xylan backbone of hemicellulose by cleaving β-1,4-glycosidic bonds between xylose units - exhibited pronounced fermentation mode-dependent variations with a global increase under SSF conditions (**Fig. 6**). *Trichoderma* spp. demonstrated the overall highest activities: *T. harzianum* (1.14 ± 0.12 U.L^-1^) and *Trichoderma* 2SA21 (1.15 ± 0.15 U.L^-1^), representing two times increase compared to their SmF counterparts (0.52 ± 0.03 and 0.61 ± 0.05 U.L^-1^, respectively). Putting this in comparison with the proteomics analysis, for both, a probable endo-1,4-β-xylanase was detected around 24 kDa in electrophoresis gel in both conditions too (23.8 kDa with 132-155 spectra in SmF; 23.8 kDa with 57-67 spectra in SSF). Interestingly, two extra beta-xylanases were detected in LC-MS for both strains and conditions, leading to the same enzymatic diversity but not the same enzymatic activity. Moreover, they probably displayed acetylxylan esterase at approximately 30 kDa in both SmF (30.2 kDa, 171 spectra) and SSF (30.2 kDa, 22 spectra), for *Trichoderma* 2SA21 and for *T. harzianum* with 60 spectra in SmF and with 34 spectra in SSF). This enzyme removes acetyl groups from xylan, which is crucial for deramifying the hemicellulose backbone and making it more accessible to endo-xylanases and other hydrolytic enzymes (Østby and Várnai, 2023). *Fusarium* 2DA63 showed similar better activity under SSF with almost three-time increase, 0.84 ± 0.15 U.L^-1^, compared to SmF with only 0.30 ± 0.04 U.L^-1^ of activity, however only one endo-1,4-β-xylanase was identified in SSF with 4 spectra, while one endo-1,4-β-xylanase and one β-xylanase were identified in SmF with 31 and 46 spectra, respectively. *P. chrysogenum* displayed a doubling of activity under SSF (0.78 ± 0.19 *vs.* 0.38 ± 0.04 U.L^-1^). β-xylanase was detected in both conditions, appearing possibly at 34 kDa in the electrophoresis gel in SmF (35.5 kDa, 438 spectra) and 35 kDa in SSF (35 kDa, 57 spectra). In contrast, *C. polyzona* exhibited a more moderate 62% increase (0.81 ± 0.09 *vs.* 0.50 ± 0.02 U.L^-1^), yet twice more β-xylanase were identified in SmF (37.4-42.7 kDA, 6-23 spectra which one could correspond to the gel band at 39 kDa). *A. terreus* showed only 24% enhancement (0.47 ± 0.13 *vs.* 0.38 ± 0.01 U.L^-1^), although no xylanase was identified in SSF but there were four β-xylanase in SmF (ranging from 32.1-42.1 kDa and identified with 5-73 spectra)

**Fig. 6:**
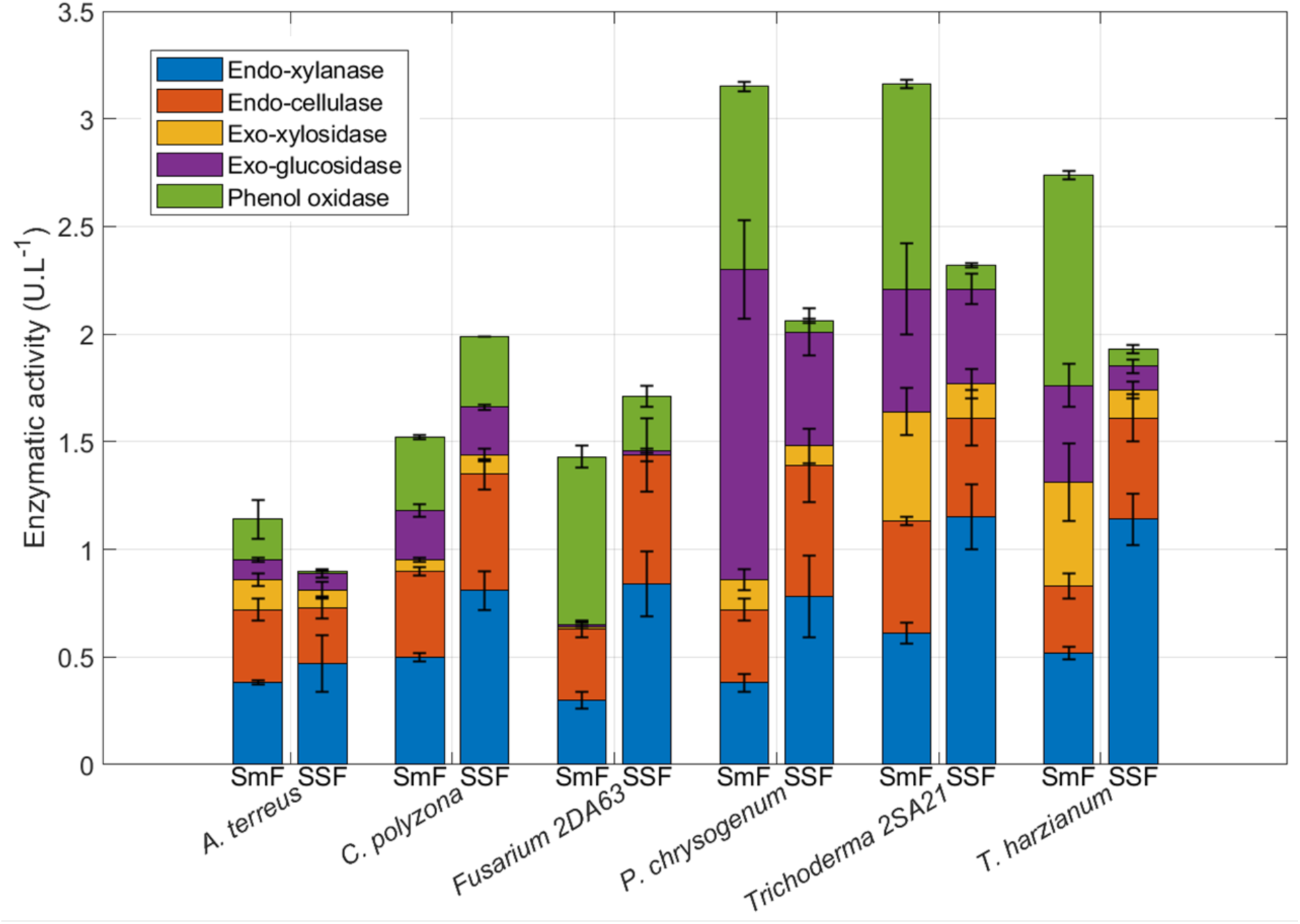
Enzymatic activities measured (n = 3) on concentrated enzyme extracts (pooled biological triplicate) from six fungi grown with submerged fermentation (SmF) and solid-state fermentation (SSF). Extraction after 7 days of culture on 2 g of flax.

Endo-cellulase activity showed more balanced responses: *C. polyzona*, *P. chrysogenum*, *Fusarium* 2DA63 and *T. harzianum* demonstrated a SSF-mediated increases by 37%, 81%, 80% and 53% respectively, while *A. terreus* and *Trichoderma* 2SA21 had similar activities, although there was no statistical difference based on fungi, only the mode of fermentation had significant influence (Supplementary Data 1, **Fig. S3**). General glucanases were the more represented enzymes found in all fungi, with 28 mass identifications in SmF (averaging 170 spectra), and 15 mass identifications in SSF (averaging 82 spectra). More specifically, endo-β-1,4-glucanases comprised eight cellulases were identified in SmF (averaging 118 spectra across all strains) and six in SSF (*C. polyzona*, *P. chrysogenum*, *Fusarium* 2DA63, *Trichoderma* 2SA21). Cellulase secretion is largely constitutive in fungi (Mach and Zeilinger, 2003), explaining their abundance. However, Sankar *et al*., (2023) reported a transient 190-fold higher cellulase abundance in SSF compared to SmF on day 5 of fermentation, which decreased drastically and showed no difference by day 9, suggesting time-dependent regulation. In the present study, the main glycoside hydrolase families identified were GH5, GH6, and GH7 in both fermentation modes, with similar proportions: GH5 (30–33%), GH6 (30%), and GH7 (40–55%).

*Trichoderma* 2SA21 and *T. harzianum* exhibited the highest SmF exo-xylosidase activities (0.51 ± 0.11 and 0.48 ± 0.18 U.L^-1^ respectively), which were greater by 69% and 73% than SSF conditions (0.16 ± 0.07 and 0.13 ± 0.04 U.L^-1^ respectively), indicating preferential SmF-based production. This enzyme removes xylose units from the non-reducing ends of xylan oligomers generated by endo-xylanases, completing hemicellulose degradation. In this case, protein identification corroborated the activity data, as uniquely in SmF, both strains could present a band at 91 kDa corresponding to xylan 1,4-β-xylosidase at 92.6 kDa (148 spectra) and at 91.2 kDa (388 spectra) for *Trichoderma* 2SA21 and *T. harzianum* respectively. No equivalent bands were present in SSF electrophoresis gel, but same enzyme was detected in LC-MS with fewer spectra (85-38). *A. terreus, C. polyzona* and *P. chrysogenum* showed no fermentation mode effects (statistically no significant) and with little activity (0.05-0.14 U.L^-1^) (Supplementary Data 1, **Fig. S5**). As for SSF mode, no such activity was detected with *Fusarium* 2DA63 in SmF, although traces of xylose were detected. Similarly, little or no exo-glucosidase activity was detected in either fermentation mode, despite the presence of glucose. A similar hypothesis could be made about a “processive” endo-xylanase, that could be expressed in both modes, but no proof exist to date (Saberi *et al*., 2024), only one enzyme belonging to GH30 family was found out with both endo activity and xylobiohydrolase activity, releasing xylobiose (Katsimpouras *et al*., 2019).

The β-glucosidase activity, which completes cellulose degradation by hydrolyzing cellobiose and short cellodextrins into glucose monomers, exhibited the highest strain-to-strain variation among all enzymes tested. *P. chrysogenum* displayed the highest activity under SmF, achieving 1.44 ± 0.23 U.L^-1^, representing 63% greater activity compared to SSF. Moreover, uniquely in SmF, *P. chrysogenum* showed a band corresponding to a possible β-glucosidase at 82 kDa (85.3 kDa, 23 spectra). Additionally, two glucan 1,3-β-glucosidase were identified, which release α-glucose. In contrast, under SSF conditions, one probable β-glucosidase and one glucan 1,3-β-glucosidase were identified but with considerably fewer spectra (5-21). *Trichoderma* spp. showed moderate activities (0.44-0.57 U.L^-1^ in SmF, 0.11-0.44 U.L^-1^ in SSF), *Trichoderma* 2SA21 uniquely showed a band at 76 kDa in SSF, which could correspond to β-glucosidase cel3A (77.2 kDa, 53 spectra), consistent with similar activities observed in both fermentation modes. For *T. harzianum*, three β-glucosidases were identified uniquely in SmF, consistent with its higher activity in this mode; one of these could correspond to a band observed at 78 kDa on gel electrophoresis, matching β-glucosidase cel3A (77.1 kDa, 117 spectra). However, proteomics data did not align with these activity measurements: for *A. terreus*, almost no bands were visible on SSF gels and no β-glucosidases were identified by LC-MS under this condition, whereas one β-glucosidase and one glucan 1,3-β-glucosidase were identified in SmF, albeit with low spectra counts (5–10). Similarly, for *C. polyzona*, two β-glucosidases were identified only in SmF despite comparable activities in both modes. To support the particular case of *Fusarium* 2DA63 without β-glucosidase activity, it was reported that enzymes with such mechanisms belonged to the GH5 and GH7 families (Payne *et al*., 2015; Wu and Wu, 2020), which was consistent with this study, *Fusarium* 2DA63 LC-MS analysis indeed revealed two cellulases and one glucanase in SSF mode, and one cellulase and two glucanases highly expressed (166-355 spectra, 55-67% coverage) in SmF mode, all belonging to these families (Supplementary Data 2).

Phenol oxidase production showed the most dramatic fermentation mode-dependent response. All strains except *C. polyzona* demonstrated substantially higher activities under SmF: *T. harzianum*: 12-fold increase in SmF (0.98 ± 0.02 *vs.* 0.08 ± 0.02 U.L^-1^), *Trichoderma* 2SA21: 8-fold increase (0.95 ± 0.02 *vs.* 0.11 ± 0.01 U.L^-1^), *P. chrysogenum*: 17-fold increase (0.85 ± 0.02 *vs.* 0.05 ± 0.01 U.L^-1^), *Fusarium* 2DA63: 3-fold increase (0.78 ± 0.05 *vs.* 0.25 ± 0.05 U.L^-1^) and *A. terreus*: 19-fold increase (0.19 ± 0.09 *vs.* 0.01 ± 0.00 U.L^-1^). This concurred with proteomics data where more oxygen-reactive enzymes were found, related to oxidative stress response (*cf.* **Table 1**) like peroxidase. Only *C. polyzona* maintained comparable phenol oxidase activity across fermentation modes (0.34 ± 0.01 *vs.* 0.33 ± 0.00 U.L^-1^), suggesting mostly constitutive expression independent of substrate physical state, which is consistent with white-rot fungi characteristics (Viswanath *et al*., 2014). Another hypothesis could be discussed, as laccase transcription was found to be strongly induced by water-soluble phenolic compounds released during lignocellulose degradation. Phenolic compounds are known for their antioxidant properties, but they are also toxic to fungi, as in the presence of oxygen, they can undergo auto-oxidation or redox cycling, generating uncontrolled reactive oxygen species, and damage membrane, proteins, and DNA (Piscitelli *et al*., 2011). On the other hand, laccases control the full oxidation of phenolic compounds. Castanera *et al*., (2012) demonstrated that laccase gene transcription in *Pleurotus ostreatus* was upregulated in induced SmF cultures but downregulated in SSF, specifically in response to water-soluble wheat straw extracts. Thus, in SmF, the greater bioavailability of soluble phenolic inducers in the liquid phase, from biomass degradation (spontaneous or enzyme-assisted), combined with a bit better dissolved oxygen concentration, due to low but present stirring, required for laccase catalytic activity, may explain the enhanced phenol oxidase activity.

When normalizing the total enzyme activity to 100% (**Fig. 7**), it is easier to identify enzymatic activity trends, and this confirms that in SmF mode, the proportion of PPO increased significantly with 3 to 14 fold change for all fungi except *C. polyzona* : *A. terreus* (from 1.5% to 16.8%), *Fusarium* 2DA63 (from 14.6% to 54.2%), *P. chrysogenum* (from 2.2% to 27.2%), *T. harzianum* (from 4.7% to 35.7%), and *Trichoderma* 2SA21 (from 4.3% to 30.1%). The trend is also shown in protein ratio identified with mass spectrometry (**Table 1**) where proteins related to polysaccharides metabolism decreased from 70% to 50% on average from SSF to SmF mode switch, with an increase of protein relative to oxidative stress, like catalase-peroxidase found with high abundance (4 to 22 peptides and 8% to 33% coverage) in *A. terreus*, *P. chrysogenum*, and *Trichoderma* spp. (Supplementary Data 2). Only one tyrosinase (23 spectra and 15% coverage) was found in *A. terreus.* Although, three times more lytic polysaccharide monooxygenase (LPMO) were found in SmF, which could react with the MBTH assay substrate, as dopamine was found to be a good reducing agent for LPMO from *Myceliophthora thermophila* (Frommhagen *et al*., 2016). Same overexpression of LPMO in SmF were observed by (Leadbeater *et al*., 2025), but not for (Liu *et al*., 2020), where 6 LPMO were found in SSF but one in SmF, maybe explained by the white-rot fungus category.

**Fig. 7:**
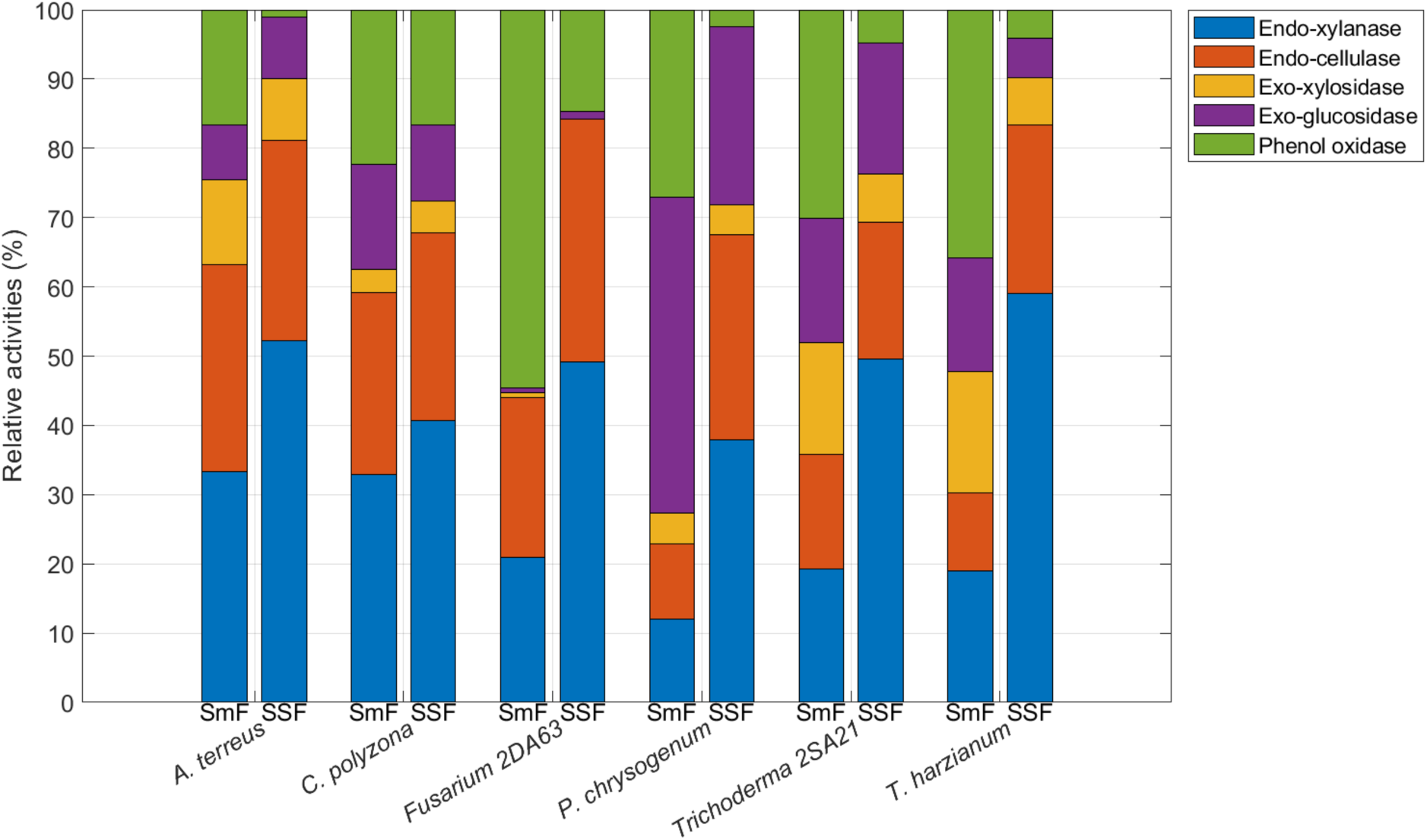
Relative enzymatic activities measured (technical replicate n = 3) on concentrated enzyme extracts (pooled biological triplicate) from six fungi grown with submerged fermentation (SmF) and solid-state fermentation (SSF). Extraction after 7 days of culture on 2 g of flax.

On the contrary, endo-xylanase activities were substantially higher under SSF despite lower total protein secretion and fewer proteins identified by proteomics. This apparent paradox suggests that SSF conditions favor the production of a more specialized and catalytically efficient enzyme repertoire rather than a diverse but dilute secretome. Xylanase expression is believed to be induced through direct contact with the xylan substrate and its own degradation products (Polizeli *et al*., 2005; Collins *et al*., 2005), which may promote the secretion of high-efficiency isoforms under SSF conditions. A more recent study by Salmon *et al*., (2014) demonstrated that induction dynamics shift over time: xylose induce xylanase activity at the beginning, but after the fifth day of fermentation, cellulose becomes the primary inducer of xylanase activity, alongside cellulase activity. Although, the proportion of endo/exo-xylanase and cellulase activity remained similar for *A. terreus* and *C. polyzona*. In (M. *et al*., 2016) the comparison between SmF and SSF *A. terreus* growth shown 1.2-48 fold more enzymatic cellulase and xylanase activity (U.mL^-1^) in SSF, although the fermentation substrate was also ten times more in quantity. *Fusarium* 2DA63 doubled its endo activities in SSF mode (49.3% and 35.2% for endo-xylanase and endo-cellulases activities respectively), although same proportion of endo enzymes were detected in both fermentation: one endo-1,4-β-xylanase, two glucanases, two cellulases for SSF and one for SmF. *P. chrysogenum* increased endo activities by 2 to 3 times, which is also not reflected on protein occurrences, as same enzymes in same number were identified in both fermentation modes (β-xylanase, cellulase, endoglucanase-4, glucanase), but reduced β-glucosidase activity by half in SSF mode (25.7% *vs.* 45.8% in SmF), supported by one extra β-glucosidase found in SmF mode. This was the most spectacular change - shift from “exo-dominant” to “endo-dominant” profile. *Trichoderma* 2SA21 exchanged the endo-xylanase (49.7%) in SSF mode in behalf of PPO, reducing it by half, while more glucanases, endoglucanases and hemicellulose ramified part debranching enzymes were detected in SmF (α-glucuronidase, α-L-arabinofuranosidase, α-L-fucosidase) (Østby and Várnai, 2023). *T. harzianum* exhibited a shift from endo- to exo-activities depending on the fermentation mode: endo-xylanase and cellulase activities increased 2- to 3-fold (58.8 and 24.3%, respectively) in SSF, despite the detection of similar numbers of endo-enzymes (glucanases and β-xylanases) in both conditions. Conversely, exo-xylanase and cellulase activities decreased three-fold in SmF, which correlated with higher abundances of β-glucosidase and β-xylosidase. As for *Trichoderma* 2SA21, a wild range of hemicellulases were detected in SmF, yet this did not translate into higher sugar yields (**Fig. 8**). *T. harzianum* showed minimal differences in unique protein profiles between fermentation modes (42 *vs.* 36). Although total protein content was consistently higher in SmF than in SSF for all strains, the overall specific enzymatic activity tended to be higher in SSF. These findings suggest that the main lignocellulosic enzymes exhibit significantly greater catalytic efficiency under SSF conditions.

**Fig. 8:**
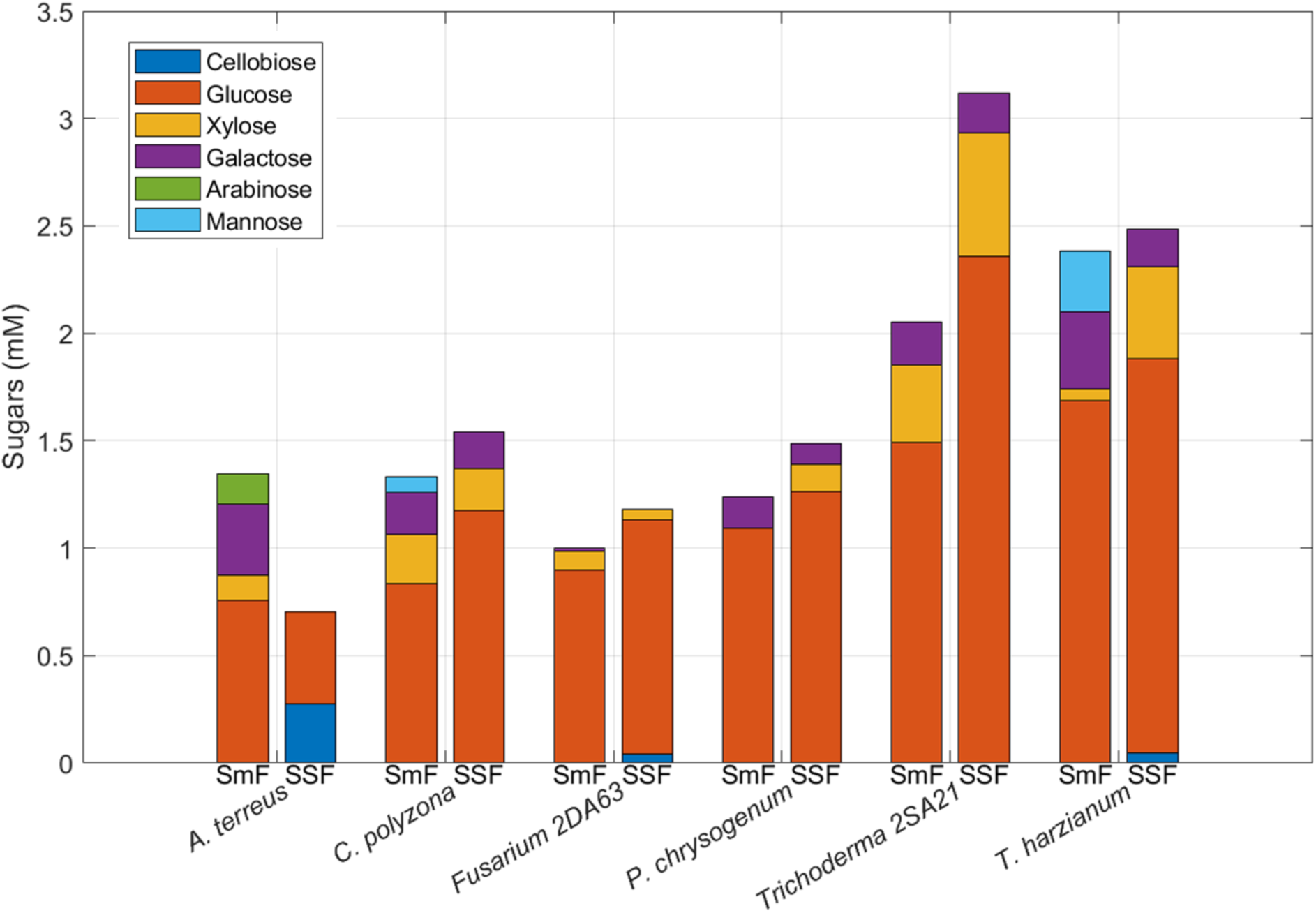
Saccharification efficiency of flax shives using fungal enzyme extracts from different fermentation modes. Crude enzyme extracts were obtained from six fungal strains cultivated under submerged fermentation (SmF) or solid-state fermentation (SSF) on flax shives (2 g) in triplicate then pooled and concentrated. Saccharification was performed at 30 °C for 48 h with (15 mg·mL⁻¹) substrate loading and 200 rpm. Sugars concentration measured before saccharification was substracted.

#### 3.2.3. Sugar production

Finally, when evaluating the saccharification efficiency of the resulting secretomes, through sugars quantification, glucose was found to be the dominant sugar across all strains and fermentation modes. This seems quite logical as biomass composition fundamentally governs both fermentable sugar yields and enzymatic accessibility, flax’s elevated cellulose proportion predicts enhanced glucose release. *Trichoderma* strains show the highest glucose levels (1.49-1.69 mM in SmF and 1.84-2.36 mM in SSF) alongside elevated endo-xylanase (0.52-0.61 U.L^-1^ in SmF and 1.14-1.15 U.L^-1^ in SSF) and endo-cellulase activities (0.31-0.52 U.L^-1^ in SmF and 0.46-0.47 U.L^-1^ in SSF). *P. chrysogenum* produced 1.09 mM glucose with notably high β-glucosidase activity (1.44 U.L^-1^, the highest among all strains in SmF), still SSF lead to 16% higher release with 1.27 mM and 63% less β-glucosidase activity (0.53 U.L^-1^). Xylose levels vary considerably and show rather inverse relationships with β-xylosidase activity. This pattern could reflect product inhibition, a well-documented regulatory mechanism for β-xylosidases. While low concentrations of xylose and glucose can stimulate exo-enzyme activity, same monosaccharides above 2 mM inhibits β-xylosidases through competitive, non-competitive, or uncompetitive mechanisms depending on the fungal species and enzyme family (Rohman *et al*., 2019; Kirikyali *et al*., 2014). Although this inhibition may appear counterproductive given that xylose is an essential carbon source for fungal pentose pathway catabolism (Chroumpi *et al*., 2021), it may serve a physiological function: since β-xylosidase is secreted extracellularly while xylose metabolism occurs intracellularly, product inhibition could prevent excessive xylose accumulation when membrane transport capacity becomes limiting. Indeed, access to sugars in complex polysaccharides depends not only on their release by hydrolytic enzymes, but also on the presence of transporters capable of effectively transporting the sugars into the cell (Sloothaak *et al*., 2016). Additionally, the endo-xylanase/β-xylosidase activity ratio could influence overall hydrolysis efficiency. Nieto-Domínguez *et al*., (2019) found that optimal xylan saccharification required endo-xylanase/β-xylosidase molar ratios of 18:1 for beechwood xylan and 125:1 for birchwood xylan, indicating that high endo-xylanase activity generating abundant xylooligosaccharides could reduce the β-xylosidase activity required for efficient xylose release. Thus, in SmF, *P. chrysogenum* showed no xylose despite moderate β-xylosidase activity (0.14 U.L^-1^), when *Trichoderma* 2SA21 enzymes produced the highest xylose content (0.36 mM) with 3.5 times more β-xylosidase activity (0.51 U.L^-1^), perhaps indicating different enzymatic cocktail synergy with different kind of enzymes. Still, the relation between main enzymatic activity and saccharification remain unclear, as (Liu *et al*., 2020) encountered also a paradox, despite lower enzymatic activities (endoglucanase, β-glucosidase, β-xylosidase), SmF enzymes achieved better saccharification yields at low substrate loading (close to present study). It was attributed to the presence of more enzymes with CBM.

Cellobiose was detected only in SSF mode (*A. terreus*, *Fusarium* 2DA63, *T. harzianum*), suggesting either the release of polymers into the medium favorized the enzyme-substrate encounter or more exo-glucosidase secretion (one and three β-glucosidases detected for *A. terreus* and *T. harzianum*) in SmF. Sankar *et al.,* (2023) found reverse results, with more cellulose released in SmF than in SMF, explained by higher β-glucosidase secretion. The SmF mode lead also to more kind of sugars generated. Particularly, *A. terreus* experienced the most dramatic shift from 4 to 67 identified proteins, increasing the number of CAZy enzymes from 4 to 32, leading to xylose (0.12 mM), galactose (0.32 mM) and arabinose (0.14 mM) with enzymes like β-xylanases, arabinogalactan endo-β-1,4-galactanase, mannan endo-1,4-β-mannosidase and α-L-arabinofuranosidase. The mannose presence with *C. polyzona* and *T. harzianum* in SmF could be explained with mannan endo-1,4-β-mannosidase presence, where it wasn’t identified in SSF.

Again, few proteins (< 10) were identified in SSF mode for *A. terreus*, *C. polyzona* and *Fusarium* 2DA63, due to low secretion, still the level of released sugars was similar to the one in the SmF where between 24 and 67 unique proteins were identified, meaning that even if the diversity of SSF was 38% lower (Supplementary Data 1, **Fig. S8**), those which remains were essential for efficient saccharification. The same conclusion was made by (Leadbeater *et al*., 2025) where despite different enzyme profiles, both fermentation achieved comparable peak saccharification activity. The SSF was more specialized, with lower diversity, dominated by smaller subset of key highly abundant enzymes (GH1, GH3, GH6, and GH7 families), with more stable expression (1-21 days). On the other hand, SmF lead to greater diversity of CAZyme families, with more volatile expression over time (rapid initial enzyme secretion followed by decline). Also, some GH families were found specifically in SmF or SSF, but in the present study extended to six fungi, only one β-galactosidase from GH2 family was found only produced by *P. chrysogenum* in SSF, while 52 CAzyme families were represented in SmF mode. In the SSF, there was no CE3, CE4, CE5 (acetylxylan esterase, cutinase), CBM20, GH13 (α -amylase), GH17 (Glucan 1,3-β-glucosidase), GH20 (β-hexosaminidase), GH25, GH27 (α-galactosidase), GH31 (α-glucosidase), GH53 (arabinogalactan endo-β-1,4-galactanase), GH67 (α-glucuronidase), GH72 (1,3-β-glucanosyltransferase), GH92 (Exo-mannosidase), GH93 (Exo-arabinanase), GH131, GT, PL3 (Pectate lyase) and PL4 (rhamnogalacturonate lyase), still some are related to primary structure breakdown (Supplementary Data 1, **Fig. S9**). Due to few enzyme secretion in SSF mode, the data a less reliable, but as (i) the enzymatic activity and sugar production were similar in both mode and (ii) few but essential enzymes for lignocellulosic hydrolysis was found in SSF (glucanases, xylanases, LPMO) it can be concluded and confirmed with previous studies that filamentous fungi generally undergo less drastic conditions (water, stirring) producing highly stable enzymes or with less proteinases (7 found in SSF and 26 in SmF), leading to a higher mass specific efficiency (Sankar *et al*., 2023; Mazumder *et al*., 2009). Leadbeater *et al*., (2025) explained that enzyme cocktails derived from SmF may contain increasingly greater levels of redundant proteins and inhibitors, which may account for the superiority of SSF at later time points despite a constant CAZyme profile at the family level. In SSF, enzymes secreted into the solid matrix are more likely to remain localized near or on the substrate, enhancing both their stability and effectiveness, whereas enzymes in SmF are diluted in the liquid phase, reducing their stability and effective concentration at the substrate interface.

To benchmark the efficiencies detailed above, a comparison was made with commercial enzyme preparations, with the study of the concentration of sugars produced relative to the mass of proteins in the saccharification medium, for both fungal secretomes and commercial cocktails (**Fig. 9**). This revealed that Pectinase, Cellulase Enzyme Blend (CEB), Cellulase from *Trichoderma reseei* (CTR), and Viscozyme® L achieved hydrolytic efficiencies intermediate between those of SmF- and SSF-derived fungal extracts. Glucose was the predominant monosaccharide released by these enzyme preparations. Since commercial cellulase cocktails are generally produced via submerged fermentation and subsequent concentration steps (Sigma-Aldrich, 2025a, 2025b, 2025c), the comparable performance between commercial preparations and our SmF-derived enzymes validates our experimental approach. More importantly, the higher efficiency observed with SSF-derived enzymes underscores the advantage of solid-state fermentation for producing potent lignocellulolytic enzyme systems.

**Fig. 9:**
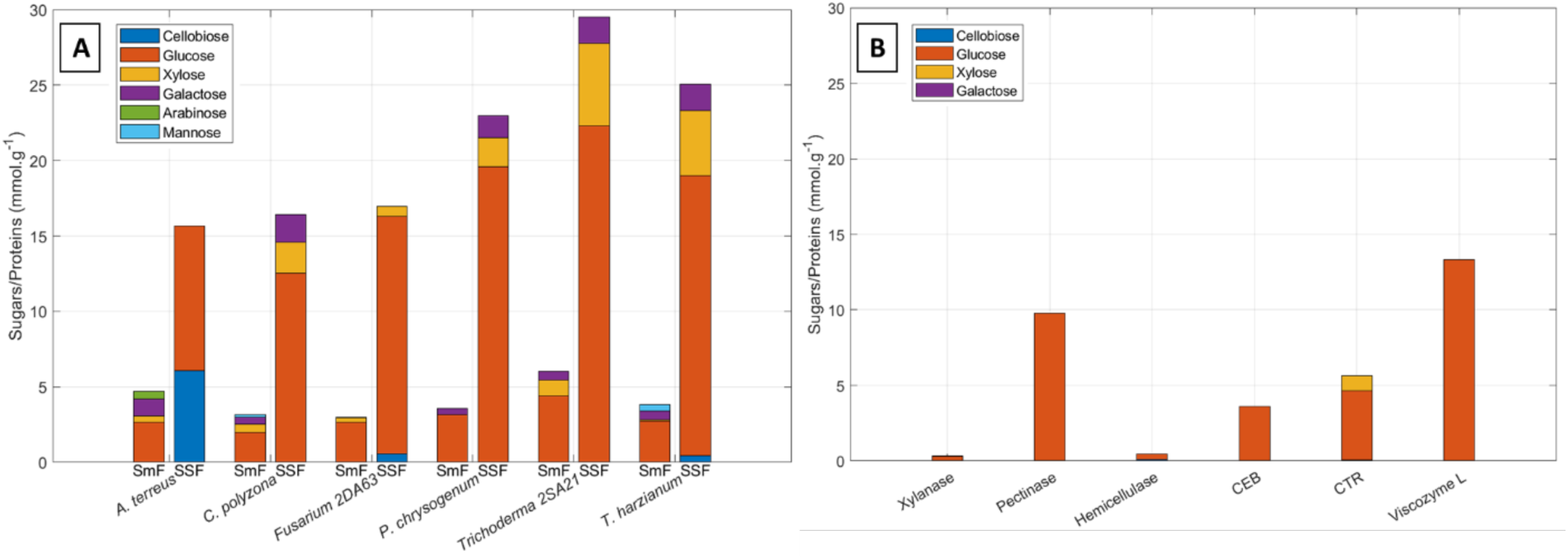
Comparison of sugar release efficiency between fungal extracts and commercial enzymes. (A) Sugar production normalized by total protein content (mmol·g⁻¹ protein) for six fungal strains cultivated under submerged fermentation (SmF) or solid-state fermentation (SSF) on flax shives. (B) Sugar production from commercial enzyme preparations under identical saccharification conditions. Sugars quantified by HPLC: cellobiose (blue), glucose (orange), xylose (yellow), galactose (purple), arabinose (green), and mannose (cyan). Saccharification was performed on flax shives (15 mg·mL⁻¹) at 30 °C for 48 h and 200 rpm. Sugars concentration measured before saccharification was subtracted.

In contrast, two commercial preparations showed limited activity on flax shives. Xylanase (Pentopan Mono BG®) is a single purified endo-β-(1→4)-xylanase (Sigma-Aldrich, 2025d) and lacks the accessory enzyme activities required for efficient hemicellulose degradation. Hemicellulase, despite containing multiple glycolytic activities (Sigma-Aldrich, 2025e), also exhibited poor performance, possibly due to suboptimal enzyme ratios for this specific lignocellulosic substrate.

### 3.3. DOE saccharification synergy

Clearly, through the present study, we demonstrated the ability of several fungi species to produce enzyme arsenals, sometimes complementary, enabling saccharification efficiencies similar to or superior to commercial cocktails. Nevertheless, at this stage, we rely on the ability of each strain to produce the complete range of enzymes adapted to the targeted biomass, which, as we have observed, can lead to the absence of crucial activities for the most complete possible depolymerization. This therefore naturally leads to the idea of mixing secretomes, particularly those whose enzymatic activities can potentially work in synergy, in order to obtain optimized cocktails. This somehow mimics how these strains work in nature, where fungi sometimes act sequentially, or even simultaneously, to degrade a given substrate (Srivastava *et al*., 2018; Janusz *et al*., 2017).

Nonetheless, the design of an efficient cocktail is challenging due to several physicochemical, structural, and compositional factors that limit the hydrolyze of these complex polysaccharides. Recent commercially enzyme cocktails demonstrated that enzymatic hydrolysis efficiency depends not only on cellulase activity but critically on the synergistic action of multiple enzyme classes including hemicellulases, pectinases, and lytic polysaccharide monooxygenases (Adsul *et al*., 2020). Kumar and Wyman, 2009 summarized that “synergism” was defined “as the ratio of the rate or yield of product release by enzymes when used at the same time to the sum of rate or yield of these products when the enzymes are used separately in the same amounts as they were employed in the mixture”. Different commercial enzymes cocktails or freshly prepared secretomes optimization were made empirically or with design of experiment (DOE) strategies with a specific enzyme supplementation, nonhydrolytic enzymes supplementation, or surfactants or other chemicals supplementation, but didn’t provided a clear understanding of what would made and efficient cocktail based on the measure of standard enzyme activities measured (soluble test substrates do not predict activity against natural fibers), as it very biomass and source-strain enzyme cocktail dependent (Berlin *et al*., 2007; Dąbkowska *et al*., 2017; Hassan *et al*., 2013; Kabel *et al*., 2006; Lopes *et al*., 2018). Saccharification assays are therefore more suitable to validate with the sugar release to assess actual efficiency (Raulo *et al*., 2021).

Consequently, to conclude our study, we sought to determine whether it was possible to combine the different secretomes generated in order to obtain an optimized cocktail for the depolymerization of our flax source. As enzyme extracts from SSF were more efficient in our case, when based on total protein content, all six secretomes obtained in this fermentation mode from the previously chosen fungi, were combined to create mixes of enzymatic cocktails and asses synergetic effect on untreated flax shives saccharification. For record, *A. terreus* produced glucose/cellobiose and cellobiose, and therefore provides substrate for β-glucosidase-rich fungi, potentially creating a productive metabolic cascade. *Coriolopsis polyzona* demonstrated diverse, balanced enzyme profile with high phenol oxidase for lignin modification, potentially improving substrate accessibility for other enzymes. *T. harzianum* and *Trichoderma* 2SA21 were already efficient saccharifiers but with different β-glucosidase profiles. *Fusarium* 2DA63 has revealed itself being pure endo-enzyme producer still generating glucose and xylose monomers, potentially synergizing with exo-enzyme producers and high lignin-modifying capacity with some laccase activity. *P. chrysogenum* had the highest total enzymatic activity with broad spectrum coverage.

In this objective, we selected the use of a Scheffé simplex-lattice mixture DOE to systematically evaluate synergistic interactions between secretomes/enzyme activities as this approach could model non-linear blending effects and identify optimal mixture compositions when individual components contribute to the overall response (Cornell, 2002). Mixture designs are particularly suited for biological association optimization, where the constraint that components must sum to 100% is inherent to the system (Piepel *et al*., 2002). Gao *et al*., (2010) found that increasing enzyme loading increase obviously the glucose yield but then it became less sensible to different enzymes ratio activities, meaning the interactions influence was lost. Therefore, the emphasis here was to study only blend interactions (Rispoli and Shah, 2007) so the design maintained constant total protein content levels across all mixture (0.04 g.L^-1^) while systematically exploring the entire experimental domain. The inclusion of all vertices (pure strains), edges (binary combinations), and higher-order interior points enables polynomial modeling of mixture responses and detection of both synergistic and antagonistic interactions between fungal strains (Scheffé, 1958).

As a result of this DOE, shown by **Fig. 10**, for pure strains, the same trend was observed as with the previous measurements (**Fig. 8**): *Trichoderma* 2SA21 exhibited the highest glucose yield (1.39 mM), followed by *T. harzianum* (1.21 mM) and *P. chrysogenum* (0.89 mM). In contrast, *A. terreus* demonstrated the lowest saccharification efficiency (0.22 mM glucose), while *C. polyzona* and *Fusarium* 2DA63 showed intermediate performance (0.74 and 0.71 mM, respectively). Glucose release was primarily biomass-dependent, indicating that individual strains contributed to production without critical interactions (R² adjusted = 0.69). However, as confirmed as only significant interactions beyond pure strain effects (Supplementary Data,1 **Table S5**), the combinations of T21 and PC (Mix-16) and T21 and F63 (Mix-17) exhibited highly significant synergy, yielding 1.74 and 1.56 mM of glucose, respectively - representing 52% and 49% improvements over the expected additive values from individual contributions.

**Fig. 10:**
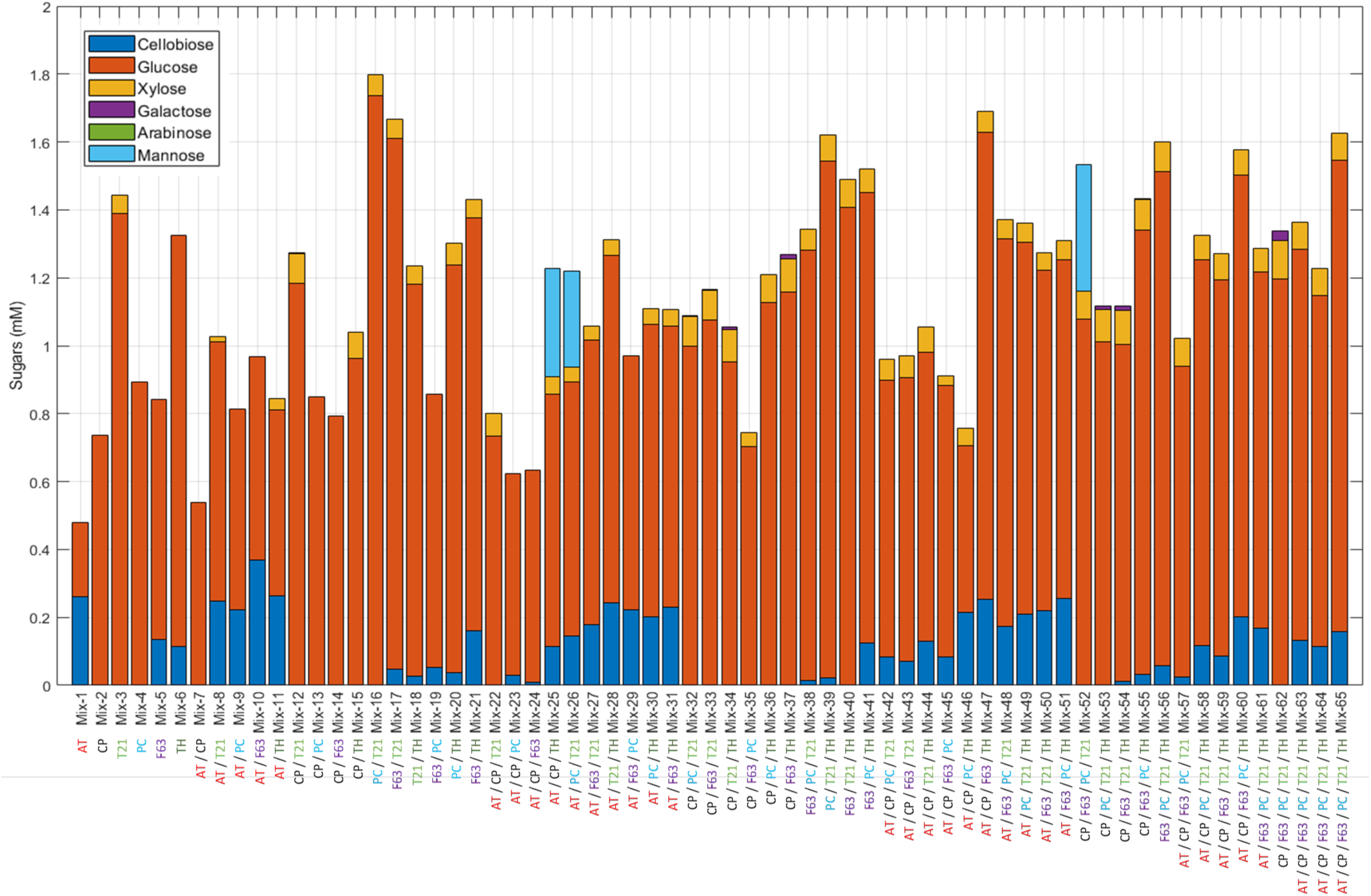
DOE mixture saccharification results from six fungi enzymatic extracts. Composition from Mix-1 to Mix-65 : 6 pure, 15 binary, 20 ternary, 15 quaternary, 6 quinary, triplicate of senary mixture, all normalized at 0.04 g.L^-1^ total proteins content before mixture. AT: *Aspergillus terreus*. CP: *Coriolopsis polyzona*. T21: *Trichoderma* 2SA21. PC: *Penicillium chrysogenum*. F63: *Fusarium* 2DA63. TH: *Trichoderma harzianum*.

The same strains also showed cellobiose accumulation with *A. terreus*, *Fusarium* 2DA63, and *T. harzianum*, as confirmed by the model with significant coefficient estimates and high synergy for the ternary interaction AT/CP/TH (Supplementary Data 1, **Tables S3** and **S4**), suggesting this is an unfavorable combination for monosaccharide production. However, the AT and CP combination showed antagonism (β = -0.64), indicating that in Mix-7, CP β-glucosidase activity enhanced cellobiose hydrolysis, resulting in 13% more glucose. A similar pattern was observed with Mix-17 and Mix-37, where cellobiose accumulation was expected, but PC addition led to 10 and 31% more glucose, respectively. Interestingly, Mix-40 showed no cellobiose accumulation despite containing TH and F63, which would typically promote it; the action of T21 enzymes appeared to counteract this effect, resulting in 28% more glucose release.

Xylose, which was generally present with all strains, was detected here only for *Trichoderma* 2SA21. This could be attributed to the dilution factor (two- to three-fold), reducing hydrolysis efficiency, as *Trichoderma* 2SA21 showed the highest xylose content (**Fig. 8**). The special cubic model explained the global variance well (R² adjusted = 0.97); however, with a lack-of-fit (LoF) p-value of 0.027, it was not optimally suited (Supplementary Data 1, **Table S3**). A more complex fitting approach was tried (data not shown), namely the full cubic model with 15 additional parameters accounting for asymmetric binary interactions, but was unbalanced and yielded a worse LoF p-value. Overall, it appeared that any combination including *T. harzianum* or *Trichoderma* 2SA21 led to xylose release.

Galactose was absent; ternary interactions were required to detect it, but only in trace amounts, precluding reliable model fitting. Nevertheless, slight synergy was observed between mixed-strain enzymes: although the most efficient pure extract was diluted, complementation with less efficient strains led to a more diverse sugar profile. A notable finding was the appearance of mannose in the ternary mixtures AT/CP/TH and AT/PC/T21, as well as in the quaternary mixture CP/F63/PC/T21. TH, T21, PC, and F63 indeed possess α-1,2-mannosidase and/or mannan endo-1,4-β-mannosidase activity; however, their pure enzymatic extracts alone could not release mannose. The addition of CP and AT appeared to enable more extensive hemicellulose hydrolysis. Due to insufficient mannose data, model fitting was not possible.

As none of the models were sufficiently robust to predict sugar release, the efficiency of interactions was assessed using simple relative deviation (percentage) from the expected sum of individual monosaccharides released compared to the observed release (Khamassi and Dumon, 2023). Cellobiose was not accounted for in synergistic effects, as its presence indicates a lack of β-glucosidase activity, and monosaccharides are the target products for biorefinery applications (Wang *et al*., 2024).

Regarding the six-strain blend replicates, high variability was observed (46 ± 17% synergy); therefore, a significant interaction above zero effect would require more than 17% deviation. As seen in **Fig. 11**, binary combinations exhibited significant synergistic interactions, with 5 out of 15 combinations (Mix-10, Mix-16, Mix-17, Mix-20, and Mix-21) showing 21–54% more sugar release than expected, while Mix-18 (T21/TH) showed an antagonistic effect with 18% less sugar release than expected.

**Fig. 11:**
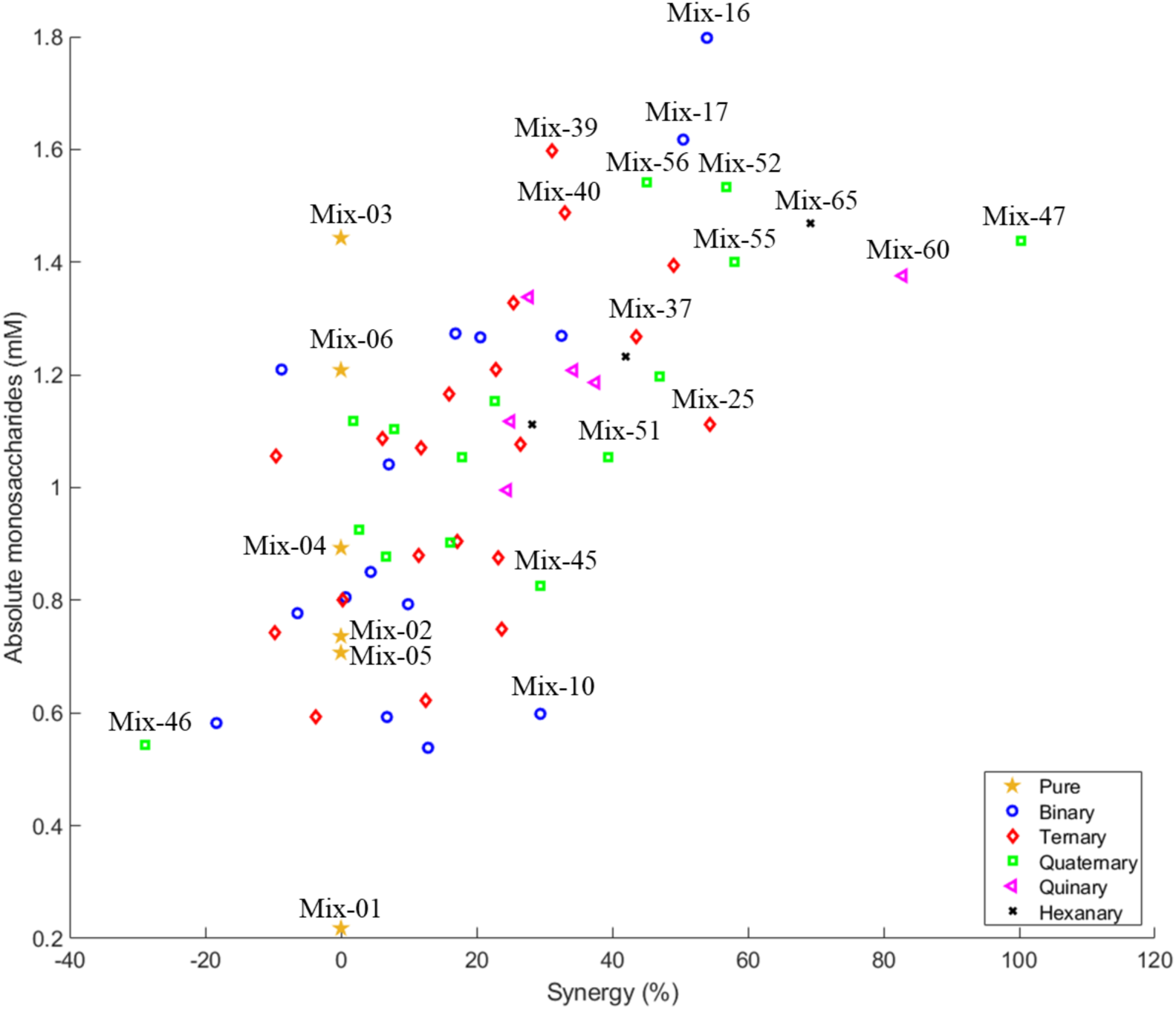
Relationship between synergistic effect and absolute monosaccharides (glucose, xylose, galactose, mannose) production across 65 fungal mixture combinations. Each point represents a distinct mixture composition, color-coded by complexity: pure (yellow star), binary (15 blue circle), ternary (20 red diamonds), quaternary (15 green squares), quinary (6 magenta triangles), and hexanary (black crosses, triplicate). Synergy was calculated as the percentage deviation from expected pure strain additive performance. Notable mixtures are labeled above the mark.

Ternary combinations showed synergistic interactions in half of the blends, with sugar release enhancement ranging from 23 to 54%. Mix-25 (AT/CP/TH) and Mix-41 (PC/F63/TH) gave 54% and 49% more sugar than expected, respectively.

Quaternary blends presented both the worst and best effects: Mix-47 (AT/CP/F63/TH) yielded twice the expected sugar quantity, corresponding to 100% synergy, while Mix-46 (AT/CP/PC/TH) showed 29% less sugar than expected—the only difference being the replacement of F63 with PC. Four other quaternary mixes showed more than 40% synergistic effect: Mix-48 (AT/T21/PC/F63) (47%), Mix-52 (CP/T21/PC/F63) (57%), Mix-55 (CP/PC/F63/TH) (58%), and Mix-56 (T21/PC/F63/TH) (45%).

Quinary blends (Mix-57–62) were exclusively and significantly synergistic, with 25–83% more sugar than expected, the highest being Mix-60 (AT/CP/PC/F63/TH). Generally, a clear trend toward increased synergy was observed as the number of enzymes from different strains increased, possibly due to dilution of inhibitor-sensitive enzymes or addition of highly active enzymes, complementary hemicellulases, or auxiliary activity enzymes, although no clear pattern emerged.

However, from a bioprocess perspective, the choice of the optimal blend is driven by high yield of the desired product, its purity, and simplicity of implementation to minimize costs (Pothakos *et al*., 2018). Therefore, even with clear synergistic effects, complex blends did not outperform Mix-16, which had a 54% synergistic effect and the highest sugar concentration (1.8 mM) with exclusively monosaccharides.

As a final note, it should be noted that Mafa *et al*., (2021) summarized studies on enzyme synergism and revealed cases where low synergy still resulted in higher soluble sugar release than with individual enzymes. This was explained by (i) uneven substrate surfaces generated by endoglucanases, which could inhibit processive cellobiohydrolases, (ii) competition between enzymes for the same substrate, (iii) biomass complexity with less accessible cellulose, and (iv) rate-limiting endoglucanase releasing dextrans slowly while cellobiohydrolase II (exo-glucanase) acts quickly to release soluble oligomers. This means that for future development and strain screening, selecting a few of the best-performing pure strains - which possess a more suitable enzymatic arsenal for a given type of biomass - is preferable to searching for individual enzyme complementarity.

## 4. Conclusion

This comprehensive study systematically evaluated the enzymatic potential of 19 fungal strains under SSF conditions, followed by detailed comparative analysis of 6 selected strains across both SSF and SmF fermentation modes, demonstrating that fermentation strategy profoundly influences enzyme production efficiency and saccharification performance on untreated flax shives. SSF exhibited higher endo-xylanase and cellulase activities (two-fold on average compared to SmF), whereas SmF showed drastically elevated PPO activity. Proteomics analysis indicated lower protein diversity in SSF but a higher proportion of enzymes for polysaccharide metabolism. Although SmF produced 5-fold more total protein than SSF, sugar yields were comparable when normalized to equivalent aqueous extraction volumes. Consequently, SSF-derived enzymes exhibited superior mass-specific efficiency, based on sugar production, confirming the principle that the natural habitat-mimicking conditions of solid-state cultivation favor the production of highly active lignocellulolytic enzymes, as previously reported for various fungal systems (Leadbeater *et al*., 2025; Sankar *et al*., 2023). As a final attempt to further improve the enzyme cocktails for flax depolymerization, a Scheffé mixture DOE revealed that strategic enzyme cocktail formulation through simple binary combinations can achieve significant synergistic effects. But higher synergy values did not necessarily correlate with higher absolute sugar yields, highlighting the importance of selecting high-performing individual strains before optimizing their combinations. Although, those results should be confirmed at higher enzymatic production and saccharification loading, as industrial applications typically require high-solids processes (15-25% w/v) where the “high-solids effect” significantly reduces saccharification efficiency due to mass transfer limitations and product inhibition (Angeltveit *et al*., 2024).

## Supporting information

Supplementary Data 2 - Mass Protein Identification

Supplementary Data 3 - Codes

## CREDIT AUTHORSHIP CONTRIBUTION STATEMENT

All authors contributed jointly to all the aspects of the work reported in the manuscript. NK, BD, SH, QH, VP and EH planned the experimental work. NK, BD, SH and QH carried out the experimental work. NK, BD, VP and EH contributed to the interpretation of the results and writing of the manuscript. VP and EH contributed to the supervision and critical review. All authors read and approved the final manuscript.

## DECLARATION OF COMPETING INTEREST

The authors declare that they have no known competing financial interests or personal relationships that could have appeared to influence the work reported in this paper.

## ACKNOWLEDGEMENTS

This work was supported by the PEECFUEL project that is financed by the French National Research Agency (ANR), with the contractual reference ‘ANR-23-CE05-0026’. The REALCAT platform is benefiting from a Governmental subvention administrated by the ANR within the frame of the ‘Future Investments’ program (PIA), with the contractual reference ‘ANR-11-EQPX0037’.

## SUPPLEMENTARY DATA

Supplementary Data 1: Strains information, statistical analysis, and proteins classification from LC-MS data.

Supplementary Data 2: LC-MS protein identification.

Supplementary Data 3: Codes used for robot programming, automated data processing, and statistical analyses.

## DATA AVAILABILITY

All the data required to reproduce this work are available in the Supplementary Data provided with the article

## DECLARATION OF GENERATIVE AI AND AI-ASSISTED TECHNOLOGIES IN THE MANUSCRIPT PREPARATION PROCESS

During the preparation of this work the author used Claude (Anthropic, Claude Opus 4.5) in order to assist with debugging code used for data analysis and statistical modeling. After using this tool/service, the author reviewed and edited the content as needed and take full responsibility for the content of the published article.

## ABBREVIATIONS

AA: auxiliary activity
CAZyme: carbohydrate-active enzyme
CBH: cellobiohydrolase
CBM: carbohydrate-binding module
CE: carbohydrate esterase
CWDE: cell wall degrading enzymes
EG: endo-glucanase
GH: glycoside hydrolase
GT: glycosyltransferase
LC-MS/MS: liquid chromatography-tandem mass spectrometry
LPMO: lytic polysaccharide monooxygenase
PDA: potato dextrose agar
PL: polysaccharide lyase
SmF: submerged fermentation
SSF: solid-state fermentation.

## Notes

### Competing Interest Statement

The authors have declared no competing interest.

