## Supplementary Data 1 for "Comparative Secretome Analysis and Enzyme Cocktail Optimization of Six Fungal Species Under Solid-State and Submerged Fermentation for Lignocellulosic Saccharification of Flax Shives"

#### 1. Strains information

Table S1: Fungi characteristics

| **Division** | **Name** | **Lignocellulose degradation group** | **Lifestyle** |
|---|---|---|---|
| **Ascomycota** | *A. oryzae* | Soft-rot | Saprotroph |
| **Ascomycota** | *A. terreus* | Soft-rot | Saprotroph/Opportunistic endophyte |
| **Basidiomycota** | *C. polyzona* | White-rot | Wood saprotroph |
| **Ascomycota** | *Fusarium 2DA63* | Soft-rot | Saprotroph/Pathogen |
| **Ascomycota** | *F. graminearum* | Soft-rot | Saprotroph/Pathogen |
| **Mucoromycota** | *M. circinelloides* | Soft-rot | Common saprotroph/Opportunistic parasite |
| **Mucoromycota** | *M. fragilis* | Soft-rot | Common saprotroph/Opportunistic parasite |
| **Mucoromycota** | *Mucor sp.* | Soft-rot | Common saprotroph/Opportunistic parasite |
| **Ascomycota** | *Penicillium 2DA2* | Soft-rot | Common saprotroph/Opportunistic parasite |
| **Ascomycota** | *P. chrysogenum* | Soft-rot | Common saprotroph/Opportunistic parasite |

| | | | |
|---|---|---|---|
| **Ascomycota** | *Paecilomyces 11SE* | Soft-rot | Common saprotroph |
| **Ascomycota** | *Paecilomyces 11SA2* | Soft-rot | Common saprotroph |
| **Ascomycota** | *Trichoderma 2DA61* | Soft-rot | Mycoparasite/saprotroph |
| **Ascomycota** | *Trichoderma 2DA62* | Soft-rot | Mycoparasite/saprotroph |
| **Ascomycota** | *Trichoderma 2DA67* | Soft-rot | Mycoparasite/saprotroph |
| **Ascomycota** | *Trichoderma 2SA21* | Soft-rot | Mycoparasite/saprotroph |
| **Ascomycota** | *T. harzianum* | Soft-rot | Mycoparasite/saprotroph |
| **Ascomycota** | *Talaromyces 47* | Soft-rot | Saprotroph/Opportunistic endophyte/Mycoparasite |
| **Basidiomycota** | *Trametes versicolor* | White-rot | Wood saprotroph |

Table S2: Protein quantification with Braford assay. Average of two sampling. Triplicate OD reading per sampling.

| **Fungi** | **Total proteins (g/L)** | **Standard error** |
|---|---|---|
| ***A. oryzae*** | 0.08 | 0.00 |
| ***A. terreus*** | 0.05 | 0.00 |
| ***C. polyzona*** | 0.09 | 0.00 |
| ***Fusarium 2DA63*** | 0.08 | 0.01 |
| ***F. graminearum*** | 0.06 | 0.02 |
| ***M. circinelloides*** | 0.08 | 0.01 |
| ***M. fragilis*** | 0.06 | 0.02 |
| ***Mucor sp.*** | 0.09 | 0.01 |
| ***Penicillium 2DA2*** | 0.11 | 0.01 |
| ***P. chrysogenum*** | 0.08 | 0.01 |
| ***Paecilomyces 11SE*** | 0.06 | 0.00 |
| ***Paecilomyces 11SA2*** | 0.08 | 0.00 |

| | | |
|---|---|---|
| ***Trichoderma 2DA61*** | 0.08 | 0.01 |
| ***Trichoderma 2DA62*** | 0.06 | 0.03 |
| ***Trichoderma 2DA67*** | 0.09 | 0.01 |
| ***Trichoderma 2SA21*** | 0.09 | 0.01 |
| ***T. harzianum*** | 0.09 | 0.01 |
| ***Taloromyces 47*** | 0.09 | 0.01 |
| ***Trametes versicolor*** | 0.09 | 0.02 |

### 2. Statistical analysis

All statistical analyses were performed using MATLAB R2022b (MathWorks, Natick, MA, USA).

#### *2.1. Correlation matrix among enzyme activities and released sugars (SSF)*

To assess the relationships between the concentrations of released sugars and the enzymatic activities measured for each fungal extract (SSF), a correlation analysis was performed. Sugar concentrations (cellobiose, glucose, xylose, and galactose) and enzymatic activities (endo-xylanase, endo-cellulase, β-xylosidase, β-glucosidase, and phenol oxidase) were standardized (z-score normalization) prior to computation. Pairwise Pearson correlation coefficients were calculated across all samples, and the resulting correlation matrix was visualized as a heatmap (Somadder et al. 2025).

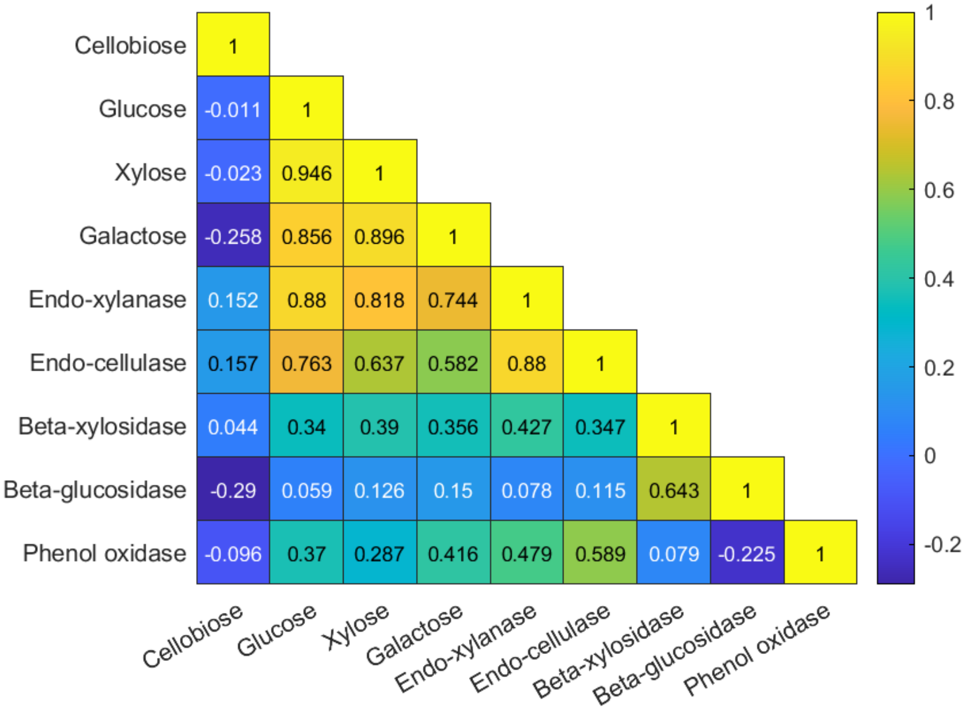


Fig. S1: Correlation matrix between sugar concentrations and enzyme activities across 19 enzyme extracts from fungi grown in SSF. Yellow: strong positive correlation; blue: weak or negative correlation.

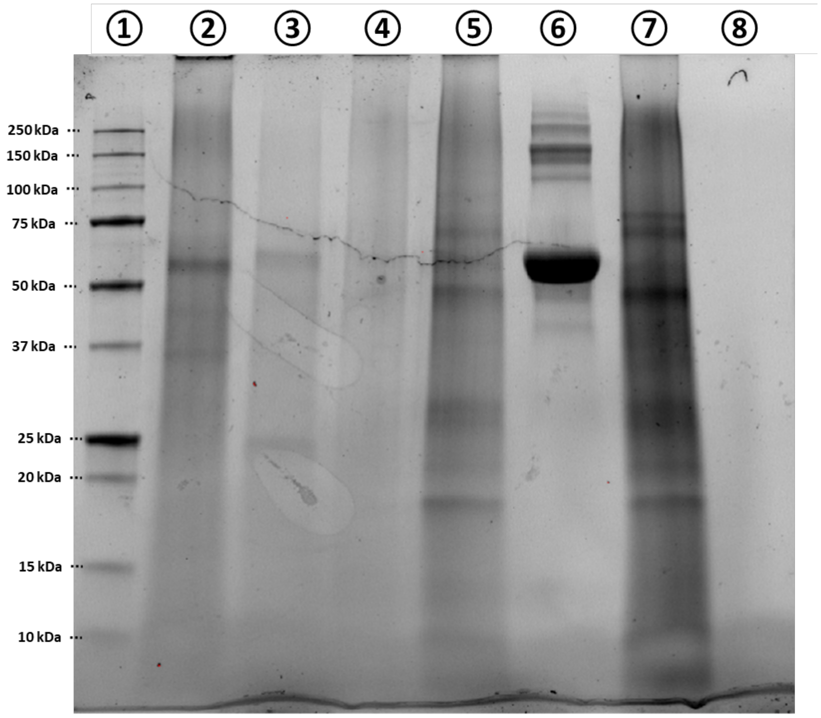


Fig. S2: SDS-PAGE of concentrated proteins extracts from SSF, stain-free imaging. ① Mark Ladder ② P. chrysogenum ③ Fusarium 2DA63 ④ A. terreus ⑤ Trichoderma 2SA21 ⑥ BSA ⑦ Trichoderma harzianum ⑧ Negative control.

#### 2.2. *SmF vs SSF enzymatic activity comparison*

Data from each enzymatic activity assay were analyzed separately with three technical replicates per condition. The normal distribution of residuals was assessed using the Shapiro-Wilk test (sample sizes below 50). Homogeneity of variances was assumed given the balanced experimental design (n = 3 per group). A two-way analysis of variance (ANOVA) was conducted to evaluate the main effects of fermentation type (SmF vs. SSF) and fungal strain (six levels), as well as their interaction on enzymes activity. Pairwise comparisons were performed using the Tukey honestly significant difference (HSD) test. Compact letter displays were generated to visualize homogeneous subsets, where treatments sharing the same letter are not significantly different ($\alpha = 0.05$).

An additional one-way ANOVA was conducted treating each fungal strain × fermentation type combination as a separate level (12 groups total), allowing direct comparison of all treatment combinations. This approach enabled the identification of specific fungus-fermentation pairings with superior enzymatic activity while accounting for potential interactions. Tukey HSD post-hoc tests with compact letter displays were used to classify combinations into statistically homogeneous groups.

All statistical tests were two-tailed with a significance threshold of $\alpha = 0.05$. Results are presented as mean ± standard error (SE). All figures were generated using MATLAB, and statistical tables were exported to Microsoft Excel for documentation.

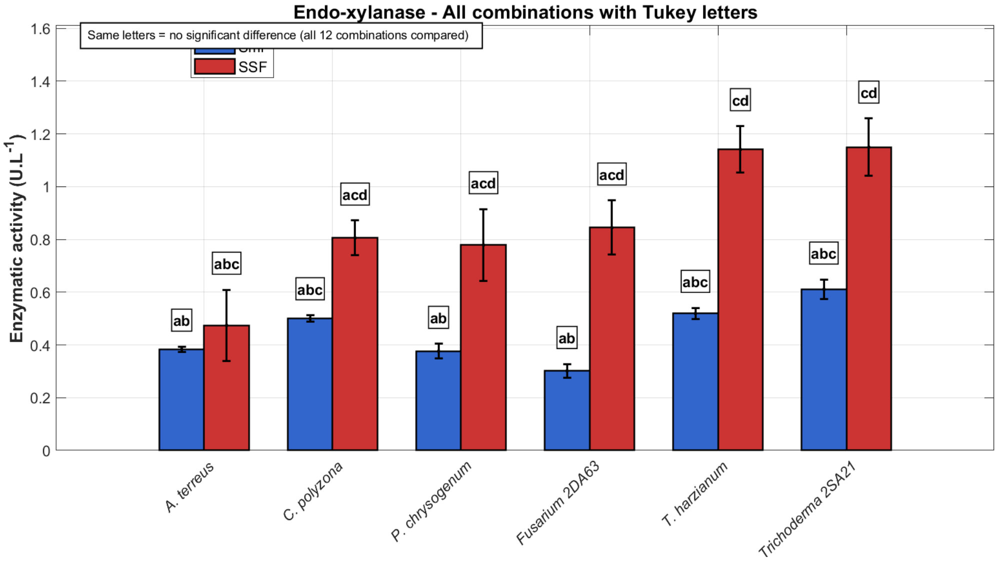


Fig. S3: Endo-Xylanase activity produced by six fungal strains under submerged fermentation (SmF) and solid-state fermentation (SSF) conditions. Enzymatic activity ($U \cdot L^{-1}$) was measured after growth on flax biomass (2g, 7 days, pooled triplicate extraction and 10 kDa concentration). Blue bars represent SmF; red bars represent SSF. Error bars indicate standard deviation (n = 3). Statistical analysis was performed using two-way ANOVA followed by Tukey's HSD post-hoc test for all 12 strains × fermentation mode combinations. Bars sharing the same letter are not significantly different ($p > 0.05$).

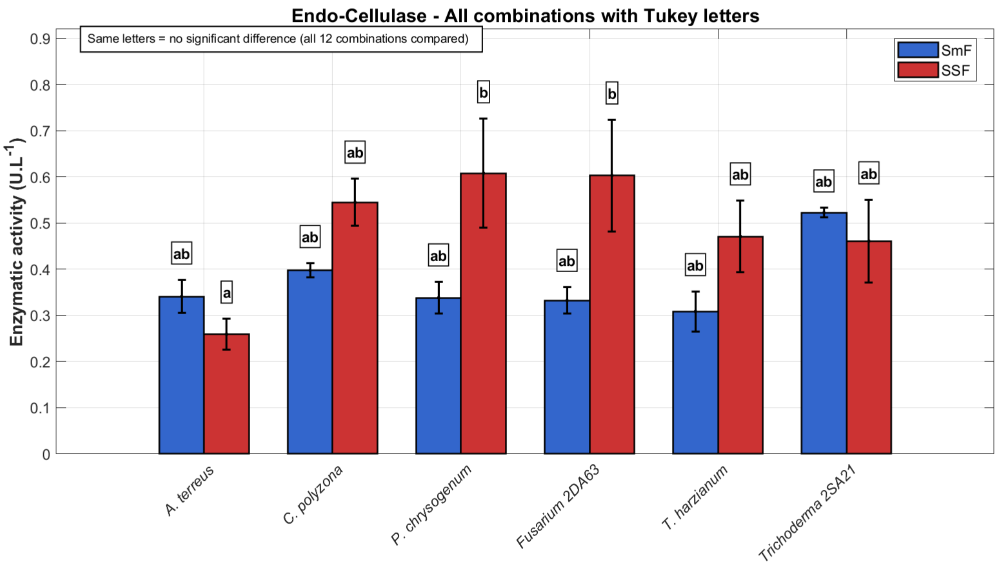


Fig. S4: Endo-Cellulase activity produced by six fungal strains under submerged fermentation (SmF) and solid-state fermentation (SSF) conditions. Enzymatic activity ($U \cdot L^{-1}$) was measured after growth on flax biomass (2g, 7 days, pooled triplicate extraction and 10 kDa concentration). Blue bars represent SmF; red bars represent SSF. Error bars indicate standard deviation (n = 3). Statistical analysis was performed using two-way ANOVA followed by Tukey's HSD post-hoc test for all 12 strains × fermentation mode combinations. Bars sharing the same letter are not significantly different ($p > 0.05$).

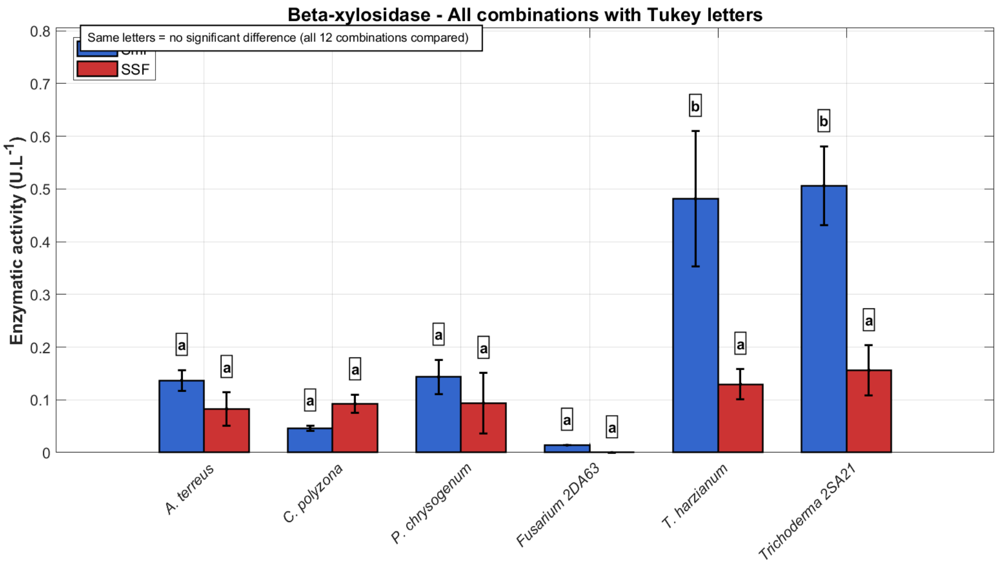


Fig. S5: β-Xylosidase activity produced by six fungal strains under submerged fermentation (SmF) and solid-state fermentation (SSF) conditions. Enzymatic activity ($U \cdot L^{-1}$) was measured after growth on flax biomass (2g, 7 days, pooled triplicate extraction and 10 kDa concentration). Blue bars represent SmF; red bars represent SSF. Error bars indicate standard deviation (n = 3). Statistical analysis was performed using two-way ANOVA followed by Tukey's HSD post-hoc test for all 12 strains × fermentation mode combinations. Bars sharing the same letter are not significantly different ($p > 0.05$).

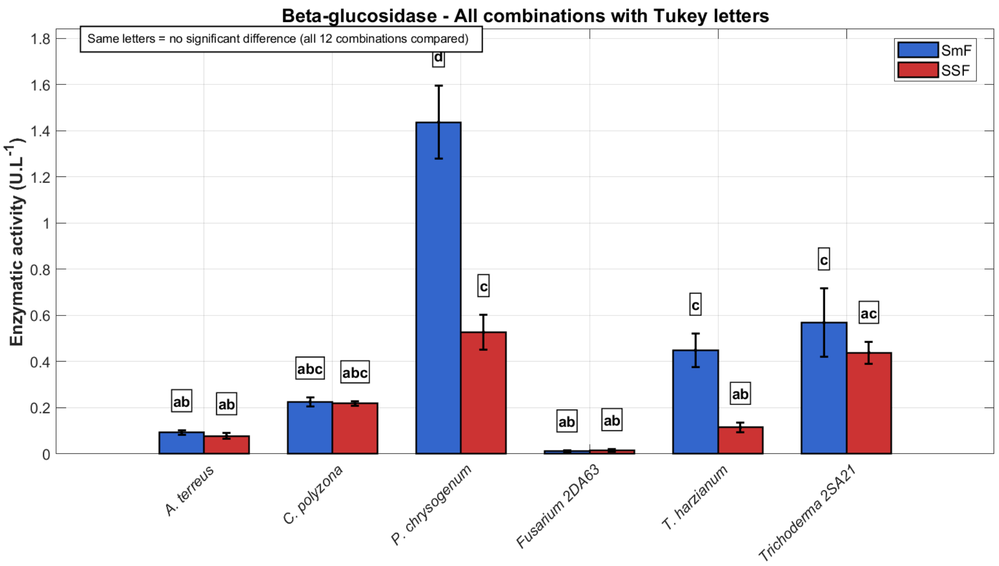


Fig. S6: β-Glucosidase activity produced by six fungal strains under submerged fermentation (SmF) and solid-state fermentation (SSF) conditions. Enzymatic activity (U·L⁻¹) was measured after growth on flax biomass (2g, 7 days, pooled triplicate extraction and 10 kDa concentration). Blue bars represent SmF; red bars represent SSF. Error bars indicate standard deviation (n = 3). Statistical analysis was performed using two-way ANOVA followed by Tukey's HSD post-hoc test for all 12 strains × fermentation mode combinations. Bars sharing the same letter are not significantly different ($p > 0.05$).

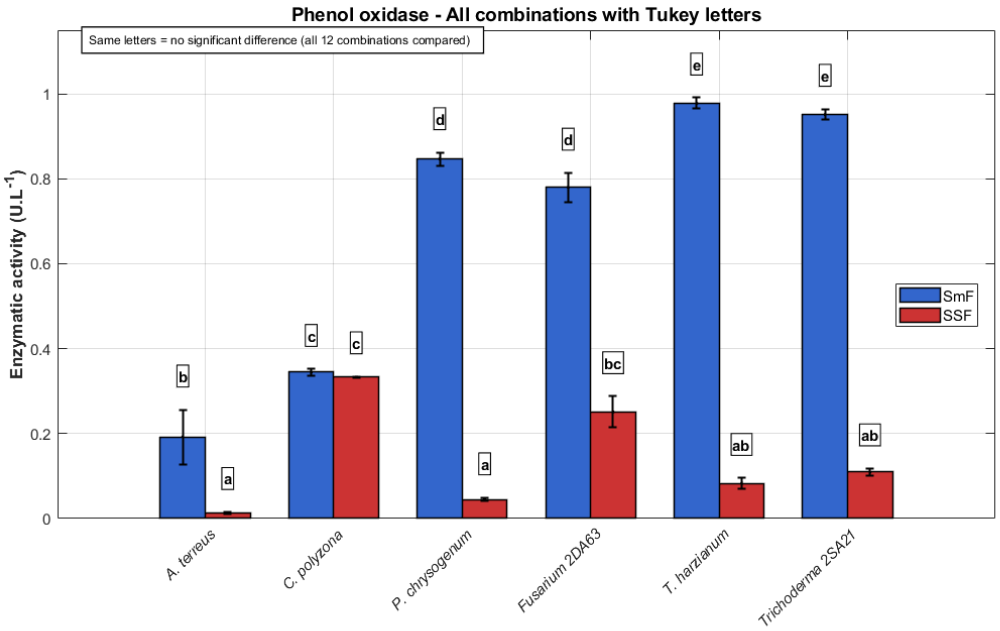


Fig. S7: Phenol oxidase activity produced by six fungal strains under submerged fermentation (SmF) and solid-state fermentation (SSF) conditions. Enzymatic activity (U·L⁻¹) was measured after growth on flax biomass (2g, 7 days, pooled triplicate extraction and 10 kDa concentration). Blue bars represent SmF; red bars represent SSF. Error bars indicate standard deviation (n = 3). Statistical analysis was performed using two-way ANOVA followed by Tukey's HSD post-hoc test for all 12 strains × fermentation mode combinations. Bars sharing the same letter are not significantly different ($p > 0.05$).

#### 2.3. *DOE sugar release and strain dependance*

Data were analyzed using Scheffé special cubic polynomial mixture (24 degree of freedom with 65 experimental data) models implemented in MATLAB (R2022b). The model included six linear blend terms ($\beta_i$) representing pure strain effects, 15 binary interaction terms ($\beta_{ij}$) quantifying synergistic or antagonistic interactions, and 20 ternary interaction terms ($\beta_{ijk}$) (Piepel et al. 2002):

$$Y = \sum_{i=1}^{q} \beta_i x_i + \sum_{i<j}^{q-1} \sum_{j<k}^{q} \beta_{ij} x_i x_j + \sum_{i<j}^{q-2} \sum_{j<k}^{q-1} \sum_{k}^{q} \beta_{ijk} x_i x_j x_k$$

where Y is the total sugar concentration (mM), x_i the proportion of strain i, and β_ij the interaction coefficient between strains i and j (i<j), and so on, q the number of strains. Model adequacy was assessed using $R^2$, adjusted $R^2$, RMSE, and lack-of-fit tests. Statistical significance of individual coefficients was evaluated using t-tests with $\alpha = 0.05$. For each sugar the model fit is presented in the following tables :

Table S3: Summary of statistical parameters for the polynomial models fitted to sugar production data using the Scheffé mixture design.

| **Polymer** | **$R^2$** | **adjusted $R^2$** | **RMSE** | **Lack-of-fit** |
|---|---|---|---|---|
| **Cellobiose** | 0.935 | 0.828 | 0.041 | 0.224 |
| **Glucose** | 1 | 0.688 | 0.163 | 0.693 |
| **Xylose** | 1 | 0.969 | 0.006 | 0.027 |
| **Galactose** | 1 | -0.167 | 0.005 | 0.000 |
| **Mannose** | 1 | -0.227 | 0.077 | 0.000 |

Table S4: Coefficient estimates (β) and p-values of significant terms for the cellobiose special cubic model.

| **Fungus** | **Interaction coefficient estimate (β)** | **pValue** |
|---|---|---|
| **AT** | 0.27 | 9.1E-07 |
| **F63** | 0.14 | 2.6E-03 |
| **AT_CP** | -0.64 | 3.1E-03 |
| **AT_CP_TH** | 3.71 | 3.4E-03 |
| **AT_F63** | 0.58 | 6.2E-03 |
| **TH** | 0.12 | 7.4E-03 |
| **AT_T21** | 0.43 | 3.6E-02 |

Table S5: Coefficient estimates (β) and p-values of significant terms for the glucose special cubic model.

| **Fungus** | **Interaction coefficient estimate (β)** | **pValue** |
|---|---|---|
| **T21** | 1.39 | 9.7E-09 |
| **TH** | 1.21 | 1.1E-07 |
| **PC** | 0.90 | 1.1E-05 |
| **CP** | 0.75 | 1.2E-04 |
| **F63** | 0.72 | 1.9E-04 |
| **T21_PC** | 2.13 | 1.2E-02 |
| **T21_F63** | 1.98 | 1.8E-02 |

Table S6: Coefficient estimates (β) and p-values of significant terms for the xylose special cubic model.

| **Fungus** | **Interaction coefficient estimate (β)** | **pValue** |
|---|---|---|
| **CP_TH** | 0.32 | 2.82E-11 |
| **T21** | 0.05 | 1.80E-09 |
| **PC_TH** | 0.26 | 2.52E-09 |
| **CP_T21** | 0.25 | 2.90E-09 |
| **F63_TH** | 0.22 | 3.98E-08 |
| **CP_PC_F63** | 1.10 | 5.96E-07 |
| **CP_F63_TH** | 0.94 | 5.90E-06 |
| **AT_TH** | 0.15 | 1.70E-05 |
| **T21_PC** | 0.14 | 3.11E-05 |
| **AT_CP_T21** | 0.74 | 1.41E-04 |
| **T21_TH** | 0.13 | 1.45E-04 |
| **T21_F63** | 0.12 | 2.06E-04 |
| **CP_T21_F63** | 0.70 | 2.44E-04 |
| **CP_T21_PC** | 0.63 | 7.35E-04 |
| **PC_F63_TH** | 0.50 | 5.44E-03 |
| **T21_PC_F63** | 0.47 | 8.33E-03 |
| **AT_T21_PC** | 0.45 | 1.02E-02 |
| **CP_PC_TH** | 0.45 | 1.07E-02 |
| **AT_T21_F63** | 0.44 | 1.23E-02 |

### 3. Proteins classification from LC-MS data

Table S7: Proteins biological functions found in different four “main identified metabolisms” from UniProt database across six fungi in SmF and SSF mode

| Fungi biological metabolism | Pathogenesis | Polysaccharides metabolism | Stress |
|---|---|---|---|
| - Assimilation of proteinaceous substrates<br>- Biosynthesis of pyriculol and pyriculariol<br>- Carbohydrate metabolic process<br>- Carboxylic ester hydrolase activity<br>- Cell elongation<br>- Cell wall constituent<br>- Cell wall organization<br>- Cytoplasmic translational elongation<br>- DNA catabolic process<br>- Energy source<br>- Fungal-type cell wall organization<br>- Polysaccharides catabolic process<br>- Glucose metabolic process<br>- Glycerol and lipid metabolic process<br>- Glycosaminoglycan metabolic process<br>- Lipid metabolic process<br>- Malate metabolic process<br>- Nicotine catabolic process<br>- Nitrogen metabolism<br>- Peptidoglycan metabolic process<br>- Regulate important biological functions (Proteolysis)<br>- RNA endonuclease activity<br>- Spore formation | - Carbohydrate metabolic process<br>- Cell wall organization<br>- Colonization<br>- Glucosylceramide catabolic process<br>- Lipid catabolic process<br>- Lipid metabolic process<br>- Lysophospholipase activity<br>- Mycoparasitism<br>- Plant immunity suppressor<br>- Plant necrosis<br>- Proteolysis<br>- Virulence<br>- Phosphatidylinositol dephosphorylation<br>- Phospholipid metabolic process<br>- Protein turnover<br>- Signal<br>- Small secreted protein | - Arabinan catabolic process<br>- L-Arabinose metabolic process<br>- Arabinose cleavage in hemicellulose<br>- Assist lignocellulosic enzymes in biomass degradation<br>- Carbohydrate metabolic process<br>- Cellulose catabolic process<br>- Cell wall organization<br>- Cell wall pectin catabolic process<br>- Enhance Cellulases activities<br>- Enhance lignocellulose saccharification<br>- Galactooligosaccharides hydrolysis<br>- Galactose metabolic process<br>- Glucan catabolic process<br>- Glucan hydrolysis<br>- Glucose metabolic process<br>- Glycogen process<br>- Glycoprotein process<br>- Pectate & Galacturonans hydrolysis<br>- Pectin catabolic process<br>- Polygalacturonase activity<br>- Polysaccharides catabolic process<br>- Lignin degradation<br>- Mannan catabolic process<br>- Uronic acid polysaccharides degradation<br>- Xylan catabolic process<br>- Xyloglucan catabolic process<br>- Xyloglucan metabolic process | - Aldehyde metabolic process<br>- Antiviral<br>- Fungi defense response<br>- Cellular response to oxidative stress<br>- Defense system<br>- Ceramide & sphingosine & fatty acid process<br>- Methionine biosynthetic process<br>- Metabolic stress<br>- Monocarboxylic acid biosynthetic process<br>- Oxidoreductase activity<br>- Pentose-phosphate shunt<br>- Phosphate-containing compound metabolic process<br>- Phosphate Stress response<br>- Protein biosynthesis<br>- Traductionel stress response<br>- Protein folding<br>- Response to endoplasmic reticulum stress<br>- Response to oxidative stress and hydrogen peroxide catabolic process<br>- Response to phosphate absence<br>- Response to reactive species and hydrogen peroxide<br>- Lignin catabolic process<br>- Sphingomyelin catabolic process<br>- Environmental stress adaptation<br>- Immunological stress<br>- Superoxide metabolic process<br>Oxidative stress |

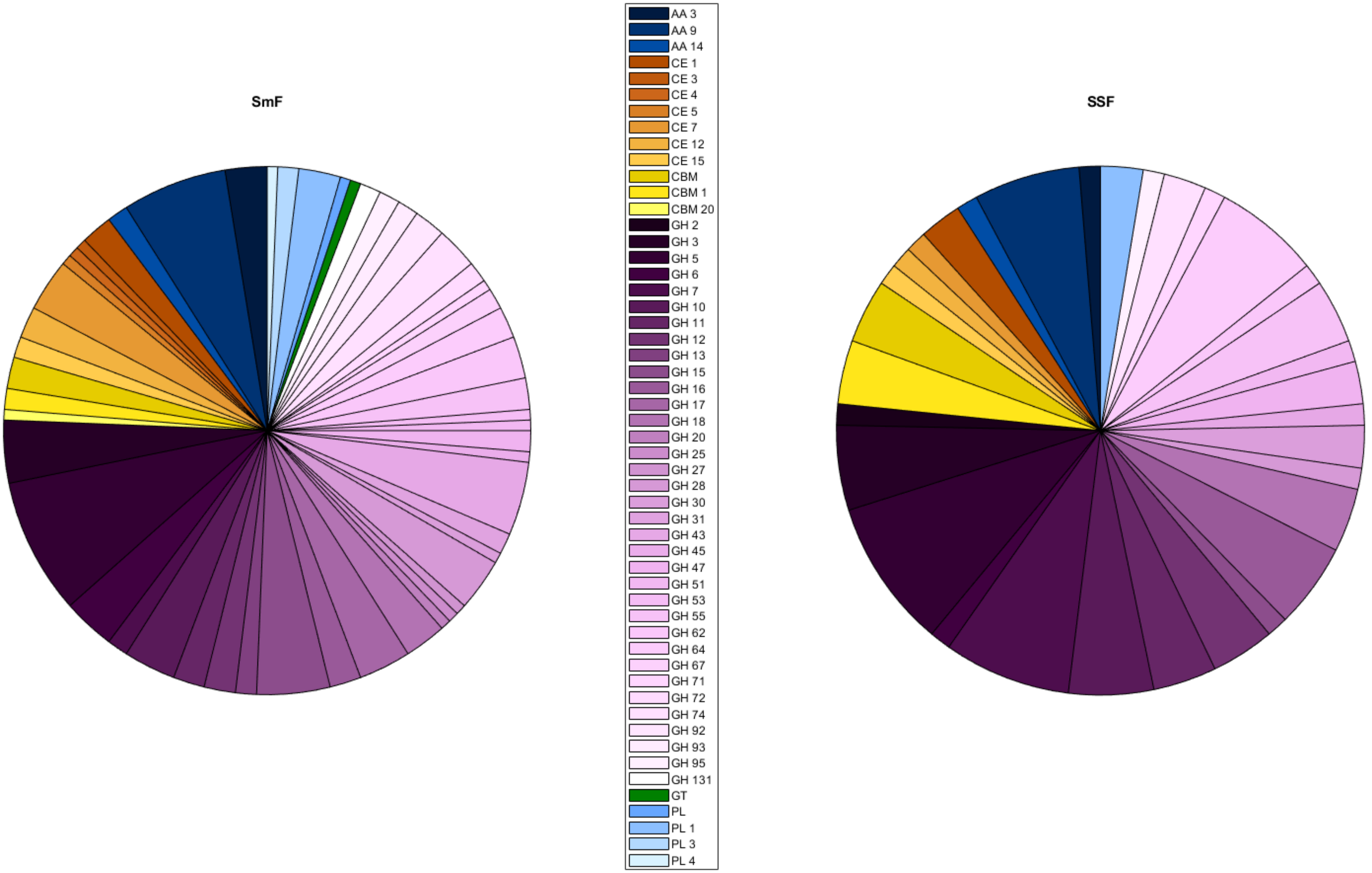


Fig. S8: Global CAZyme families distribution (percentage base on unique proteins) in SmF and SSF all fungi combined. AA: Auxiliary Activities. CE: Carbohydrate Esterase. CBM: Carbohydrate-binding module. GH: Glycosyl hydrolase. GT: Glycosyltransferase. PL: Polysaccharide lyase. When a family number is absent, the full information was missing from the Uniprot Database.

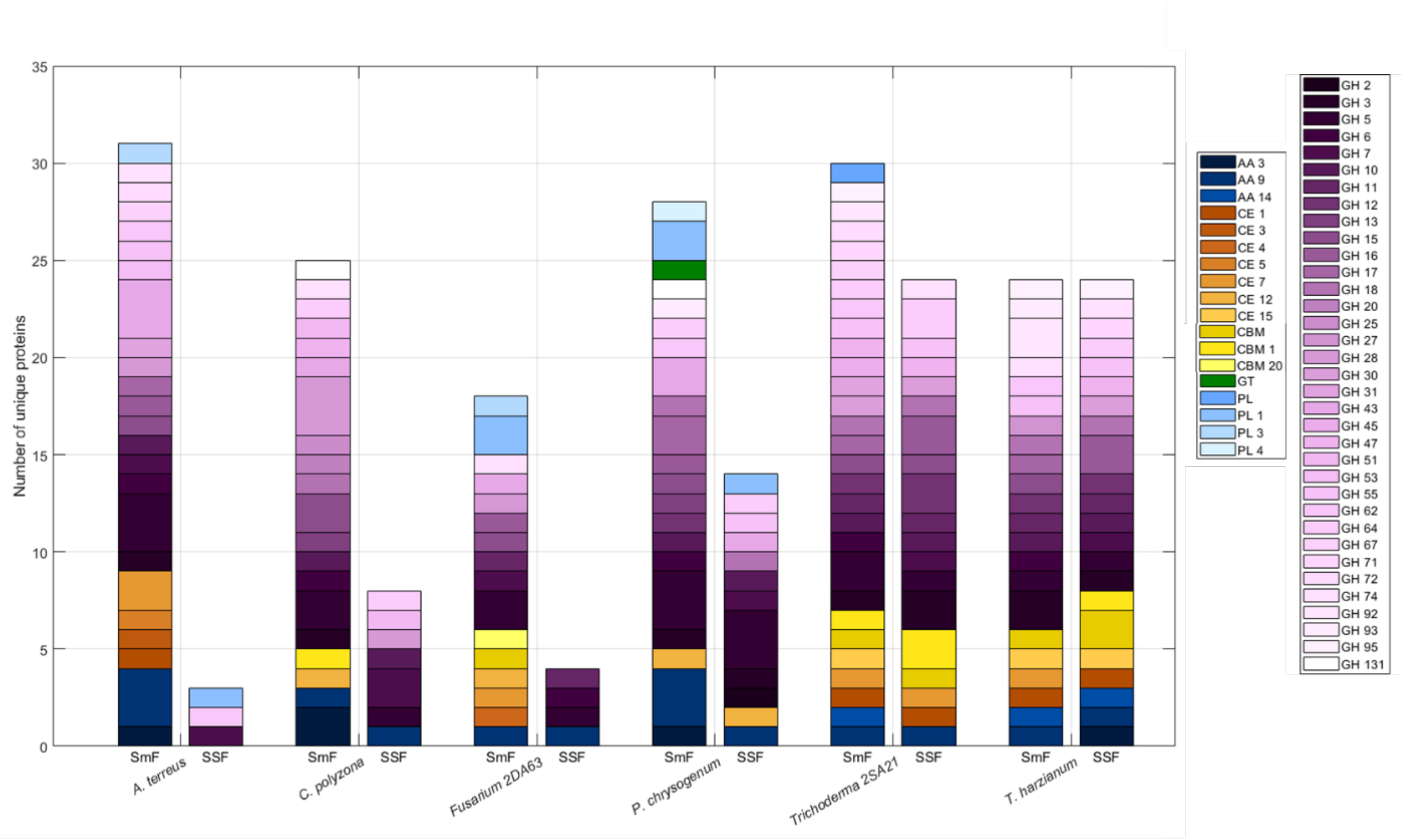


Fig. S9: Global CAZyme families distribution (number of unique proteins) in SmF and SSF with distinguished fungi. AA: Auxiliary Activities. CE: Carbohydrate Esterase. CBM: Carbohydrate-binding module. GH: Glycosyl hydrolase. GT: Glycosyltransferase. PL: Polysaccharide lyase. When a family number is absent, the full information was missing from the Uniprot Database.
