## Supplementary Data 3 - Codes for "Comparative Secretome Analysis and Enzyme Cocktail Optimization of Six Fungal Species Under Solid-State and Submerged Fermentation for Lignocellulosic Saccharification of Flax Shives": Analyse_DOE_Exp_OptiFungi.m

```
% analyse_lattice_bayes_multicritere.m
% -------------------------------------------------------------------------
% Bayesian optimization for fungal mixture design
% Objective: Maximize glucose production and glucose purity
% Outputs: Recommended strain composition and predicted saccharification profile
% -------------------------------------------------------------------------
clearvars -except X Y; close all; clc;

%% === 1) Data Loading and Validation ===
if ~exist('X','var') || ~exist('Y','var')
    error('Load matrices X (6 columns: strains) and Y (6 columns: sugars) first.');
end

assert(size(X,2) == 6, 'X must have 6 columns (Strain1..Strain6).');
assert(size(Y,2) == 6, 'Y must have 6 columns (Cellobiose, Glucose, Xylose, Galactose, Mannose).');

% Strain names (modify according to your strains)
strainNames = {'AT', 'CP', 'T21', 'PC', 'F63', 'TH'};

% Sugar names
sugarNames = {'Cellobiose', 'Glucose', 'Xylose', 'Galactose', 'Mannose'};

% Normalize mixture design (sum = 1)
rowSum = sum(X, 2);
X = X ./ rowSum;

%% === 2) Response Variables Calculation ===
% Individual sugars
cellobiose = Y(:,1);
glucose    = Y(:,2);
xylose     = Y(:,3);
galactose  = Y(:,4);

mannose    = Y(:,6);

% Derived metrics
totalSugars = sum(Y, 2);
monosaccharides = glucose + xylose + galactose + mannose;
glucosePurity = glucose ./ totalSugars;
glucosePurityMono = glucose ./ monosaccharides; % Purity among monosaccharides only

%% === 3) GPR Model Training for All Sugars ===
fprintf('Training GPR models...\n');

kernelFcn = 'ardsquaredexponential';

gpr_cellobiose = fitrgp(X, cellobiose, 'KernelFunction', kernelFcn, 'Standardize', true);
gpr_glucose    = fitrgp(X, glucose,    'KernelFunction', kernelFcn, 'Standardize', true);
gpr_xylose     = fitrgp(X, xylose,     'KernelFunction', kernelFcn, 'Standardize', true);
gpr_galactose  = fitrgp(X, galactose,  'KernelFunction', kernelFcn, 'Standardize', true);

gpr_mannose    = fitrgp(X, mannose,    'KernelFunction', kernelFcn, 'Standardize', true);
gpr_purity     = fitrgp(X, glucosePurity, 'KernelFunction', kernelFcn, 'Standardize', true);

fprintf('GPR models trained successfully.\n\n');

%% === 4) Multi-Objective Optimization Weights ===
w_glucose = 0.5;   % Weight for glucose yield (adjust as needed)
w_purity  = 0.5;   % Weight for glucose purity (adjust as needed)
xi = 0.01;         % Exploration parameter for Expected Improvement

fprintf('Optimization weights: Glucose yield = %.1f, Purity = %.1f\n\n', w_glucose, w_purity);

%% === 5) Acquisition Function Definition ===
EI_fun = @(x5) negEI_multi_for_fmincon(x5, gpr_glucose, gpr_purity, xi, w_glucose, w_purity);

%% === 6) Optimization Constraints and Options ===
lb = zeros(5,1);        % Lower bounds
ub = ones(5,1);         % Upper bounds
nonlcon = @(x) deal(sum(x) - 1, []);  % Inequality constraint: sum(x) <= 1

opts = optimoptions('fmincon', ...
    'Display', 'none', ...
    'Algorithm', 'sqp', ...
    'MaxFunctionEvaluations', 10000, ...
    'OptimalityTolerance', 1e-8);

%% === 7) Multi-Start Optimization ===
fprintf('Running multi-start optimization...\n');

Xinit = X(:, 1:5);
nStarts = min(50, size(Xinit,1) + 20);
rng(42);  % Reproducibility

% Generate starting points
starts = [Xinit; rand(nStarts - size(Xinit,1), 5)];
starts = starts(1:nStarts, :);

bestEI = -Inf;
bestX  = [];

for i = 1:size(starts, 1)
    x0 = starts(i,:)';
    
    % Ensure feasibility
    if sum(x0) > 1
        x0 = x0 / sum(x0) * 0.95;
    end
    
    try
        [xopt, fval] = fmincon(@(x) EI_fun(x'), x0, [], [], [], [], lb, ub, nonlcon, opts);
    catch
        continue
    end
    
    EIval = -fval;
    if EIval > bestEI
        bestEI = EIval;
        bestX = xopt;
    end
end

if isempty(bestX)
    error('Optimization failed: no feasible point found.');
end

% Complete the mixture (6th component)
x6 = max(0, 1 - sum(bestX));  % Ensure non-negative
x_recommended = [bestX' x6];

% Normalize to ensure sum = 1
x_recommended = x_recommended / sum(x_recommended);

fprintf('Optimization completed.\n\n');

%% === 8) Predictions for All Sugars ===
[pred_cellobiose, std_cellobiose] = predict(gpr_cellobiose, x_recommended);
[pred_glucose, std_glucose]       = predict(gpr_glucose, x_recommended);
[pred_xylose, std_xylose]         = predict(gpr_xylose, x_recommended);
[pred_galactose, std_galactose]   = predict(gpr_galactose, x_recommended);

[pred_mannose, std_mannose]       = predict(gpr_mannose, x_recommended);
[pred_purity, std_purity]         = predict(gpr_purity, x_recommended);

% Ensure non-negative predictions
pred_cellobiose = max(0, pred_cellobiose);
pred_glucose    = max(0, pred_glucose);
pred_xylose     = max(0, pred_xylose);
pred_galactose  = max(0, pred_galactose);

pred_mannose    = max(0, pred_mannose);

% Calculate totals
pred_total = pred_cellobiose + pred_glucose + pred_xylose + pred_galactose + pred_mannose;
pred_monosaccharides = pred_glucose + pred_xylose + pred_galactose + pred_mannose;

% Calculate purity metrics
calc_purity_total = pred_glucose / pred_total * 100;
calc_purity_mono = pred_glucose / pred_monosaccharides * 100;

%% === 9) Display Results ===
fprintf('==========================================================\n');
fprintf('           RECOMMENDED STRAIN COMPOSITION                 \n');
fprintf('==========================================================\n');
for k = 1:6
    if x_recommended(k) > 0.001
        fprintf('  %-8s : %6.2f %%\n', strainNames{k}, x_recommended(k)*100);
    else
        fprintf('  %-8s : %6.2f %% (excluded)\n', strainNames{k}, x_recommended(k)*100);
    end
end
fprintf('----------------------------------------------------------\n');
fprintf('  Total    : %6.2f %%\n', sum(x_recommended)*100);
fprintf('==========================================================\n\n');

fprintf('==========================================================\n');
fprintf('         PREDICTED SACCHARIFICATION PROFILE               \n');
fprintf('==========================================================\n');
fprintf('  Sugar         | Conc. (mM)  |  Std Dev   |  %% Total\n');
fprintf('----------------------------------------------------------\n');
fprintf('  Cellobiose    | %8.4f    | ± %6.4f  | %6.2f %%\n', pred_cellobiose, std_cellobiose, pred_cellobiose/pred_total*100);
fprintf('  Glucose       | %8.4f    | ± %6.4f  | %6.2f %%\n', pred_glucose, std_glucose, pred_glucose/pred_total*100);
fprintf('  Xylose        | %8.4f    | ± %6.4f  | %6.2f %%\n', pred_xylose, std_xylose, pred_xylose/pred_total*100);
fprintf('  Galactose     | %8.4f    | ± %6.4f  | %6.2f %%\n', pred_galactose, std_galactose, pred_galactose/pred_total*100);
fprintf('  Mannose       | %8.4f    | ± %6.4f  | %6.2f %%\n', pred_mannose, std_mannose, pred_mannose/pred_total*100);
fprintf('----------------------------------------------------------\n');
fprintf('  TOTAL SUGARS  | %8.4f    |            |\n', pred_total);
fprintf('  Monosacch.    | %8.4f    |            |\n', pred_monosaccharides);
fprintf('==========================================================\n\n');

fprintf('==========================================================\n');
fprintf('              OPTIMIZATION METRICS                        \n');
fprintf('==========================================================\n');
fprintf('  Glucose purity (vs total)   : %6.2f %%\n', calc_purity_total);
fprintf('  Glucose purity (vs mono.)   : %6.2f %%\n', calc_purity_mono);
fprintf('  GPR predicted purity        : %6.4f ± %.4f\n', pred_purity, std_purity);
fprintf('  Expected Improvement (EI)   : %8.6f\n', bestEI);
fprintf('  Combined score (%.1f/%.1f)    : %6.4f\n', w_glucose, w_purity, ...
    w_glucose * pred_glucose + w_purity * pred_purity);
fprintf('==========================================================\n\n');

%% === 10) Comparison with Best Observed Data ===
fprintf('==========================================================\n');
fprintf('           COMPARISON WITH EXPERIMENTAL DATA              \n');
fprintf('==========================================================\n');

% Best glucose yield
[bestGlu, idxGlu] = max(glucose);
fprintf('\nBest observed GLUCOSE yield (Mix %d):\n', idxGlu);
fprintf('  Glucose: %.4f mM | Purity: %.2f %%\n', bestGlu, glucosePurity(idxGlu)*100);
fprintf('  Composition: ');
for k = 1:6
    if X(idxGlu, k) > 0.001
        fprintf('%s=%.2f%% ', strainNames{k}, X(idxGlu,k)*100);
    end
end
fprintf('\n');

% Best purity
[bestPur, idxPur] = max(glucosePurity);
fprintf('\nBest observed PURITY (Mix %d):\n', idxPur);
fprintf('  Glucose: %.4f mM | Purity: %.2f %%\n', glucose(idxPur), bestPur*100);
fprintf('  Composition: ');
for k = 1:6
    if X(idxPur, k) > 0.001
        fprintf('%s=%.2f%% ', strainNames{k}, X(idxPur,k)*100);
    end
end
fprintf('\n');

% Best total sugars
[bestTot, idxTot] = max(totalSugars);
fprintf('\nBest observed TOTAL SUGARS (Mix %d):\n', idxTot);
fprintf('  Total: %.4f mM | Glucose: %.4f mM | Purity: %.2f %%\n', ...
    bestTot, glucose(idxTot), glucosePurity(idxTot)*100);
fprintf('  Composition: ');
for k = 1:6
    if X(idxTot, k) > 0.001
        fprintf('%s=%.2f%% ', strainNames{k}, X(idxTot,k)*100);
    end
end
fprintf('\n');
fprintf('==========================================================\n');

%% === 11) Export Results to Structure ===
results.recommended_composition = x_recommended;
results.strain_names = strainNames;
results.predicted_sugars = struct(...
    'cellobiose', pred_cellobiose, ...
    'glucose', pred_glucose, ...
    'xylose', pred_xylose, ...
    'galactose', pred_galactose, ...
    'mannose', pred_mannose, ...
    'total', pred_total);
results.predicted_std = struct(...
    'cellobiose', std_cellobiose, ...
    'glucose', std_glucose, ...
    'xylose', std_xylose, ...
    'galactose', std_galactose, ...
    'mannose', std_mannose);
results.purity = struct(...
    'vs_total', calc_purity_total, ...
    'vs_monosaccharides', calc_purity_mono);
results.optimization = struct(...
    'EI', bestEI, ...
    'weights', [w_glucose, w_purity]);

fprintf('\nResults exported to "results" structure.\n');

%% === End of Script ===
% -------------------------------------------------------------------------
```
