## Supplementary Data 3 - Codes for "Comparative Secretome Analysis and Enzyme Cocktail Optimization of Six Fungal Species Under Solid-State and Submerged Fermentation for Lignocellulosic Saccharification of Flax Shives": Biomek_prog_DOE_fungi_v2.pdf

Method
Author: Beckman Coulter
Description:

---

Start
Define the following values for this method:
  Sd_Mix = 200
  nb_Mix = 5
  HPLC = 1
  Barman2 = 0
  Barman = 1

---

Comment
Description:
Add 1 mL of the negative control to well F6 of the mixing plate.

---

Comment
Description:
Place a U-shaped tray under the filters to collect the filtrate.

---

Comment
Description:
Sterile hood activation for Barman.

---

If
If "Barman = 1":

---

Then

---

Device Action
Send the following command to "Simple1": "On".
Parameters:

---

Start heating the device.

---

---

Transfer From File
Using Pod2, execute the following transfer:
Use the file "C:\Users\Beckman Coulter\Documents\REALCAT Users\PEECFUEL\Biomek TFF BarmanFungi.csv" (which has a header row) to read in the following data:
  Source position from column 1
  Source well from column 2
  Destination position from column 3
  Destination well from column 4
  Volume from column 5
Skip zero volume transfers.
From: Reserv_Modular_40mL, Water
Use the following custom technique:
  Use the following pipetting template: S8 Pipetting
  Calibration Offset: 5.227
  Calibration Slope: 1.033
  Minimum Pipetting Height: 0.5 mm
  Prewet: False
  Aspirate Blowout: True
  Follow Liquid: True
  Height: 1 mm from the bottom
  Mix: True
  Mix Aspirate Speed: 200µL/s
  Mix Aspirate Height: 1 mm from the bottom
  Mix Dispense Speed: 200µL/s
  Mix Dispense Height: 1 mm from the bottom
  Mix Count: =nb_Mix
  Mix Volume: 200 µL
  Operation speed: 5µL/s
  Tip Touch: False
  Trailing Air Gap: True
  Override Liquid Type Settings
    Trailing Air Gap Volume: 5 µL

Aspirate Delay: 200 ms
Aspirate Speed: 100 µL/s
Blowout Volume: 25 µL
Blowout Delay: 200 ms
Prewet Overage: 0 µL
Prewet Delay: 200 ms
Dispense Delay: 200 ms
Dispense Speed: 100 µL/s
Dispense Cutoff Velocity: 112.5 µL/s
Tip Touch Height: -1.5 mm from the top
Tip Touch Angle: 0
Tip Touch Speed: 50 %
Tip Touch Delay: 500 ms

Override the technique height by moving to -4 mm from the liquid.
To: DeepWell_96_V_Square_Greiner, Water
Set the mark at the last well transferred.
Auto-select the technique.
Override the technique height by moving to 2 mm from the liquid.
Dispense up to 1 time(s) per draw.
Load Tips_1070 onto the pod.
Unload the tips when finished.
Change tips between transfers.
Split transfer volumes that exceed tip capacity, do not clean tips between partial transfers.
All probes will be used.

---

Comment
Description:
Transfer 0.7 mL into the Eppendorf tubes using the pipette.

---

Pause
Pause everything while waiting for a user response to the following prompt: " Place the racks with the Eppendorf tubes".

---

View Data Sets

---

Transfer
Using Pod2, execute the following transfer:
From: DOE_Enz, with the following pattern:

| | 1 | 2 | 3 | 4 | 5 | 6 | 7 | 8 | 9 | 10 | 11 | 12 |
|---|---|---|---|---|---|---|---|---|---|---|---|---|
| A | >< | >< | >< | >< | >< | >< | >< | >< | >< | >< | >< | >< |
| B | >< | >< | >< | >< | >< | >< | >< | >< | >< | >< | >< | >< |
| C | () | () | () | () | () | () | () | () | () | () | () | () |
| D | () | () | () | () | () | () | () | () | () | () | () | () |
| E | () | () | () | () | () | () | () | () | () | () | () | () |
| F | () | () | () | () | () | () | () | () | () | () | () | () |
| G | () | () | () | () | () | () | () | () | () | () | () | () |
| H | () | () | () | () | () | () | () | () | () | () | () | () |

, Water
Proceed left first, then top to bottom.
Start from the beginning of the selection.
Set the mark at the last well transferred.
Use the following custom technique:
Use the following pipetting template: S8 Pipetting
Calibration Offset: 5.227
Calibration Slope: 1.033
Minimum Pipetting Height: 0.5 mm
Prewet: False
Aspirate Blowout: True
Follow Liquid: True
Height: -2 mm from the liquid
Mix: True
Mix Aspirate Speed: 200µL/s
Mix Aspirate Height: -2 mm from the liquid
Mix Dispense Speed: 200µL/s
Mix Dispense Height: -2 mm from the liquid
Mix Count: =nb_Mix

Mix Volume: 500 µL
Operation speed: 5µL/s
Tip Touch: False
Trailing Air Gap: True
Override Liquid Type Settings
Trailing Air Gap Volume: 5 µL
Aspirate Delay: 200 ms
Aspirate Speed: 100 µL/s
Blowout Volume: 25 µL
Blowout Delay: 200 ms
Prewet Overage: 0 µL
Prewet Delay: 200 ms
Dispense Delay: 200 ms
Dispense Speed: 100 µL/s
Dispense Cutoff Velocity: 112.5 µL/s
Tip Touch Height: -1.5 mm from the top
Tip Touch Angle: 0
Tip Touch Speed: 50 %
Tip Touch Delay: 500 ms

To: Sacch_DOE_1, 700µL, with the following pattern:

```
   1  2  3  4  5  6
 A >< >< >< >< >< ><
 B >< >< >< >< >< ><
 C >< >< >< >< >< ><
 D >< >< >< >< >< ><
```

, Water
Proceed left first, then top to bottom.
Start from the beginning of the selection.
Do not set the mark.
Use the following custom technique:
Use the following pipetting template: S8 Pipetting
Calibration Offset: 5.227
Calibration Slope: 1.033
Minimum Pipetting Height: 0.5 mm
Prewet: False
Blowout: True
Follow Liquid: True
Height: 2 mm from the liquid
Mix: False
Mix Aspirate Speed: 50µL/s
Mix Aspirate Height: 2 mm from the bottom
Mix Dispense Speed: 50µL/s
Mix Dispense Height: -2 mm from the liquid
Mix Count: 3
Mix Volume: 200 µL
Operation speed: 5µL/s
Tip Touch: False
Override Liquid Type Settings
Trailing Air Gap Volume: 5 µL
Aspirate Delay: 200 ms
Aspirate Speed: 50 µL/s
Blowout Volume: 25 µL
Blowout Delay: 200 ms
Prewet Overage: 0 µL
Prewet Delay: 200 ms
Dispense Delay: 200 ms
Dispense Speed: 100 µL/s
Dispense Cutoff Velocity: 112.5 µL/s
Tip Touch Height: -1.5 mm from the top
Tip Touch Angle: 0
Tip Touch Speed: 50 %
Tip Touch Delay: 500 ms

Dispense up to 1 time(s) per draw.
Create 1 replicate(s) of each source well.
Load Tips_Sacchar onto the pod.
Unload the tips when finished.

Change tips between transfers.
Split transfer volumes that exceed tip capacity, do not clean tips between partial transfers.
Stop when finished with Destinations.
All probes will be used.

---

Transfer
Using Pod2, execute the following transfer:
From: DOE_Enz, with the following pattern:

```
   1  2  3  4  5  6  7  8  9  10 11 12
 A () () () () () () () () () () () ()
 B () () () () () () () () () () () ()
 C >< >< >< >< >< >< >< >< >< >< >< ><
 D >< >< >< >< >< >< >< >< >< >< >< ><
 E () () () () () () () () () () () ()
 F () () () () () () () () () () () ()
 G () () () () () () () () () () () ()
 H () () () () () () () () () () () ()
```

, Water
Proceed left first, then top to bottom.
Start from the beginning of the selection.
Set the mark at the last well transferred.
Use the following custom technique:
  Use the following pipetting template: S8 Pipetting
  Calibration Offset: 5.227
  Calibration Slope: 1.033
  Minimum Pipetting Height: 0.5 mm
  Prewet: False
  Aspirate Blowout: True
  Follow Liquid: True
  Height: -2 mm from the liquid
  Mix: True
  Mix Aspirate Speed: 200µL/s
  Mix Aspirate Height: -2 mm from the liquid
  Mix Dispense Speed: 200µL/s
  Mix Dispense Height: -2 mm from the liquid
  Mix Count: =nb_Mix
  Mix Volume: 500 µL
  Operation speed: 5µL/s
  Tip Touch: False
  Trailing Air Gap: True
  Override Liquid Type Settings
    Trailing Air Gap Volume: 5 µL
    Aspirate Delay: 200 ms
    Aspirate Speed: 100 µL/s
    Blowout Volume: 25 µL
    Blowout Delay: 200 ms
    Prewet Overage: 0 µL
    Prewet Delay: 200 ms
    Dispense Delay: 200 ms
    Dispense Speed: 100 µL/s
    Dispense Cutoff Velocity: 112.5 µL/s
    Tip Touch Height: -1.5 mm from the top
    Tip Touch Angle: 0
    Tip Touch Speed: 50 %
    Tip Touch Delay: 500 ms

To: Sacch_DOE_2, 700µL, with the following pattern:

```
   1  2  3  4  5  6
 A >< >< >< >< >< ><
 B >< >< >< >< >< ><
 C >< >< >< >< >< ><
 D >< >< >< >< >< ><
```

, Water
Proceed left first, then top to bottom.
Start from the beginning of the selection.
Do not set the mark.
Use the following custom technique:

Use the following pipetting template: S8 Pipetting
Calibration Offset: 5.227
Calibration Slope: 1.033
Minimum Pipetting Height: 0.5 mm
Prewet: False
Blowout: True
Follow Liquid: True
Height: 2 mm from the liquid
Mix: False
Mix Aspirate Speed: 50µL/s
Mix Aspirate Height: 2 mm from the bottom
Mix Dispense Speed: 50µL/s
Mix Dispense Height: -2 mm from the liquid
Mix Count: 3
Mix Volume: 200 µL
Operation speed: 5µL/s
Tip Touch: False
Override Liquid Type Settings
Trailing Air Gap Volume: 5 µL
Aspirate Delay: 200 ms
Aspirate Speed: 50 µL/s
Blowout Volume: 25 µL
Blowout Delay: 200 ms
Prewet Overage: 0 µL
Prewet Delay: 200 ms
Dispense Delay: 200 ms
Dispense Speed: 100 µL/s
Dispense Cutoff Velocity: 112.5 µL/s
Tip Touch Height: -1.5 mm from the top
Tip Touch Angle: 0
Tip Touch Speed: 50 %
Tip Touch Delay: 500 ms

Dispense up to 1 time(s) per draw.
Create 1 replicate(s) of each source well.
Load Tips_Sacchar onto the pod.
Unload the tips when finished.
Change tips between transfers.
Split transfer volumes that exceed tip capacity, do not clean tips between partial transfers.
Stop when finished with Destinations.
All probes will be used.

---

Transfer
Using Pod2, execute the following transfer:
From: DOE_Enz, with the following pattern:

| | 1 | 2 | 3 | 4 | 5 | 6 | 7 | 8 | 9 | 10 | 11 | 12 |
|---|---|---|---|---|---|---|---|---|---|---|---|---|
| A | () | () | () | () | () | () | () | () | () | () | () | () |
| B | () | () | () | () | () | () | () | () | () | () | () | () |
| C | () | () | () | () | () | () | () | () | () | () | () | () |
| D | () | () | () | () | () | () | () | () | () | () | () | () |
| E | >< | >< | >< | >< | >< | >< | >< | >< | >< | >< | >< | >< |
| F | >< | >< | >< | >< | >< | >< | () | () | () | () | () | () |
| G | () | () | () | () | () | () | () | () | () | () | () | () |
| H | () | () | () | () | () | () | () | () | () | () | () | () |

, Water
Proceed left first, then top to bottom.
Start from the beginning of the selection.
Set the mark at the last well transferred.
Use the following custom technique:
Use the following pipetting template: S8 Pipetting
Calibration Offset: 5.227
Calibration Slope: 1.033
Minimum Pipetting Height: 0.5 mm
Prewet: False
Aspirate Blowout: True
Follow Liquid: True
Height: -2 mm from the liquid

Mix: True
Mix Aspirate Speed: 200µL/s
Mix Aspirate Height: -2 mm from the liquid
Mix Dispense Speed: 200µL/s
Mix Dispense Height: -2 mm from the liquid
Mix Count: =nb_Mix
Mix Volume: 500 µL
Operation speed: 5µL/s
Tip Touch: False
Trailing Air Gap: True
Override Liquid Type Settings
Trailing Air Gap Volume: 5 µL
Aspirate Delay: 200 ms
Aspirate Speed: 100 µL/s
Blowout Volume: 25 µL
Blowout Delay: 200 ms
Prewet Overage: 0 µL
Prewet Delay: 200 ms
Dispense Delay: 200 ms
Dispense Speed: 100 µL/s
Dispense Cutoff Velocity: 112.5 µL/s
Tip Touch Height: -1.5 mm from the top
Tip Touch Angle: 0
Tip Touch Speed: 50 %
Tip Touch Delay: 500 ms

To: Sacch_DOE_3, 700µL, with the following pattern:

```
   1  2  3  4  5  6
 A >< >< >< >< >< ><
 B >< >< >< >< >< ><
 C >< >< >< >< >< ><
 D () () () () () ()
```

, Water
Proceed left first, then top to bottom.
Start from the beginning of the selection.
Do not set the mark.
Use the following custom technique:
Use the following pipetting template: S8 Pipetting
Calibration Offset: 5.227
Calibration Slope: 1.033
Minimum Pipetting Height: 0.5 mm
Prewet: False
Blowout: True
Follow Liquid: True
Height: 2 mm from the liquid
Mix: False
Mix Aspirate Speed: 50µL/s
Mix Aspirate Height: 2 mm from the bottom
Mix Dispense Speed: 50µL/s
Mix Dispense Height: -2 mm from the liquid
Mix Count: 3
Mix Volume: 200 µL
Operation speed: 5µL/s
Tip Touch: False
Override Liquid Type Settings
Trailing Air Gap Volume: 5 µL
Aspirate Delay: 200 ms
Aspirate Speed: 50 µL/s
Blowout Volume: 25 µL
Blowout Delay: 200 ms
Prewet Overage: 0 µL
Prewet Delay: 200 ms
Dispense Delay: 200 ms
Dispense Speed: 100 µL/s
Dispense Cutoff Velocity: 112.5 µL/s
Tip Touch Height: -1.5 mm from the top
Tip Touch Angle: 0
Tip Touch Speed: 50 %

Tip Touch Delay: 500 ms

Dispense up to 1 time(s) per draw.
Create 1 replicate(s) of each source well.
Load Tips_Sacchar onto the pod.
Unload the tips when finished.
Change tips between transfers.
Split transfer volumes that exceed tip capacity, do not clean tips between partial transfers.
Stop when finished with Destinations.
All probes will be used.

---

Comment
Description:
Saccharification 48h
Comment:
Seal the tubes and incubate at 30°C, 200 rpm for 48 hours

---

Device Action
Send the following command to "Simple1": "Off".
Parameters:

---

Stop heating the device.

---

End

---

Else

---

End

---

If
If "HPLC = 1":

---

Then

---

Comment
Description:
HPLC saccharification analysis.

---

Comment:
Stop saccharification 5 min 100 degC
10 minutes 11000xg
Place the Eppendorf tubes on the racks

---

Comment
Description:
Saccharifications transfert done.

---

Transfer
Using Pod2, execute the following transfer:
From: Sacch_DOE_1, with the following pattern:

```
   1  2  3  4  5  6
 A >< >< >< >< >< ><
 B >< >< >< >< >< ><
 C >< >< >< >< >< ><
 D >< >< >< >< >< ><
```

, Water
Proceed left first, then top to bottom.
Start from the beginning of the selection.
Set the mark at the last well transferred.
Use the following custom technique:
  Use the following pipetting template: S8 Pipetting
  Calibration Offset: 5.227
  Calibration Slope: 1.033
  Minimum Pipetting Height: 0.5 mm
  Prewet: False
  Aspirate Blowout: True
  Follow Liquid: True

Height: -1 mm from the liquid
Mix: False
Mix Aspirate Speed: =Sd_MixµL/s
Mix Aspirate Height: 2 mm from the liquid
Mix Dispense Speed: =Sd_MixµL/s
Mix Dispense Height: -2 mm from the liquid
Mix Count: 3
Mix Volume: 200 µL
Operation speed: 5µL/s
Tip Touch: False
Trailing Air Gap: True
Override Liquid Type Settings
Trailing Air Gap Volume: 5 µL
Aspirate Delay: 200 ms
Aspirate Speed: 20 µL/s
Blowout Volume: 25 µL
Blowout Delay: 200 ms
Prewet Overage: 0 µL
Prewet Delay: 200 ms
Dispense Delay: 200 ms
Dispense Speed: 100 µL/s
Dispense Cutoff Velocity: 112.5 µL/s
Tip Touch Height: -1.5 mm from the top
Tip Touch Angle: 0
Tip Touch Speed: 50 %
Tip Touch Delay: 500 ms

Override the technique height by moving to -2 mm from the liquid.
To: Sacch_Transfert, 650µL, with the following pattern:

| | 1 | 2 | 3 | 4 | 5 | 6 | 7 | 8 | 9 | 10 | 11 | 12 |
|---|---|---|---|---|---|---|---|---|---|---|---|---|
| A | >< | >< | >< | >< | >< | >< | >< | >< | >< | >< | >< | >< |
| B | >< | >< | >< | >< | >< | >< | >< | >< | >< | >< | >< | >< |
| C | () | () | () | () | () | () | () | () | () | () | () | () |
| D | () | () | () | () | () | () | () | () | () | () | () | () |
| E | () | () | () | () | () | () | () | () | () | () | () | () |
| F | () | () | () | () | () | () | () | () | () | () | () | () |
| G | () | () | () | () | () | () | () | () | () | () | () | () |
| H | () | () | () | () | () | () | () | () | () | () | () | () |

, Water
Proceed left first, then top to bottom.
Start from the beginning of the selection.
Set the mark at the last well transferred.
Use the following custom technique:
Use the following pipetting template: S8 Pipetting
Calibration Offset: 5.227
Calibration Slope: 1.033
Minimum Pipetting Height: 0.5 mm
Prewet: False
Blowout: True
Follow Liquid: True
Height: 2 mm from the liquid
Mix: False
Mix Aspirate Speed: 50µL/s
Mix Aspirate Height: 2 mm from the bottom
Mix Dispense Speed: 50µL/s
Mix Dispense Height: -2 mm from the liquid
Mix Count: 3
Mix Volume: 200 µL
Operation speed: 5µL/s
Tip Touch: True
Override Liquid Type Settings
Trailing Air Gap Volume: 5 µL
Aspirate Delay: 200 ms
Aspirate Speed: 50 µL/s
Blowout Volume: 25 µL
Blowout Delay: 200 ms
Prewet Overage: 0 µL
Prewet Delay: 200 ms

```
    Dispense Delay: 200 ms
    Dispense Speed: 100 µL/s
    Dispense Cutoff Velocity: 112.5 µL/s
    Tip Touch Height: -1.5 mm from the top
    Tip Touch Angle: 0
    Tip Touch Speed: 50 %
    Tip Touch Delay: 200 ms

Dispense up to 1 time(s) per draw.
Create 1 replicate(s) of each source well.
Load Tips_Sacchar onto the pod.
Unload the tips when finished.
Change tips between transfers.
Do not split transfer volumes that exceed tip capacity.
Stop when finished with Destinations.
All probes will be used.
```

---

```
Pause
Pause everything while waiting for a user response to the following prompt: " Is the
Eppendorf rack 2 ready?".
```

---

```
Transfer
Using Pod2, execute the following transfer:
From: Sacch_DOE_2, with the following pattern:
   1  2  3  4  5  6
 A >< >< >< >< >< ><
 B >< >< >< >< >< ><
 C >< >< >< >< >< ><
 D >< >< >< >< >< ><
, Water
Proceed left first, then top to bottom.
Start from the beginning of the selection.
Set the mark at the last well transferred.
Use the following custom technique:
  Use the following pipetting template: S8 Pipetting
  Calibration Offset: 5.227
  Calibration Slope: 1.033
  Minimum Pipetting Height: 0.5 mm
  Prewet: False
  Aspirate Blowout: True
  Follow Liquid: True
  Height: -1 mm from the liquid
  Mix: False
  Mix Aspirate Speed: =Sd_MixµL/s
  Mix Aspirate Height: 2 mm from the liquid
  Mix Dispense Speed: =Sd_MixµL/s
  Mix Dispense Height: -2 mm from the liquid
  Mix Count: 3
  Mix Volume: 200 µL
  Operation speed: 5µL/s
  Tip Touch: False
  Trailing Air Gap: True
  Override Liquid Type Settings
    Trailing Air Gap Volume: 5 µL
    Aspirate Delay: 200 ms
    Aspirate Speed: 20 µL/s
    Blowout Volume: 25 µL
    Blowout Delay: 200 ms
    Prewet Overage: 0 µL
    Prewet Delay: 200 ms
    Dispense Delay: 200 ms
    Dispense Speed: 100 µL/s
    Dispense Cutoff Velocity: 112.5 µL/s
    Tip Touch Height: -1.5 mm from the top
    Tip Touch Angle: 0
    Tip Touch Speed: 50 %
    Tip Touch Delay: 500 ms
```

Override the technique height by moving to -2 mm from the liquid.
To: Sacch_Transfert, 650µL, with the following pattern:

```
   1  2  3  4  5  6  7  8  9  10 11 12
 A () () () () () () () () () () () ()
 B () () () () () () () () () () () ()
 C >< >< >< >< >< >< >< >< >< >< >< ><
 D >< >< >< >< >< >< >< >< >< >< >< ><
 E () () () () () () () () () () () ()
 F () () () () () () () () () () () ()
 G () () () () () () () () () () () ()
 H () () () () () () () () () () () ()
```

, Water
Proceed left first, then top to bottom.
Start from the beginning of the selection.
Set the mark at the last well transferred.
Use the following custom technique:
  Use the following pipetting template: S8 Pipetting
  Calibration Offset: 5.227
  Calibration Slope: 1.033
  Minimum Pipetting Height: 0.5 mm
  Prewet: False
  Blowout: True
  Follow Liquid: True
  Height: 2 mm from the liquid
  Mix: False
  Mix Aspirate Speed: 50µL/s
  Mix Aspirate Height: 2 mm from the bottom
  Mix Dispense Speed: 50µL/s
  Mix Dispense Height: -2 mm from the liquid
  Mix Count: 3
  Mix Volume: 200 µL
  Operation speed: 5µL/s
  Tip Touch: False
  Override Liquid Type Settings
    Trailing Air Gap Volume: 5 µL
    Aspirate Delay: 200 ms
    Aspirate Speed: 50 µL/s
    Blowout Volume: 25 µL
    Blowout Delay: 200 ms
    Prewet Overage: 0 µL
    Prewet Delay: 200 ms
    Dispense Delay: 200 ms
    Dispense Speed: 100 µL/s
    Dispense Cutoff Velocity: 112.5 µL/s
    Tip Touch Height: -1.5 mm from the top
    Tip Touch Angle: 0
    Tip Touch Speed: 50 %
    Tip Touch Delay: 500 ms

Dispense up to 1 time(s) per draw.
Create 1 replicate(s) of each source well.
Load Tips_Sacchar onto the pod.
Unload the tips when finished.
Change tips between transfers.
Do not split transfer volumes that exceed tip capacity.
Stop when finished with Destinations.
All probes will be used.

---

Pause
Pause everything while waiting for a user response to the following prompt: " Is the Eppendorf rack 3 ready?".

---

Transfer
Using Pod2, execute the following transfer:
From: Sacch_DOE_3, with the following pattern:

```
   1  2  3  4  5  6
 A >< >< >< >< >< ><
 B >< >< >< >< >< ><
```

| | | | | | | |
|---|---|---|---|---|---|---|
| C | >< | >< | >< | >< | >< | >< |
| D | () | () | () | () | () | () |

, Water
Proceed left first, then top to bottom.
Start from the beginning of the selection.
Set the mark at the last well transferred.
Use the following custom technique:
  Use the following pipetting template: S8 Pipetting
  Calibration Offset: 5.227
  Calibration Slope: 1.033
  Minimum Pipetting Height: 0.5 mm
  Prewet: False
  Aspirate Blowout: True
  Follow Liquid: True
  Height: -1 mm from the liquid
  Mix: False
  Mix Aspirate Speed: =Sd_MixµL/s
  Mix Aspirate Height: 2 mm from the liquid
  Mix Dispense Speed: =Sd_MixµL/s
  Mix Dispense Height: -2 mm from the liquid
  Mix Count: 3
  Mix Volume: 200 µL
  Operation speed: 5µL/s
  Tip Touch: False
  Trailing Air Gap: True
  Override Liquid Type Settings
    Trailing Air Gap Volume: 5 µL
    Aspirate Delay: 200 ms
    Aspirate Speed: 20 µL/s
    Blowout Volume: 25 µL
    Blowout Delay: 200 ms
    Prewet Overage: 0 µL
    Prewet Delay: 200 ms
    Dispense Delay: 200 ms
    Dispense Speed: 100 µL/s
    Dispense Cutoff Velocity: 112.5 µL/s
    Tip Touch Height: -1.5 mm from the top
    Tip Touch Angle: 0
    Tip Touch Speed: 50 %
    Tip Touch Delay: 500 ms

Override the technique height by moving to -2 mm from the liquid.
To: Sacch_Transfert, 650µL, with the following pattern:

| | 1 | 2 | 3 | 4 | 5 | 6 | 7 | 8 | 9 | 10 | 11 | 12 |
|---|---|---|---|---|---|---|---|---|---|---|---|---|
| A | () | () | () | () | () | () | () | () | () | () | () | () |
| B | () | () | () | () | () | () | () | () | () | () | () | () |
| C | () | () | () | () | () | () | () | () | () | () | () | () |
| D | () | () | () | () | () | () | () | () | () | () | () | () |
| E | >< | >< | >< | >< | >< | >< | >< | >< | >< | >< | >< | >< |
| F | >< | >< | >< | >< | >< | >< | () | () | () | () | () | () |
| G | () | () | () | () | () | () | () | () | () | () | () | () |
| H | () | () | () | () | () | () | () | () | () | () | () | () |

, Water
Proceed left first, then top to bottom.
Start from the beginning of the selection.
Set the mark at the last well transferred.
Use the following custom technique:
  Use the following pipetting template: S8 Pipetting
  Calibration Offset: 5.227
  Calibration Slope: 1.033
  Minimum Pipetting Height: 0.5 mm
  Prewet: False
  Blowout: True
  Follow Liquid: True
  Height: 2 mm from the liquid
  Mix: False
  Mix Aspirate Speed: 50µL/s
  Mix Aspirate Height: 2 mm from the bottom

```
  Mix Dispense Speed: 50µL/s
  Mix Dispense Height: -2 mm from the liquid
  Mix Count: 3
  Mix Volume: 200 µL
  Operation speed: 5µL/s
  Tip Touch: False
  Override Liquid Type Settings
    Trailing Air Gap Volume: 5 µL
    Aspirate Delay: 200 ms
    Aspirate Speed: 50 µL/s
    Blowout Volume: 25 µL
    Blowout Delay: 200 ms
    Prewet Overage: 0 µL
    Prewet Delay: 200 ms
    Dispense Delay: 200 ms
    Dispense Speed: 100 µL/s
    Dispense Cutoff Velocity: 112.5 µL/s
    Tip Touch Height: -1.5 mm from the top
    Tip Touch Angle: 0
    Tip Touch Speed: 50 %
    Tip Touch Delay: 500 ms

Dispense up to 1 time(s) per draw.
Create 1 replicate(s) of each source well.
Load Tips_Sacchar onto the pod.
Unload the tips when finished.
Change tips between transfers.
Do not split transfer volumes that exceed tip capacity.
Stop when finished with Destinations.
All probes will be used.
```

---

Comment
Description:
Saccharifications filtration.

---

Loop
Loop from "n" = "1" to "4", incrementing by "1".

---

```
Transfer
Using Pod1, execute the following transfer:
From: Sacch_Transfert, section 1, Water
Use the following custom technique:
  Use the following pipetting template: MC Pipetting
  Calibration Offset: 0
  Calibration Slope: 1
  Minimum Pipetting Height: 0.5 mm
  Prewet: False
  Aspirate Blowout: True
  Follow Liquid: True
  Height: -2 mm from the liquid
  Mix: True
  Mix Aspirate Speed: =Sd_MixµL/s
  Mix Aspirate Height: -2 mm from the liquid
  Mix Dispense Speed: =Sd_MixµL/s
  Mix Dispense Height: -2 mm from the liquid
  Mix Count: 1
  Mix Volume: 200 µL
  Operation speed: 5µL/s
  Tip Touch: False
  Trailing Air Gap: True
  Override Liquid Type Settings
    Trailing Air Gap Volume: 5 µL
    Aspirate Delay: 200 ms
    Aspirate Speed: 100 µL/s
    Blowout Volume: 25 µL
    Blowout Delay: 200 ms
    Prewet Overage: 0 µL
    Prewet Delay: 200 ms
```

Dispense Delay: 200 ms
Dispense Speed: 100 µL/s
Dispense Cutoff Velocity: 112.5 µL/s
Tip Touch Height: -1.5 mm from the top
Tip Touch Angle: 0
Tip Touch Speed: 50 %
Tip Touch Delay: 500 ms

To: SPEBase, 200µL, section 1, Water
Auto-select the technique.
Dispense up to 1 time(s) per draw.
Load Tips_Filtre onto the pod.
Unload the tips when finished.
Do not split transfer volumes that exceed tip capacity.
Stop when finished with Destinations.

---

Group:
SPE Pump 30s

---

Device Action
Send the following command to "Simple2" (SPE Valve):
"On". Parameters:

---

Device Action
Send the following command to "Simple3" (SPE Pump):
"On". Parameters:

---

Pause
Pause "the whole system" for "5" seconds.

---

BeckmanPump: Timed Vacuum

---

Pause
Pause "the whole system" for "1" seconds.

---

Device Action
Send the following command to "Simple2": "Off".
Parameters:

---

Device Action
Send the following command to "Simple3": "Off".
Parameters:

---

End Group

---

---

End Loop

---

View Data Sets

---

End

---

Else

---

End

---

Finish
Clear current instrument setup of all labware.
Clear current device setup of all labware.
Unload disposable tips from all pods.
Move all pods and grippers to their park locations.
Clear all global variables.

---
