## Supplementary Data 3 - Codes for "Comparative Secretome Analysis and Enzyme Cocktail Optimization of Six Fungal Species Under Solid-State and Submerged Fermentation for Lignocellulosic Saccharification of Flax Shives": mixture_model_selection_and_ternary_plots.m

```
%% mixture_model_selection_and_ternary_plots.m
% Full pipeline for Scheffé mixture models
%
% Features:
% - Fit Linear, Quadratic, and Special Cubic models
% - Backward elimination with p>0.05 removal
% - One Excel file per model type (with all sugars as sheets)
% - One combined figure (4x5 grid) per sugar per model
% - Strain names directly in coefficient names (AT, CP, AT_CP, etc.)
%
% Inputs required in workspace:
% - X : n x 6 mixture proportions (rows sum to 1)
% - Y : n x 6 responses (sugars)
%
% Compatible with MATLAB 2022b

%% ================================================================
clearvars ; close all;
load('XetY.mat')
Y = Y(:,[1:4 6])

%% ---------------------------------------------------------------
% USER INPUT
%% ---------------------------------------------------------------
strain_names = {'AT', 'CP', 'T21', 'PC', 'F63', 'TH'};
sugar_names = {'Cellobiose', 'Glucose', 'Xylose', 'Galactose', 'Mannose'};
model_types = {'Linear', 'Quadratic', 'SpecialCubic', 'FullCubic'};
model_orders = [1, 2, 3, 4];  % Corresponding Scheffé orders

outFolder = 'Mixture_Results_Sheffe';
if ~exist(outFolder, 'dir')
    mkdir(outFolder);
end

[n, q] = size(X);  % q = 6 strains

%% ================================================================
% MAIN LOOP: FOR EACH MODEL TYPE
%% ================================================================

for m_type = 1:length(model_types)
    
    model_name = model_types{m_type};
    model_order = model_orders(m_type);
    
    fprintf('\n\n########################################################\n');
    fprintf('   MODEL TYPE: %s (order %d)\n', upper(model_name), model_order);
    fprintf('########################################################\n');
    
    % Create Excel file for this model type
    excel_file = sprintf('Mixture_Results_%s.xlsx', model_name);
    
    %% ============================================================
    % LOOP OVER SUGARS
    %% ============================================================
    
    for s = 1:5
        fprintf('\n--- %s - %s ---\n', model_name, sugar_names{s});
        
        y = Y(:, s);
        
        % Build design matrix with strain names
        [M, term_names] = build_scheffe_named(X, model_order, strain_names);
        
        % Display term count (first sugar only)
        if s == 1
            fprintf('Model %s: %d terms\n', model_name, length(term_names));
        end
        
        % Create table with named variables
        T = array2table(M, 'VariableNames', term_names);
        T.Response = y;
        
        % Fit model (no intercept for mixture models) - NO BACKWARD ELIMINATION
        formula = sprintf('Response ~ %s - 1', strjoin(term_names, ' + '));
        mdl_final = fitlm(T, formula);
        
        fprintf('Model fitted with %d terms\n', mdl_final.NumCoefficients);
        
        %% -------------------------------------------------------
        % EXPORT TO EXCEL
        %% -------------------------------------------------------
        
        sheet = sugar_names{s};
        
        % Coefficients table (already has strain names)
        coef = mdl_final.Coefficients;
        writetable(coef, fullfile(outFolder, excel_file), ...
            'Sheet', sheet, 'WriteRowNames', true, 'Range', 'A1');
        
        % ANOVA Summary
        anova_table = compute_anova_summary(X, y, mdl_final, strain_names);
        writetable(anova_table, fullfile(outFolder, excel_file), ...
            'Sheet', sheet, 'WriteRowNames', true, 'Range', 'H1');
        
        % Model info
        model_info = table(...
            mdl_final.Rsquared.Ordinary, ...
            mdl_final.Rsquared.Adjusted, ...
            mdl_final.RMSE, ...
            mdl_final.ModelCriterion.AIC, ...
            mdl_final.ModelCriterion.BIC, ...
            'VariableNames', {'R2', 'R2_adj', 'RMSE', 'AIC', 'BIC'});
        writetable(model_info, fullfile(outFolder, excel_file), ...
            'Sheet', sheet, 'Range', 'H12');
        
        fprintf('Excel exported: %s (sheet: %s)\n', excel_file, sheet);
        
        %% -------------------------------------------------------
        % TERNARY PLOTS - Combined figure (4x5 = 20 triplets)
        %% -------------------------------------------------------
        
        triplets = nchoosek(1:6, 3);
        
        % --- First pass: compute global min/max for colormap normalization ---
        fprintf('Computing colormap range for %s...\n', sugar_names{s});
        all_predictions = [];
        
        for t_idx = 1:size(triplets, 1)
            preds = compute_ternary_predictions(mdl_final, X, triplets(t_idx,:), strain_names);
            all_predictions = [all_predictions; preds]; %#ok<AGROW>
        end
        
        cmin = 0;  % Force minimum to 0
        cmax = max(all_predictions);
        fprintf('Colormap range: [0, %.4f]\n', cmax);
        
        % --- Second pass: create plots with normalized colormap ---
        fig = figure('Name', sprintf('%s - %s', model_name, sugar_names{s}), ...
            'NumberTitle', 'off', 'Units', 'normalized', ...
            'Position', [0.02, 0.02, 0.96, 0.92], 'Visible', 'off');
        
        tl = tiledlayout(4, 5, 'TileSpacing', 'compact', 'Padding', 'compact');
        title(tl, sprintf('Modele %s : %s (echelle: 0 - %.3f)', model_name, sugar_names{s}, cmax), ...
            'FontSize', 14, 'FontWeight', 'bold');
        
        for t_idx = 1:size(triplets, 1)
            nexttile;
            plot_ternary_surface(mdl_final, X, triplets(t_idx,:), strain_names, [cmin, cmax]);
            title(sprintf('%s-%s-%s', ...
                strain_names{triplets(t_idx,1)}, ...
                strain_names{triplets(t_idx,2)}, ...
                strain_names{triplets(t_idx,3)}), 'FontSize', 9);
        end
        
        % Save combined figure
        fig_name = sprintf('Surfaces_%s_%s.png', model_name, sugar_names{s});
        saveas(fig, fullfile(outFolder, fig_name));
        close(fig);
        
        fprintf('Figure saved: %s\n', fig_name);
        
    end  % End sugar loop
    
    fprintf('\n==> Excel file completed: %s\n', excel_file);
    
end  % End model type loop

fprintf('\n\n*** ANALYSIS COMPLETE ***\n');
fprintf('Output folder: %s\n', outFolder);


%% ================================================================
% HELPER FUNCTIONS
%% ================================================================

%% ---------------------------------------------------------------
% Build Scheffé design matrix with strain names as term names
%% ---------------------------------------------------------------

function [M, term_names] = build_scheffe_named(X, order, strain_names)
    q = size(X, 2);
    term_names = {};
    M = [];
    
    % Linear terms: AT, CP, T21, PC, F63, TH (6 terms)
    if order >= 1
        M = [M, X];
        for i = 1:q
            term_names{end+1} = strain_names{i}; %#ok<AGROW>
        end
    end
    
    % Quadratic terms: AT_CP, AT_T21, etc. (15 terms)
    if order >= 2
        for i = 1:q
            for j = i+1:q
                M(:, end+1) = X(:,i) .* X(:,j); %#ok<AGROW>
                term_names{end+1} = sprintf('%s_%s', strain_names{i}, strain_names{j}); %#ok<AGROW>
            end
        end
    end
    
    % Special cubic terms: AT_CP_T21, etc. (20 terms)
    if order >= 3
        for i = 1:q
            for j = i+1:q
                for k = j+1:q
                    M(:, end+1) = X(:,i) .* X(:,j) .* X(:,k); %#ok<AGROW>
                    term_names{end+1} = sprintf('%s_%s_%s', ...
                        strain_names{i}, strain_names{j}, strain_names{k}); %#ok<AGROW>
                end
            end
        end
    end
    
    % Full cubic terms: xi·xj·(xi-xj) noted as d_AT_CP (15 terms)
    % These capture asymmetric blending behavior
    if order >= 4
        for i = 1:q
            for j = i+1:q
                M(:, end+1) = X(:,i) .* X(:,j) .* (X(:,i) - X(:,j)); %#ok<AGROW>
                term_names{end+1} = sprintf('d_%s_%s', strain_names{i}, strain_names{j}); %#ok<AGROW>
            end
        end
    end
end

%% ---------------------------------------------------------------
% Compute ANOVA summary table
%% ---------------------------------------------------------------

function anova_table = compute_anova_summary(X, y, mdl, strain_names)
    n_obs = mdl.NumObservations;
    
    % Fitted values and residuals
    y_fitted = mdl.Fitted;
    residuals = mdl.Residuals.Raw;
    
    % SS Total (no intercept)
    SS_Total = sum(y.^2);
    DF_Total = n_obs;
    
    % SS Model
    SS_Model = sum(y_fitted.^2);
    DF_Model = mdl.NumCoefficients;
    MS_Model = SS_Model / DF_Model;
    
    % SS Residual
    SS_Residual = sum(residuals.^2);
    DF_Residual = n_obs - DF_Model;
    MS_Residual = SS_Residual / DF_Residual;
    
    F_Model = MS_Model / MS_Residual;
    p_Model = 1 - fcdf(F_Model, DF_Model, DF_Residual);
    
    % SS Linear (fit linear model separately)
    [M_lin, lin_terms] = build_scheffe_named(X, 1, strain_names);
    T_lin = array2table(M_lin, 'VariableNames', lin_terms);
    T_lin.Response = y;
    formula_lin = sprintf('Response ~ %s - 1', strjoin(lin_terms, ' + '));
    mdl_lin = fitlm(T_lin, formula_lin);
    
    SS_Linear = sum(mdl_lin.Fitted.^2);
    DF_Linear = mdl_lin.NumCoefficients;
    MS_Linear = SS_Linear / DF_Linear;
    F_Linear = MS_Linear / MS_Residual;
    p_Linear = 1 - fcdf(F_Linear, DF_Linear, DF_Residual);
    
    % SS Nonlinear
    SS_Nonlinear = max(0, SS_Model - SS_Linear);
    DF_Nonlinear = max(0, DF_Model - DF_Linear);
    
    if DF_Nonlinear > 0
        MS_Nonlinear = SS_Nonlinear / DF_Nonlinear;
        F_Nonlinear = MS_Nonlinear / MS_Residual;
        p_Nonlinear = 1 - fcdf(F_Nonlinear, DF_Nonlinear, DF_Residual);
    else
        MS_Nonlinear = 0;
        F_Nonlinear = NaN;
        p_Nonlinear = NaN;
    end
    
    % Lack of Fit / Pure Error
    [SS_LOF, SS_PE, DF_LOF, DF_PE] = compute_lack_of_fit(X, y, residuals, DF_Model);
    
    if DF_PE > 0
        MS_PE = SS_PE / DF_PE;
        MS_LOF = SS_LOF / DF_LOF;
        F_LOF = MS_LOF / MS_PE;
        p_LOF = 1 - fcdf(F_LOF, DF_LOF, DF_PE);
    else
        MS_PE = NaN;
        MS_LOF = SS_LOF / max(1, DF_LOF);
        F_LOF = NaN;
        p_LOF = NaN;
    end
    
    % Build table
    Row = {'Total'; 'Model'; '. Linear'; '. Nonlinear'; 'Residual'; '. Lack of fit'; '. Pure error'};
    SumSq = [SS_Total; SS_Model; SS_Linear; SS_Nonlinear; SS_Residual; SS_LOF; SS_PE];
    DF = [DF_Total; DF_Model; DF_Linear; DF_Nonlinear; DF_Residual; DF_LOF; DF_PE];
    MeanSq = [SS_Total/DF_Total; MS_Model; MS_Linear; MS_Nonlinear; MS_Residual; MS_LOF; MS_PE];
    F_val = [NaN; F_Model; F_Linear; F_Nonlinear; NaN; F_LOF; NaN];
    pValue = [NaN; p_Model; p_Linear; p_Nonlinear; NaN; p_LOF; NaN];
    
    anova_table = table(SumSq, DF, MeanSq, F_val, pValue, ...
        'RowNames', Row, ...
        'VariableNames', {'SumSq', 'DF', 'MeanSq', 'F', 'pValue'});
end

%% ---------------------------------------------------------------
% Compute Lack of Fit and Pure Error
%% ---------------------------------------------------------------

function [SS_LOF, SS_PE, DF_LOF, DF_PE] = compute_lack_of_fit(X, y, residuals, DF_Model)
    n = size(X, 1);
    
    % Round X to group replicates
    X_round = round(X, 6);
    [~, ~, group_idx] = unique(X_round, 'rows');
    n_groups = max(group_idx);
    
    % Pure Error (within-group variance)
    SS_PE = 0;
    DF_PE = 0;
    
    for g = 1:n_groups
        idx_g = (group_idx == g);
        n_g = sum(idx_g);
        
        if n_g > 1
            y_g = y(idx_g);
            y_mean_g = mean(y_g);
            SS_PE = SS_PE + sum((y_g - y_mean_g).^2);
            DF_PE = DF_PE + (n_g - 1);
        end
    end
    
    % Lack of Fit
    SS_Residual = sum(residuals.^2);
    SS_LOF = max(0, SS_Residual - SS_PE);
    
    DF_Residual = n - DF_Model;
    DF_LOF = max(0, DF_Residual - DF_PE);
end

%% ---------------------------------------------------------------
% Compute predictions for a triplet (for colormap normalization)
%% ---------------------------------------------------------------

function ypred_vals = compute_ternary_predictions(mdl, X, triplet, strain_names)
    % Fixed values for other components
    other = setdiff(1:6, triplet);
    fixed_vals = mean(X(:, other), 1);
    avail = 1 - sum(fixed_vals);
    
    % Grid
    gridN = 40;
    ypred_vals = [];
    
    coefNames = mdl.CoefficientNames;
    
    for i = 0:gridN
        for j = 0:gridN
            if i + j > gridN
                continue
            end
            k = gridN - i - j;
            u = i / gridN;
            v = j / gridN;
            w = k / gridN;
            
            % Real composition
            comp = zeros(1, 6);
            comp(triplet(1)) = u * avail;
            comp(triplet(2)) = v * avail;
            comp(triplet(3)) = w * avail;
            comp(other) = fixed_vals;
            
            % Predict
            y_pred = predict_from_composition(mdl, comp, strain_names, coefNames);
            ypred_vals = [ypred_vals; y_pred]; %#ok<AGROW>
        end
    end
end

%% ---------------------------------------------------------------
% Ternary surface plot (for tiledlayout)
%% ---------------------------------------------------------------

function plot_ternary_surface(mdl, X, triplet, strain_names, clim_range)
    % Triangle vertices
    A = [0, 0];
    B = [1, 0];
    C = [0.5, sqrt(3)/2];
    
    % Fixed values for other components
    other = setdiff(1:6, triplet);
    fixed_vals = mean(X(:, other), 1);
    avail = 1 - sum(fixed_vals);
    
    % Grid
    gridN = 40;
    xs = [];
    ys = [];
    ypred_vals = [];
    
    coefNames = mdl.CoefficientNames;
    
    for i = 0:gridN
        for j = 0:gridN
            if i + j > gridN
                continue
            end
            k = gridN - i - j;
            u = i / gridN;
            v = j / gridN;
            w = k / gridN;
            
            % Cartesian coordinates
            px = u*A(1) + v*B(1) + w*C(1);
            py = u*A(2) + v*B(2) + w*C(2);
            xs = [xs; px]; %#ok<AGROW>
            ys = [ys; py]; %#ok<AGROW>
            
            % Real composition
            comp = zeros(1, 6);
            comp(triplet(1)) = u * avail;
            comp(triplet(2)) = v * avail;
            comp(triplet(3)) = w * avail;
            comp(other) = fixed_vals;
            
            % Predict using coefficient names
            y_pred = predict_from_composition(mdl, comp, strain_names, coefNames);
            ypred_vals = [ypred_vals; y_pred]; %#ok<AGROW>
        end
    end
    
    % Interpolation
    F = scatteredInterpolant(xs, ys, ypred_vals, 'natural', 'none');
    xlin = linspace(0, 1, 150);
    ylin = linspace(0, C(2), 150);
    [Xgrid, Ygrid] = meshgrid(xlin, ylin);
    Zgrid = F(Xgrid, Ygrid);
    
    % Triangle mask
    mask = Ygrid <= sqrt(3) * min(Xgrid, 1 - Xgrid) + 1e-6;
    Zgrid(~mask) = NaN;
    
    % Plot
    surf(Xgrid, Ygrid, Zgrid, 'EdgeColor', 'none', 'FaceAlpha', 0.9);
    view(2);
    shading interp;
    colormap parula;
    
    % Apply normalized color limits
    if nargin >= 5 && ~isempty(clim_range)
        caxis(clim_range);  % Use caxis for MATLAB 2022b compatibility
    end
    colorbar;
    hold on;
    
    % Triangle outline
    plot([A(1), B(1), C(1), A(1)], [A(2), B(2), C(2), A(2)], 'k-', 'LineWidth', 1.2);
    
    % Experimental points
    Xtri = X(:, triplet);
    Xtri_sum = sum(Xtri, 2);
    Xtri_norm = Xtri ./ max(Xtri_sum, 1e-6);
    xpts = Xtri_norm(:,1)*A(1) + Xtri_norm(:,2)*B(1) + Xtri_norm(:,3)*C(1);
    ypts = Xtri_norm(:,1)*A(2) + Xtri_norm(:,2)*B(2) + Xtri_norm(:,3)*C(2);
    scatter(xpts, ypts, 15, 'k', 'filled');
    
    % Vertex labels
    text(A(1)-0.06, A(2)-0.04, strain_names{triplet(1)}, 'FontSize', 7, 'FontWeight', 'bold');
    text(B(1)+0.02, B(2)-0.04, strain_names{triplet(2)}, 'FontSize', 7, 'FontWeight', 'bold');
    text(C(1), C(2)+0.05, strain_names{triplet(3)}, 'FontSize', 7, 'FontWeight', 'bold', 'HorizontalAlignment', 'center');
    
    axis equal off;
    hold off;
end

%% ---------------------------------------------------------------
% Predict from composition using model coefficients
%% ---------------------------------------------------------------

function y_pred = predict_from_composition(mdl, comp, strain_names, coefNames)
    % Build all possible terms from composition
    coeffs = mdl.Coefficients.Estimate;
    
    y_pred = 0;
    
    for c = 1:length(coefNames)
        term_name = coefNames{c};
        
        % Check for full cubic terms: d_AT_CP means AT·CP·(AT-CP)
        if startsWith(term_name, 'd_')
            % Full cubic term: xi·xj·(xi-xj)
            rest = term_name(3:end);  % Remove 'd_'
            parts = strsplit(rest, '_');
            if length(parts) == 2
                idx1 = find(strcmp(strain_names, parts{1}));
                idx2 = find(strcmp(strain_names, parts{2}));
                if ~isempty(idx1) && ~isempty(idx2)
                    y_pred = y_pred + coeffs(c) * comp(idx1) * comp(idx2) * (comp(idx1) - comp(idx2));
                end
            end
        else
            % Parse term name to get indices
            parts = strsplit(term_name, '_');
            
            if length(parts) == 1
                % Linear term: AT, CP, etc.
                idx = find(strcmp(strain_names, parts{1}));
                if ~isempty(idx)
                    y_pred = y_pred + coeffs(c) * comp(idx);
                end
                
            elseif length(parts) == 2
                % Quadratic term: AT_CP, etc.
                idx1 = find(strcmp(strain_names, parts{1}));
                idx2 = find(strcmp(strain_names, parts{2}));
                if ~isempty(idx1) && ~isempty(idx2)
                    y_pred = y_pred + coeffs(c) * comp(idx1) * comp(idx2);
                end
                
            elseif length(parts) == 3
                % Cubic term: AT_CP_T21, etc.
                idx1 = find(strcmp(strain_names, parts{1}));
                idx2 = find(strcmp(strain_names, parts{2}));
                idx3 = find(strcmp(strain_names, parts{3}));
                if ~isempty(idx1) && ~isempty(idx2) && ~isempty(idx3)
                    y_pred = y_pred + coeffs(c) * comp(idx1) * comp(idx2) * comp(idx3);
                end
            end
        end
    end
end
```
