## Supplementary Data 3 - Codes for "Comparative Secretome Analysis and Enzyme Cocktail Optimization of Six Fungal Species Under Solid-State and Submerged Fermentation for Lignocellulosic Saccharification of Flax Shives": negEI_multi_for_fmincon.m

```
function nEI = negEI_multi_for_fmincon(x5, gpr_total, gpr_pgx, xi, w_total, w_pgx)
    % Combine deux GPR : total et pureté
    xsum = sum(x5);
    x6 = 1 - xsum;
    if any(x5 < 0) || x6 < 0
        nEI = 1e6;
        return
    end
    xfull = [x5(:)' x6];

    [mu_tot, s_tot] = predict(gpr_total, xfull);
    [mu_pgx, s_pgx] = predict(gpr_pgx, xfull);

    % Score combiné
    mu_comb = w_total*mu_tot + w_pgx*mu_pgx;
    s_comb  = sqrt((w_total*s_tot)^2 + (w_pgx*s_pgx)^2);

    bestY = max(w_total*gpr_total.Y + w_pgx*gpr_pgx.Y);

    if s_comb <= 0
        EI = max(0, mu_comb - bestY - xi);
    else
        z = (mu_comb - bestY - xi) / s_comb;
        EI = (mu_comb - bestY - xi) * normcdf(z) + s_comb * normpdf(z);
    end

    nEI = -EI;
end
```
